# The tiny germline chromosomes of *Paramecium aurelia* have an exceptionally high recombination rate and are capped by a new class of Helitrons

**DOI:** 10.1101/2025.11.06.686955

**Authors:** Olivier Arnaiz, Frédéric Guérin, Arnaud Couloux, Caridad Miró-Pina, Guillaume Pellerin, Irina Nekrasova, Joëlle Amselem, Jean-Marc Aury, Simran Bhullar, Andrea Frapporti, Emmanuelle Lerat, Isabelle Luyten, Sophie Malinsky, Nathalie Mathy, Alexey Potekhin, Vinciane Régnier, Natalia Sawka-Gądek, Amandine Touzeau, Augustin de Vanssay, Coralie Zangarelli, Hadi Quesneville, Mireille Bétermier, Karine Labadie, Laurent Duret, Eric Meyer, Sandra Duharcourt, Linda Sperling

## Abstract

**Background:** Paramecia belong to the ciliate phylum of unicellular eukaryotes characterized by nuclear dimorphism. A diploid germline micronucleus (MIC) transmits genetic information across sexual generations. A polyploid transcriptionally active somatic macronucleus (MAC) develops at each sexual generation from a copy of the MIC through programmed DNA elimination (PDE) of > 30% of germline DNA. PDE requires the domesticated PiggyMac (Pgm) transposase. Assembly of *Paramecium* germline genomes has presented an enormous challenge owing to the difficulty of MIC isolation.

**Results:** We report chromosome-scale short-read MIC assemblies for 7 species from the *P. aurelia* species complex. We discovered a novel clade of Helitrons, with 9-10 kb transposase ORFs under purifying selection, that have remained active in all *P. aurelia* lineages. A long-read assembly for *P. tetraurelia* together with a genetic linkage map provided a nearly telomere-to-telomere assembly.

**Conclusions.:** The genome consists of tiny (300 kb – 1.2 Mb) and numerous (∼160) germline chromosomes with the highest recombination rate ever reported for a eukaryote (420 cM/Mb). The ends of the chromosomes consist of Helitrons inserted in telomeric C_4_A_2_ repeats, forming a distinct genomic compartment that is eliminated very early during MAC development in a Pgm-independent manner.

## Background

The monophyletic Ciliophora, that emerged over a billion years ago [1], is comprised of highly diverse, very successful unicellular eukaryotes. Their most striking common feature is nuclear dimorphism, the separation of germline from somatic chromosomes in structurally and functionally distinct nuclei [2,3]. Diploid micronuclei (MIC) ensure transmission of the genome across sexual generations but are not expressed during vegetative growth while a polyploid macronucleus (MAC) is responsible for gene expression. At each sexual generation, a new MAC develops from a copy of the MIC through programmed DNA elimination (PDE) that removes transposable elements (TEs) and their remnants and restores functional genes [4–6].

The first ciliate genomes to be sequenced were the somatic MAC genomes of *Paramecium, Tetrahymena* and the distantly related spirotrich, *Oxytricha* [7–9], providing gene catalogues and information about MAC chromosome organization. The *P. tetraurelia* MAC genome assembly and annotations revealed a series of whole genome duplications (WGD) that occurred after the divergence of *Paramecium* and *Tetrahymena* (∼500 MYa). Partially resolved by gene loss over evolutionary time [7,10,11], the most recent WGD occurred concomitantly to the emergence of the *P. aurelia* complex of at least 16 morphologically identical but reproductively isolated sibling species [12–14]. While paramecia reproduce sexually by conjugation of two cells with compatible mating types, the individuals from *P. aurelia* species can also undergo a self-fertilization process, autogamy, that yields 100% homozygous progeny [15].

The germline MIC genomes of only a handful of model ciliates have been assembled and annotated. The degree of fragmentation of the assemblies varies depending on repeat content and the feasibility of purifying the MICs [16–20]. The amount of eliminated DNA, estimated by comparison of MIC and MAC assemblies, varies from ∼30% in *P. aurelia* and *Tetrahymena* to at least 95% in *Oxytricha* and *P. caudatum* [4,14]. So far, the only model ciliate for which we know the number and overall architecture of the germline chromosomes with confidence is *Tetrahymena thermophila* [18]. Centromeric regions are found at the center of 5 large, linear germline chromosomes and most TEs are restricted to the center and the ends of the chromosomes. The assembly confirmed that *Tetrahymena* uses a specific 15 nt Chromosome Breakage Sequence to fragment MIC chromosomes during MAC development [18]. In striking contrast, pioneer cytogenetic studies in *P. aurelia* species suggested karyotypes with tens of small MIC chromosomes (Dippell 1954; Jones 1956). Further, epigenetically determined fragmentation in *Paramecium* creates a variable set of linear chromosomes covering MAC-destined regions [6,21,22]. Analysis of the variability led to the suggestion that MAC scaffolds could represent the arms of MIC chromosomes [22]. This “metacentric chromosome” hypothesis has been difficult to test for lack of knowledge about *Paramecium* centromeres.

TEs can affect host viability by disrupting genes or altering their expression pattern [23]. A remarkable feature of *Paramecium* and other ciliates is the way they control TEs. Epigenetic mechanisms, involving meiosis-specific small RNA pathways that guide histone modifications, silence TEs in *Paramecium* as in most eukaryotes [24]. However, in *Paramecium* and other ciliates, the process is taken to the extreme of their physical elimination from the somatic MAC genome. In the germline MIC genome of *Paramecium*, exons are interrupted by single copy elements as small as 26 bp that are TE (Tc1/mariner DNA transposon) remnants known as Internal Eliminated Sequences (IES) [14,25]. Tens of thousands of IESs are precisely excised during MAC development, and time-course analysis of DNA elimination showed that IESs are eliminated earlier than TE or satellite sequences [16,26].

Elimination of IESs and most TEs and satellite repeats requires the domesticated PiggyMac (Pgm) transposase, which together with its Pgm-like partners, introduces DNA double strand breaks at the extremities of IESs [16,25,27,28]. This activity is tightly coupled to canonical non-homologous end-joining repair and gap filling of the cleavage sites [29–33]. Given the lack of a strictly conserved motif among eliminated sequences, how the Pgm excision complex is recruited to the boundaries of eliminated sequences is not fully understood. Data obtained so far indicate that all eliminated sequences do not rely on the same set of proteins for their elimination, suggesting that distinct yet partially overlapping pathways tether the Pgm excision machinery [34–36]. Indeed, most TEs and young, long IESs depend for their elimination on 25 nucleotide small RNAs, the scanRNAs (scnRNAs), whereas old, short IESs do not [14]. Initially produced from the entire MIC genome during meiosis by MIC-specific transcription factors [37,38] and a dedicated RNAi interference pathway [35,39–42], the scnRNA population corresponding to the non-eliminated sequences is subsequently degraded [41–44]. The remaining scnRNAs corresponding to TEs, in association with non-coding transcripts, then guide the Polycomb Repressive Complex 2 (PRC2) to catalyze histone modifications on TEs in the developing MAC, providing specificity despite the lack of a conserved sequence motif [34,36,45–47]. It is then believed that these repressive histone modifications recruit or activate the Pgm excision complex [24], which, assisted by chromatin remodelers [48–50], triggers DNA cleavage at the boundaries of the eliminated sequences.

Purification of vegetative *Paramecium* MICs by a fluorescence-activated cell sorting strategy [16] was a *tour de force* given the MAC ploidy of ∼1600n [51]: MIC DNA represents ∼ 0.25% of total nuclear DNA, compared to ∼ 5% in *Tetrahymena*. The first assembly using DNA from sorted MICs was fragmented (N50 ∼37 kb) but adequate for confirmation that all IESs are retained when Pgm endonuclease expression is knocked down [16]. The assembly allowed genome-wide discovery of TEs, revealing 38 non-LTR retrotransposon families and expanding knowledge of Tc1-mariner DNA transposons to 13 families. A fraction (∼3%) of the MIC assembly, enriched in satellite DNA but not in known TEs, was found to be eliminated independently of the Pgm endonuclease [16].

The present article describes the production of chromosome-scale assemblies of the MIC genomes of 7 species from the *aurelia* group. Our motivation was to understand chromosome organization in *P. aurelia*, determine the size and number of MIC chromosomes, identify and annotate repeated sequences and provide reference genomes for functional studies. Efforts to obtain a telomere-to-telomere assembly for the *P. tetraurelia* model, using long-read technology coupled with information from a genetic linkage map, revealed the highest recombination rate ever reported in a eukaryote and a diploid germline genome consisting of ∼160 pairs of small chromosomes. We discovered a new clade of Helitrons that have remained active in all lineages of the *aurelia* clade. Unlike previously annotated *Paramecium* TEs, most Helitrons are inserted in germline telomeres or sub-telomeric regions. Their elimination occurs earlier than that of other TEs and does not require the Pgm excision complex.

## RESULTS

### Assembly and annotation of Paramecium MIC genomes

#### MIC genome assemblies for 7 *P. aurelia* species

We selected species spread across the *Paramecium aurelia* clade (Figure 1A) for vegetative MIC purification by fluorescence-activated cell sorting and high-throughput sequencing using the Illumina platform, as previously described [14,16]. The strategy to build chromosome-scale MIC assemblies using these datasets is summarized in SupFigure1A. MIC DNA read-pairs were assembled into contigs. Then *PGM*-RNAi DNA isolated from cells depleted for the Pgm endonuclease by RNAi, available in greater quantities necessary to build mate-pair libraries, was used for scaffolding, as detailed in Methods.

**Figure 1.**
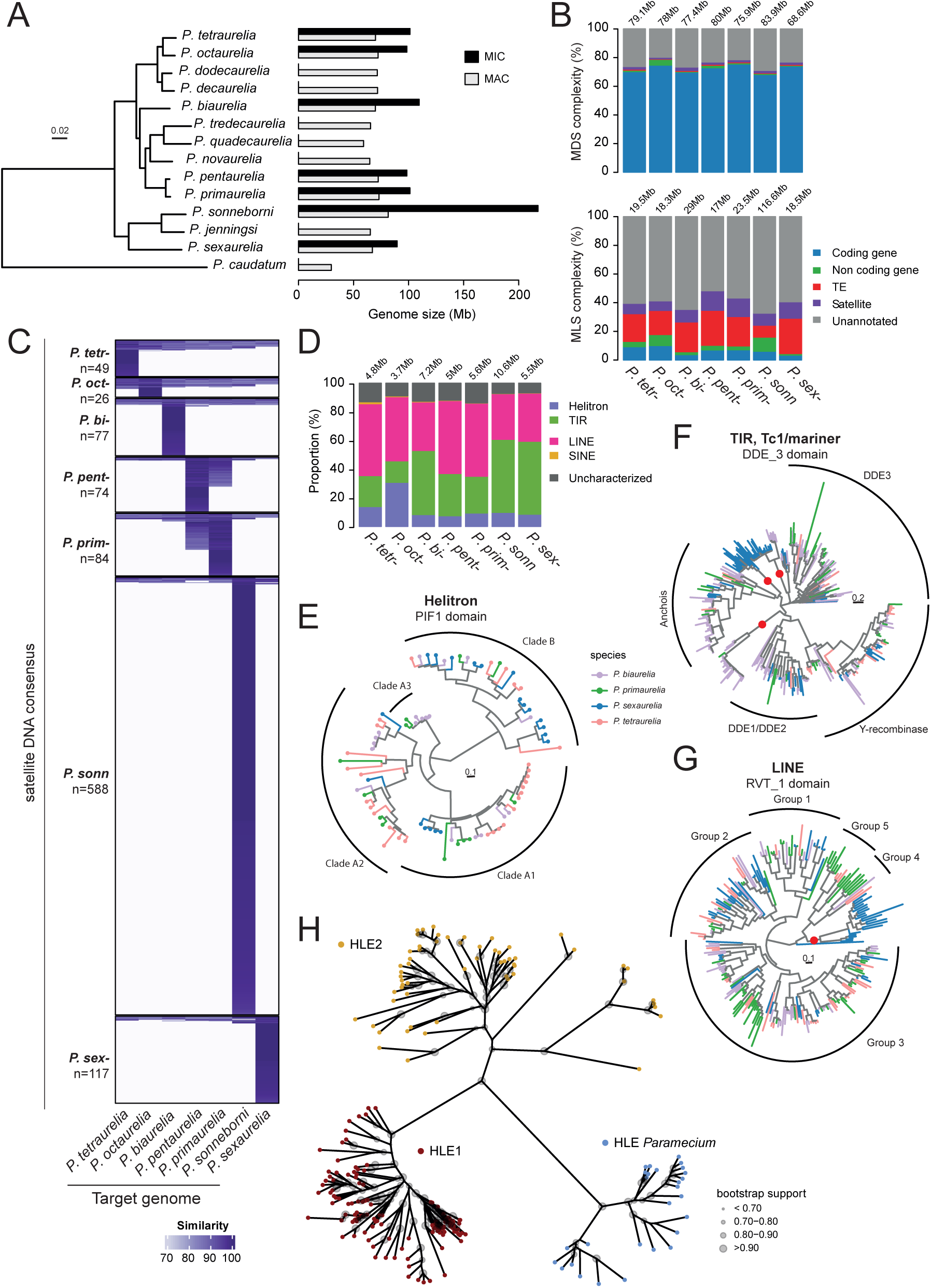
Paramecium aurelia MIC genome content. (A) Phylogenetic tree of P. aurelia species with P. caudatum as outgroup, modified from [14], showing the size of MIC and MAC assemblies. (B) Barplots representing genome occupancy of Coding and non-coding genes, Transposable Elements, and Satellites in the MAC-destined (MDS) and MIC-limited (MLS) regions of the P. aurelia MIC genome assemblies. Genome occupancy in Mb is given in SupFigure4. (C) Heatmap showing conservation of annotated minisatellites mapped to MIC genomes of each species using RepeatMasker (Methods). Each row corresponds to a minisatellite consensus and each column to a target species. The intensity of the blue rectangle is proportional to the similarity of the best hit to the target genome. Satellites are ordered by hierarchical clustering, showing the most conserved at the top. (D) TEs annotated using REPET were classified as Helitrons, TIRs, LINEs, SINEs and uncharacterized elements. Barplots show their MIC genome occupancy as percentage of total TE annotation. (E) Circular tree of Helitron PIF1 PFAM domains calculated with nucleotide alignments for 4 species. (F) PFAM DDE_3 domains were used to make a tree representing Tc1/mariner elements. Clades are labeled according to the classification described in [16]. Note recent acquisition of P. sexaurelia and P. biaurelia copies branching near the root of the DDE3, Anchois or DDE1/DDE2 clades, denoted by a red dot. (G) Circular tree of LINE RVT_1 PFAM domains, organized in 5 Groups according to the classification in [16]. Note recent acquisition of P. sexaurelia copies that branch before the separation of Group4 and Group5. (H) Phylogeny of concatenated Rep and Hel domains from across the tree of life [52] re-aligned with concatenated Paramecium Rep and Hel domains using MAFFT. The unrooted tree was constructed with phyML and drawn with ggtree. Bootstrap support is represented by the size of the circles drawn at nodes. The tree shows that a Paramecium HLE clade (blue tips) emerged very early and is distinct from the Helitron (HLE1) (red tips) and the Helentron/Helitron2 (HLE2) (yellow tips) clades.

The six scaffolded assemblies have an average MIC genome size around 100 Mb (Figure 1A), of which around 30% is eliminated during MAC development. With an average N50 of ∼500 kb, these assemblies are of good quality (SupTable1). Indeed, >99% of genes previously annotated using MAC genome assemblies [11,14,53] and >97% of previously annotated IESs [14,25] could be remapped to these assemblies. Analysis of the assemblies using the K-mer Analysis Toolkit [54] provided an assembly-agnostic assessment that the genomes are complete, with the possible exception of the *P. sexaurelia* assembly (SupFigure2).

The *P. sonneborni* genome is a case apart. Although scaffolds could not be built from the contigs for technical reasons (cf. Methods), the assembly (N50 ∼33 kb) seems to be complete (SupTable1; SupFigure2). The assembly size of 217 Mb is at least twice that of the other *P. aurelia* MIC genomes (Figure 1A), consistent with previously reported genome size estimates [14]. The large genome size is essentially owing to an increase of the MIC-specific compartment (the MAC genome size is ∼ 82 Mb compared to ∼70 Mb in the other *aurelia* species). Examination of the origins of the MIC sequences revealed that recurrent horizontal transfer of sequences from several other *aurelia* species can account for the increased genome size (see [55] ).

#### MIC genome content

*Paramecium* MIC genomes consist of MAC-destined sequences (MDS) colinear with the corresponding MAC genome and MIC-limited sequences (MLS) that are eliminated during MAC development. To discover sequence features, especially within the MLS compartment, we used dedicated software to annotate genes, tandem repeats (satellite DNA) and dispersed repeats (TEs) for each of the 7 MIC assemblies.

*Paramecium aurelia* MAC genomes are at least 70% coding harboring ∼40,000 genes of median size ∼1.2 kb with tiny introns, separated by very short intergenic regions [11,53,56,57] as shown in Figure 1B. Since examples of genes that contain MIC-specific sequences have been documented [14,40], we looked for genes completely embedded in *P. aurelia* MIC-limited regions (Methods). The high MDS gene density sharply contrasts with the low MLS gene density that was found. Indeed, as shown in Figure 1B, annotation of the assemblies detected a low density of 5 – 10% potential protein-coding genes (ATG … TGA, with or without introns), and an even lower density of potential non-coding genes (GC-content expected for coding regions but with no detectable ORF) in MLS. Moreover, the size of the ORFs in the MLS is significantly smaller than the size of ORFs in annotated MAC genes (SupFigure3A; median sizes ∼470 bp compared to ∼1.1 kb), as expected for genes that have decayed (pseudogenes). The same trend is found in a comparison of non-coding genes, though the difference is smaller (SupFigure3A). Using RNA-Seq data sets available for *P. tetraurelia* [53], expression of annotated MLS genes was examined (SupFigure3B). Except for a handful of genes (n = 29 with RPKM > 1, top of heatmap), none of the 3257 putative protein-coding genes or 1177 putative non-coding genes appear to be expressed, neither in vegetative cells nor during MAC development. This suggests that they are mainly annotation errors or pseudogenes. However in cells depleted for Ezl1, the catalytic subunit of Polycomb Repressive Complex 2 (PRC2) [45,46], the fraction of MLS genes that have transcripts (RPKM > 1 at T50) is larger (30%), showing they have the potential to be expressed.

To annotate satellite repeats, we combined two approaches, MREPS and TAREAN (Methods). Minisatellites, mainly localized in MLS, represent only ∼ 2% of the size of each *Paramecium* MIC assembly and ∼ 6% of the MLS (Figure 1B; SupFigure4A). Minisatellite repeat units are generally species-specific, apart from minisatellites common to *P. primaurelia* and *P. pentaurelia,* the most closely related species (Figure 1C, SupFigure5A). The large number of identified satellite repeats in *P. sonneborni* is very likely related to its large genome size. In order to see if the most abundant repeats are conserved across species, we sorted the units by abundance (SupFigure5B) and found that the most abundant units are not necessarily the most conserved. Since the diversification of the *aurelia* clade is relatively ancient (estimated 100 – 300 Mya) ([10,11], this is not surprising: rapid evolution of satellite repeats has been observed in other eukaryotes [58–61]. A notable exception is the 126 nt repeat (P126) similar to a WD40 repeat, the first *Paramecium* minisatellite identified because of its association with MAC telomeres [62].

Annotation of TEs (Methods) found elements from known superfamilies (LINE non-LTR retrotransposons and Tc1/mariner copy-and-paste DNA transposons) [16], and revealed Helitrons, DNA transposons that use a peel-and-paste mechanism of single-strand transposition [63,64]. SINEs, MITEs and uncharacterized repeats were also annotated. Despite the critical role of a domesticated piggyBac transposase in PDE, we found no traces of piggyBac transposons in any of the *P. aurelia* MIC genomes, in contrast with *T. thermophila* [18]. Surprisingly, although widespread in eukaryotic genomes, there was no hint of any LTR retrotransposons in these assemblies. LTR retrotransposons have not been found either in *Tetrahymena thermophila* [18] or the early-branching karyorelict ciliate *Loxedes magnus* [65], suggesting that these elements did not colonize ciliates.

The annotated TEs only account for ∼5 Mb of the total genome size (almost exclusively in MLS), less than 20% of the germline DNA eliminated during MAC development (Figure 1B; SupFigure4A).

Overall, TIR (Tc1/mariner) and LINE elements were the most abundant (∼ 2 - 3 Mb each) (Figure 1D; SupFigure4B). Helitron occupancy varied across the *aurelia* clade, on average ∼1 Mb (Figure 1D; SupFigure4B). In the *P. sexaurelia* /*P. sonneborni* sub-clade, LINEs appear to be less abundant than Tc1/mariner TEs. This difference could result from invasion or proliferation of elements after the divergence of the two *P. aurelia* sub-clades.

We also used a guided approach to annotate TE protein domains for the purpose of reconstituting the evolutionary dynamics of *Paramecium* Helitron, Tc1/mariner and LINE superfamily elements. We used Pfam domains specific for each superfamily to identify their occurrence in six-frame translations of the genome assemblies (Methods). The protein sequences of the least degenerate copies for Pif1 DNA helicase (Helitron), DDE_3 DDE endonuclease (TIR) and reverse transcriptase RVT_1 (LINE) domains were used to build phylogenies (Figure 1 E-G). The phylogenetic trees support ancient invasion of Helitron, Tc1/mariner, and LINE elements, pre-dating diversification of the *aurelia* clade.

Human curation of the Helitron transposase ORFs was carried out for the 7 *P. aurelia* species. Remarkably, compared to curated LINE and TIR transposase ORF copies [16], a greater proportion of the Helitron ORF copies appear to be full size (SupFigure6). Conceptual translations of 112 full size Helitron ORF copies were aligned and used to build a phyML tree that identified 29 elements (SupFigure7A and B) and has the same clade topology (A1, A2 and B) as the Pif1 domain tree (Figure 1E). The analysis of synonymous (dS) and non-synonymous (dN) substitution rates showed clear evidence of purifying selection, with an average dN/dS ratio over all branches of the phylogeny of 0.15±0.03 (see Methods).

A phylogeny of Helitron-like Elements (HLE) from across the tree of life was recently proposed using a new automated algorithm for their identification and classification [52]. Since Helitron transposases contain an HUH endonuclease domain (Rep) [63,66] followed by a Pif1-like helicase (Hel) domain [67], these authors made a phylogeny using Rep-Hel domains from across the eukaryotic tree and found only two clades, HLE1 and HLE2. We added the homologous domains from curated *Paramecium* Helitron transposase ORFs, aligned all the sequences and built an unrooted phyML tree (Figure 1H). The *Paramecium* HLEs are found in an independent clade (“HLE *Paramecium”)* that diverged from HLE1 and HLE2, with high bootstrap support (Methods).

Intriguingly, Helitron ORFs are interrupted by up to 4 IES insertions in 9 of the 29 identified elements (8 from clade A2 and 1 from clade B), a frequency close to that observed for cellular genes. These IESs are generally conserved in all annotated copies from all species. One remarkable case is an IES inserted at the same site, in a conserved domain of the transposase, in all copies of 5 related elements forming a well-defined A2 subclade represented in all species (SupFigure7B). This IES insertion therefore likely became fixed in a common ancestor of the 5 elements before the beginning of speciation in the *P. aurelia* complex. In contrast to the flanking coding sequences, the IES appears to have evolved with almost no constraint, as previously noted for IESs inserted within cellular genes (SupFigure7C). Excision timing is similar to that of other IESs (SupFigure7D).

Even with improved annotation tools compared to the approach used previously for P. tetraurelia [16], a large portion of Paramecium MIC-limited genomes (50% - 70%) does not correspond to any annotated feature (gene, TE or satellite) (Figure 1B). It remains very difficult to know whether regions with no annotation correspond to “dark matter” i.e. sequence features no longer under selection that have evolved beyond recognition [68], or sequence features we do not yet know how to annotate.

### Counting chromosomes

It has long been suspected that *Paramecium aurelia* species harbor a large number of small germline chromosomes, based on very early cytogenetics [69,70]. We report in this section the use of long-read sequencing to achieve a distinct, more complete assembly for *P. tetraurelia* as well as the construction of a genetic linkage map, leading to prediction of the karyotype of *P. tetraurelia*.

Pgm depletion does not block elimination of ∼3 Mb of MIC-specific genome sequence, enriched in satellite DNA [16]. The short-read assemblies scaffolded with *PGM* RNAi DNA libraries are thus likely to be sub-optimal, especially for highly repeated regions of the genome. To try to improve closure and obtain telomere-to-telomere chromosomes, we used the *Oxford Nanopore Technology* (ONT) to generate long reads. Since more DNA than was available from sorted vegetative MICs was required to build the sequencing libraries, we used DNA from new MACs of *P. tetraurelia* cells depleted in Ezl1, the catalytic subunit of Polycomb Repressive Complex 2 (PRC2) required for PDE [34,45,46], to prepare enough unrearranged DNA for sequencing. In *EZL1*-RNAi DNA, 3 Mb of additional MLS are retained compared to *PGM* RNAi DNA [16], however ∼33% of IESs (0.73 Mb; SupTable1) are excised [34]. With long reads of this *EZL1*-RNAi DNA (1M reads with a median size ∼14 kb), a high quality *de novo* assembly was achieved (N50 ∼615 kb, 202 contigs, assembly size ∼104 Mb; Methods, SupFigure1B, SupFigure2, SupTable1). This assembly is less fragmented than the short-read assembly, contains all genes and the expected number of IESs (67%) and provides an improved reference to study the organization of the *P. tetraurelia* MIC genome.

#### Construction of a genetic linkage map

To help complete the assembly we constructed a genetic linkage map. We sequenced (*Illumina* platform) and analyzed total DNA (essentially MAC DNA owing to high MAC ploidy) from 39 F2 clones of a genetic cross between two *P. tetraurelia* strains, 51 and 32 (Methods; SupFigure8A). We retained 220,994 biallelic SNP markers for genotyping (SNP density of 0.3% in MAC-destined regions of the genome). We detected a total of 12,206 cross over events (CO) in 39 meioses (313 COs per F2) (Figure 2A&B; SupFigure8A&D). This corresponds to a recombination rate of 420 cM/Mb in the MAC-destined compartment, a value that is extremely high compared to that found for most of the other species studied to date (SupFigure8E), and a genetic length of 31,000 cM, by far the highest ever reported (Figure 2C).

**Figure 2.**
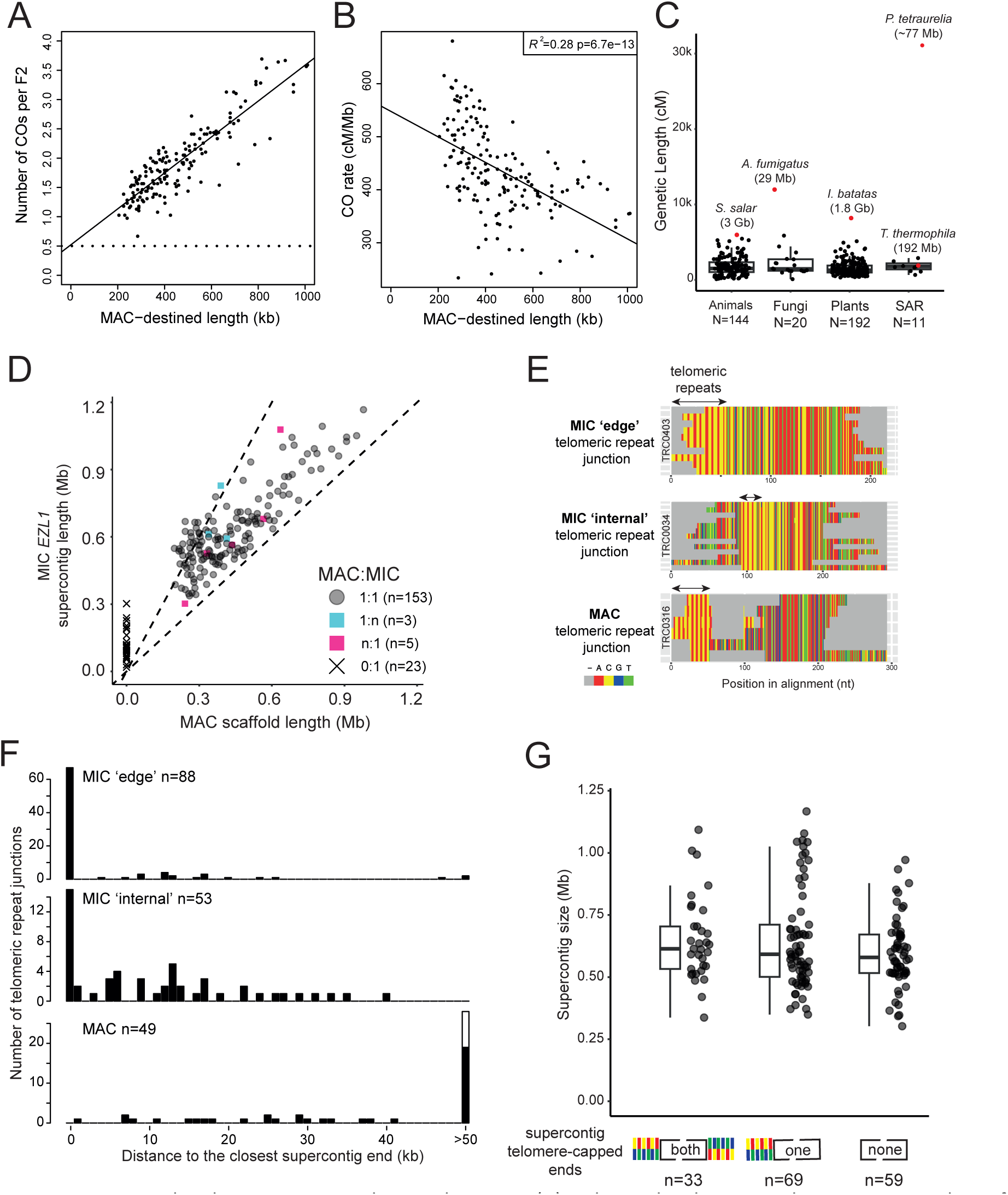
P. tetraurelia chromosomes and recombination. (A) Relationship between the average number of crossovers per F2 and the length of MAC-destined regions. (cf. Methods and SupFigure8). The 161 points correspond to 158 supercontigs of the MIC genome assembly, and to 3 groups of 2 supercontigs that are linked based on the MAC assembly. The horizontal dotted line at 0.5 is equivalent to 50 cM. (B) Relationship between the recombination rate (cM/Mb) and the length of MAC-destined regions (N=161). (C) Distribution of genetic map lengths (cM) across eukaryotes. (D) Correspondence of MIC supercontig length (n=187) with the length of MAC scaffolds (MAC-constitutive scaffolds, MAC v2.0; n = 162). Each dot gives the (sum of the) length(s) of the MIC supercontig(s) as a function of the (sum of the) length(s) of the corresponding MAC scaffold(s). The dotted lines show that MIC supercontigs are up to twice as large as the corresponding MAC scaffolds. In three cases, two supercontigs encompassed contiguous segments of the same MAC scaffold (category 1:n). In 5 cases, one MIC supercontig corresponded to 2 MAC scaffolds (category n:1). See SupData1_MIC_MAC for details. (E) Illumina MIC reads with telomere repeats (mix of C4A2 and C3A3 hexamers) were clustered and aligned (Methods; SupData2_TelomereJunctions). Three categories of junctions were found: MIC ‘edge’ telomeric repeat junctions, MIC ‘internal’ telomeric repeat junctions. The last category, MAC telomeric repeat junctions are heterogeneous among the sequenced reads, the expectation for de novo telomeric repeats added to DNA extremities generated during PDE [21]. (F) Distance between telomere repeat junctions and the nearest supercontig end, for each category. The last bin of the barplots represents distances of 50 kb or greater. MIC ‘edge’ junctions are mostly at supercontig ends (n = 88): 66 (75%) map at a distance of 0 from the supercontig end as expected for true MIC telomeres in the unrearranged MIC genome. The others (n= 22, 25%) may represent MIC internal telomeric junctions. The telomere repeat clusters classified as MIC internal telomeric junctions (n=53) are mostly (n=43; 81%) less than 20 kb from supercontig ends. MAC telomeric repeat junctions are further from supercontig ends (9 / 49, 18% at <20kb). Note that telomeric repeats can be found in either of two orientations: 5’-CA or 5’-GT. Only MAC telomeres can be detected in both orientations (black and white bars). (G) Lengths of supercontigs containing MLS that are capped by telomere repeats at both ends, one end or neither end.

In most eukaryotes, CO interference [71] constrains spacing of otherwise randomly located CO sites along chromosomes. We measured a 50% CO deficit at a distance of 22 kb comparable to that reported in *S. cerevisiae* (30 kb) (SupFigure8C) [72]. We also found evidence of aneuploidy (Methods) since 8 F2s (21%) carried some scaffolds with a doubled ploidy (5 with only one scaffold, the others with respectively 3, 4 and 13 aneuploid scaffolds) (SupFigure9). Similarly, using 74 independent published DNA-Seq datasets prepared from 100% homozygous cell lines, we found 21 additional cases of doubled ploidy for 1 or more scaffolds (28%) (SupFigure9). These cases of aneuploidy most probably result from chromosome segregation errors during meiosis and they occur with a frequency comparable to that reported in human (23.6%; [73]) (SupFigure9; cf. Discussion).

Analysis of linkage revealed that 37 *P. tetraurelia* contigs could be clustered into 17 linkage groups, and that a few contigs consisted of two genetically unlinked segments, suggestive of chimeric assembly. We therefore built a composite genome, called the “MIC *EZL1* assembly”, using the linkage map to split or join the contigs into 187 supercontigs (N50 ∼620 kb, 104.3 Mb). This assembly includes 164 supercontigs (101.4 Mb in total) that encompass 166 MAC scaffolds, and 23 supercontigs (2.9 Mb in total) consisting only of MLS, for which it was not possible to analyze genetic linkage. There is an essentially 1:1 relationship between the MIC *EZL1* supercontigs and the MAC scaffolds (Figure 2D; SupData1_Synteny_MIC_MAC.html; SupFigure10). Often, the MIC *EZL1* supercontig is at least twice as large as the MAC scaffold, corresponding to significant DNA elimination. More detailed analysis of the synteny between MIC *EZL1* and MAC regions (Methods) showed that the sequences present in MIC *EZL1* that are absent from the MAC scaffolds are found primarily at supercontig ends but sometimes are more centrally located (cf. annotated Circos drawings: SupData1_ Synteny_MIC_MAC.html). We conclude that the fragmentation sites used for MAC development are situated at the extremities of MIC chromosomes, although additional sites are sometimes more internal.

### MIC telomeres

Have we achieved a telomere-to-telomere MIC genome assembly? Are MIC telomere repeats the same as MAC telomere repeats? To answer these questions, we first sought to identify MIC telomere repeats using the Illumina read data. In eukaryotes, telomeres almost always consist of tandem repeats of a 6, 7 or 8 nt motif, usually GT-rich in the strand oriented 5’ to 3’ towards the end of the chromosome. The well-characterized *Paramecium* MAC telomere repeats that assemble *de novo* to heal programmed and accidental double-strand breaks consist of random mixtures of TTGGGG and TTTGGG hexamers [74], the result of misincorporation at a single templating position in the telomerase RNA [75,76].

Assuming that *P. aurelia* MIC telomeres are also tandem repeats of a short motif, we looked for candidates using a *sans a priori* analysis of the *k*-mers in all read pairs that did not map properly to the MAC assembly (Methods). We found the most abundant G-rich motif was TTGGGG (equivalent to 5’CCCCAA3’ repeats at the beginning of the scaffold), suggesting that MIC and MAC telomere repeats are the same. The repeats are located at or near scaffold or supercontig ends. For *P. tetraurelia,* the repeats were not only restricted to contig ends but were much more abundant in the MIC *EZL1* long-read assembly than in the short-read assembly (SupFigure11A), suggesting that MIC telomere repeats were probably collapsed or on small contigs not incorporated into the short-read assembly.

We used an orthogonal approach, independent of the assemblies, to gain insight on telomere branch points. We selected *Illumina* MIC sequencing reads containing a junction between telomeric repeats and non-telomeric sequences, clustered them and used multiple alignments to obtain consensus sequences (Methods, SupData2). Although this approach is limited by sequencing depth and cannot provide an exhaustive catalogue, it is possible to identify the junctions between MIC telomeric repeats and flanking sequences and to distinguish them from MAC junctions. Indeed, in the case of MAC telomeres, telomere addition does not always occur at the same position among all reads because chromosome fragmentation is an imprecise process that occurs in the new MAC after several rounds of endoreplication [21]. In the case of MIC telomeres present in the unrearranged germline genome, telomere branch points are expected at exactly the same position in all reads.

Surprisingly, we found two categories among the sampled MIC junctions. In the first category, telomeric repeats are flanked on only one side by non-telomeric sequences (“edge” junctions), as must be the case for the telomeres of MIC chromosomes. In the second category, the blocks of telomeric repeats are flanked on both sides with non-telomeric sequences (“internal” junctions), corresponding to telomeric repeat blocks within MIC chromosomes. It is important to note that a MIC “edge” junction does not guarantee that the telomeric repeats are at the end of a chromosome, because the clusters are short local alignments (180 – 400 bp, mean 252 bp). Examples of MAC, MIC “edge”, and MIC “internal” telomeric junctions are shown in Figure 2E. We were able to map 87% of these telomere junction consensus sequences to the *P. tetraurelia* MIC *EZL1* genome assembly (Methods; SupData2_TelomereJunctions). After removing redundant junctions that correspond to the same loci, we retained 88 MIC “edge”, 53 MIC “internal” and 49 MAC junctions for further analysis. To see whether these telomeric repeat junctions are at chromosome ends, we plotted their distance from the closest supercontig end (Figure 2F). As expected, MAC telomere junctions do not map at supercontig ends, a majority of 57% are at a distance of at least 50 kb (28/49) and are found in both orientations (GT or CA repeats near the 5’ end of supercontigs). The MIC junction mapping provides a totally different picture. We found that 75% (66/88) of the MIC “edge” junctions map at the very end of supercontigs (distance = 0) in the expected orientation (CA strand of telomeric repeats on the 5’-end strand) and may thus represent the telomeres of MIC chromosomes. The “internal” junctions are also close to the supercontig ends and, intriguingly, are in the same orientation. There are 8 cases (15%) where internal junctions do map to the very end of a supercontig, but this corresponds to situations where a few long-reads suggest that the supercontig could be extended to the end of the chromosome.

These results indicate that MAC telomere repeat junctions are farther from supercontig ends than MIC telomere repeat junctions. Since none of the sampled MAC junctions was closer than 1.5 kb from a supercontig end, correctly oriented telomeric repeats that cap supercontigs are most probably MIC telomeres. Considering only the 161 supercontig groups that are syntenic with MAC scaffolds (Figure 2D and SupData1), we analyzed the size distributions of supercontigs capped by telomere repeats at both ends (n = 33), one end (n = 69) or neither end (n= 59) (Figure 2G). The 33 supercontigs capped at both ends are likely complete MIC chromosomes. We note that the size distribution of supercontigs that are missing telomere repeats does not differ significantly from the size distribution of supercontigs that are capped at both ends (p > 0.09, t-tests, Figure 2G). This suggests that they all could be quasi-complete and hence that *P. tetraurelia* has as many as 161 MIC chromosomes.

### Chromosome organization

#### Chromosome ends do not require Pgm for elimination

The *Paramecium* germline genome is organized in distinct regions according to developmental fate. Constitutively MAC-destined regions are always retained in the new MAC and variable MAC-destined regions are partially retained via alternative elimination [14]. MIC-limited regions are eliminated during MAC development. They include IES and OES (Other Eliminated Sequences) [77]. The question addressed in this section, is where these regions are located on the chromosomes.

Genomic compartments were operationally defined using short read coverage of DNA extracted from sorted nuclei: MICs, MACs, and developing MACs after *PGM-*RNAi (SupFigure12; Methods). This is illustrated for one *P. tetraurelia* supercontig in Figure 3A and drawings of all supercontigs are provided in SupData3 & 4. Constitutively MAC-destined regions represent 70.2 Mb of the assembly, variably MAC-destined regions 9.2 Mb, and the MIC-limited compartment 19.9 Mb (IES 3.9 Mb, OES 16 Mb). The OES is further divided into a Pgm-dependent compartment (13.3 Mb) maintained in new MACs after Pgm depletion and a Pgm-independent compartment (2.7 Mb) first noticed by Guérin et al. 2017 [16], that is eliminated even upon Pgm depletion. Although considered to be a distinct compartment, all IESs are Pgm-dependent [16,25].

**Figure 3.**
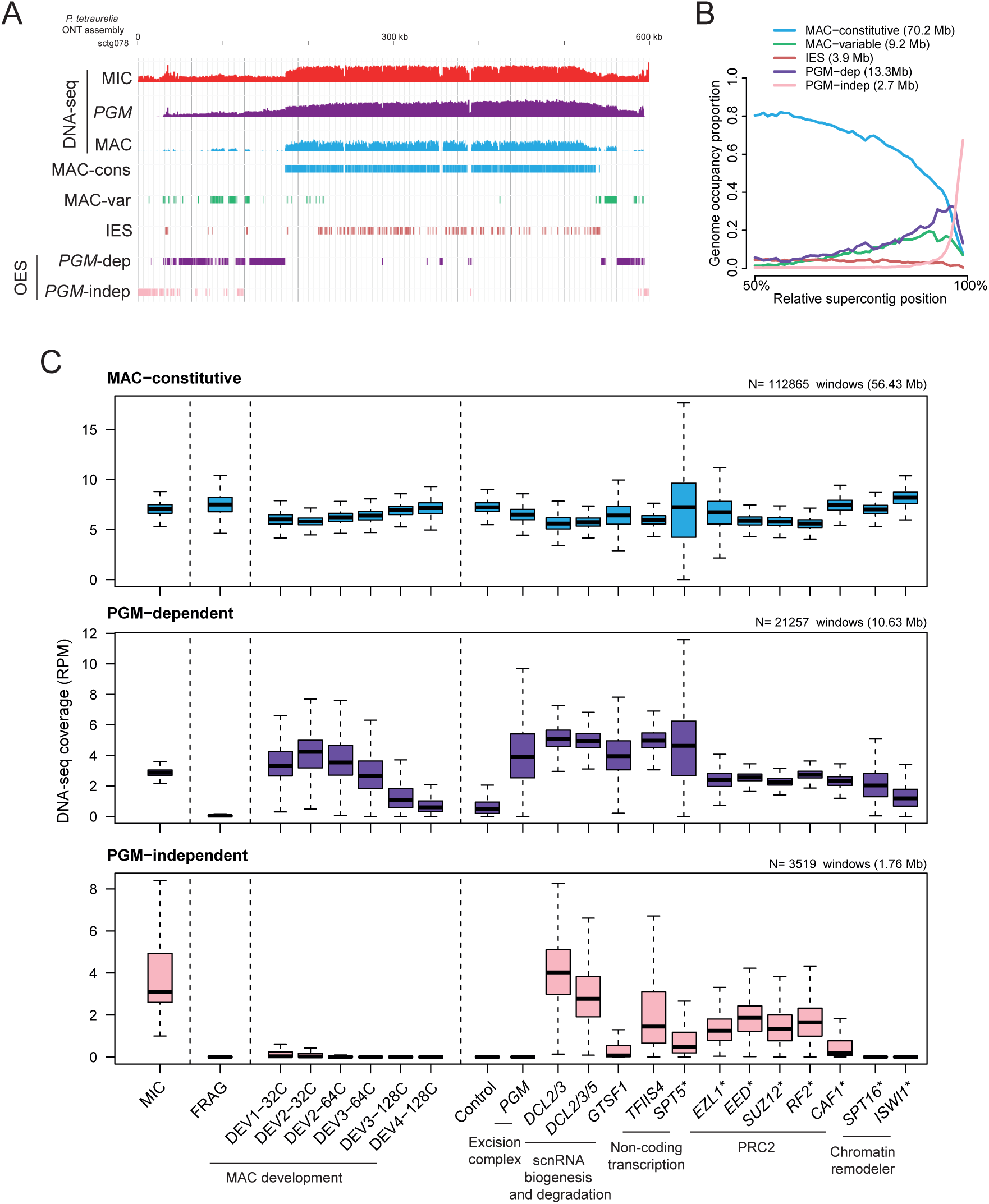
Definition and characteristics of genomic compartments. (A) Genomic compartments defined by sequencing read coverage. JBrowse screen shot shows DNA-Seq coverage of a region of the MIC EZL1 assembly by MIC, PGM-RNAi and MAC Illumina sequence reads (top three tracks). MAC-constitutive and MAC-variable compartments are determined by a high and low MAC read coverage, respectively. IESs were annotated using ParTIES [78]. The OES (Other Eliminated Sequence) regions were defined by no MAC coverage (IESs excluded) then divided into two compartments: PGM-dependent and PGM-independent, based on the PGM RNAi DNA coverage. See Methods and SupFigure12 for details. (B) Density of each genome compartment across assembly supercontigs showing their relative localization on folded chromosomes: 50% corresponds to the center, 100% the ends. (C) Normalized DNA coverage (RPM) for non-overlapping 500 nt windows. The windows were separated into 3 groups, as a function of their compartment (MAC-constitutive, Pgm-dependent, or Pgm- independent). The boxplots summarize the presence of DNA of each compartment, for different DNA-Seq samples: sorted vegetative micronuclei (MIC), sorted parental old MAC fragments (FRAG), sorted developing MAC time course (DEV1 - DEV4) at different ploidy levels (32C to 128C) [26], and sorted or enriched (designated by a “*”) new MACs after gene depletion by RNAi; the samples are grouped by pathway (excision complex, scnRNA biogenesis and degradation, non-coding transcription, PRC2 components, and chromatin remodelers). These DNA-seq datasets are described in SupTable2.

Figure 3B shows the relative position of each compartment calculated across all the supercontigs of the assembly. As in the example (Figure 3A), the relative position of the MAC-constitutive compartment and of the IESs is greatest at the center of the supercontigs and vanishes towards the ends, while the PGM-independent compartment is restricted to the ends. MAC-variable and PGM- dependent compartment occupancy curves peak towards the chromosome ends but then decrease sharply.

Time-course analysis of DNA elimination indicates that the Pgm-independent compartment is eliminated very early during MAC development, before IESs are excised and before the ploidy in the new MAC reaches 32C (Figure 3C) [26]. To gain insight into the mechanisms of elimination of this compartment, we examined the consequences of depletion of 32 known proteins involved in PDE, using DNA-Seq of new MACs (see Methods) (Figure 3C, SupFigure12). As expected, the Pgm excision complex (Pgm, PgmL and Ku70/80c proteins) is dispensable for the elimination of this compartment. This is also the case for the chromatin remodelers ISWI1 [49] and Spt16 [48], which are required for IES excision. Given that the assembly was constructed using long reads from DNA of cells depleted for Ezl1, the catalytic subunit of PRC2, the Pgm-independent compartment is not eliminated in this condition (Figure 3C). Unsurprisingly, depletion of other PRC2 components or cofactors (Eed, Suz12, Caf1, Rf2) also results in failure to eliminate this compartment. We note that two PRC2 cofactors, Eap1 and Rf4 [46,47], appear to have a weaker effect compared to the other PRC2 components (SupFigure12). Depletion of the proteins required for scnRNA biogenesis (Spt5 [37]; Dcl2/3 [34,35]), degradation of MIC-limited scnRNAs (Gtsf1 [43,44]), and non-coding transcription (TfIIs4, [36]) showed they are important for its elimination as well (Figure 3C, SupFigure12). Consistent with previous reports demonstrating that scnRNAs are produced from the entire MIC genome during meiosis [40,42], the PGM-independent compartment is covered with scnRNAs (SupFigure13). We conclude that the elimination of the Pgm-independent compartment occurs very early during MAC development and involves the scnRNA and PRC2 pathways.

The next question is the localization of annotated sequence features with respect to the genomic compartments. A majority of TEs (75.3%) are in the Pgm-dependent compartment (Figure 4A). More specifically, this is the case for TIR (Tc1/mariner) and LINE copies, while a majority of Helitrons are found in the Pgm-independent compartment. Consistent with the position of the different genomic compartments (Figure 3A), the Helitrons are overwhelmingly localized towards supercontig ends while LINEs and TIRs have a widespread distribution (Figure 4B). Minisatellites are also widely distributed across all compartments, as illustrated by P126. However, some abundant minisatellites (CHBD0, SAT149, SAT118, SAT189 and SAT51) are exclusively present (>90%) in the Pgm- independent compartment. As shown in Figure 4C, SAT149 and SAT51 are located at the very end of supercontigs, while CHBD0, SAT118 and SAT189 occur near the extremity.

**Figure 4.**
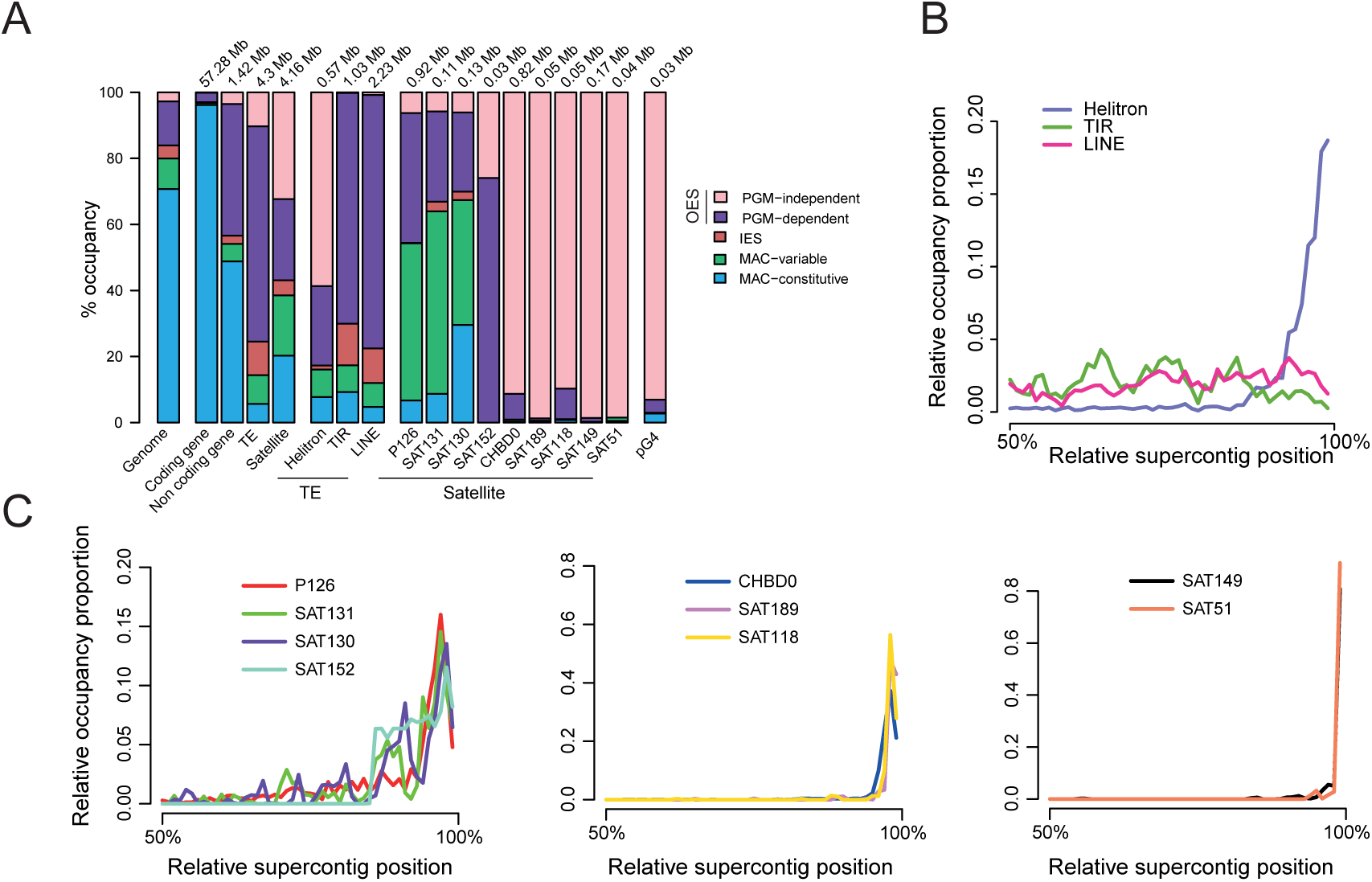
Genomic compartments and annotation landscape. (A) Barplots show the proportion of each annotated sequence feature (genes, TE and satellites) occupied by the 5 P. tetraurelia genomic compartments of the MIC EZL1 assembly. The compartment occupancy proportion for each TE order (Helitron, TIR, LINE) and of the most abundant satellite families is also presented. The last bar shows the compartment occupancy of regions (n=977) that have the potential to form G-quadruplex (pG4) and do not overlap telomeric repeats known to potentially form G4 (see Methods; SupFigure12). (B) Relative supercontig position of Helitron, TIR and LINE occupancy on MIC supercontigs (as in Figure 3B, the abscissa presents normalized distance to the nearest supercontig end); Helitrons are restricted to the ends of the supercontigs, while TIRs and LINEs are found everywhere. (C) Localization of the most abundant minisatellites on MIC supercontings.

### Helitron insertions are restricted to chromosome ends

Helitrons are mainly localized in the Pgm-independent compartment (Figure 4A&B). We looked more precisely at excision timing, excision requirements and expression for each of the different Helitron copies with 9-10 kb ORFS (n=79). The heatmap (Figure 5A), showing sequencing coverage in different contexts, confirms that these elements are present in the MIC, absent from the MAC, retained upon PRC2 depletion, but for the most part are still eliminated upon Pgm excision complex depletion. As shown for the entire Pgm-independent compartment (Figure3C), the elimination of the majority of Helitron ORFs is sensitive to depletion of factors involved in non-coding transcription and the scnRNA pathway (Figure 5A, SupFigure14). Helitron ORFs that belong to Clade A1 (Hel01, Hel03, Hel04, and Hel18) are eliminated early in new MAC development (before 32C endoreplication stage [26]). The other Helitrons are eliminated later, but earlier than LINEs and TIRs which are eliminated very late in MAC development (from 128C). Interestingly, some Clade A2 Helitrons (Hel09, Hel19), those that contain IESs and are retained after Pgm depletion, are also eliminated very late. Finally, some Helitron ORFs can be transcribed (Figure 5A last column of heatmap) as observed when Ezl1 or other PRC2 components are depleted (SupFigure14).

**Figure 5.**
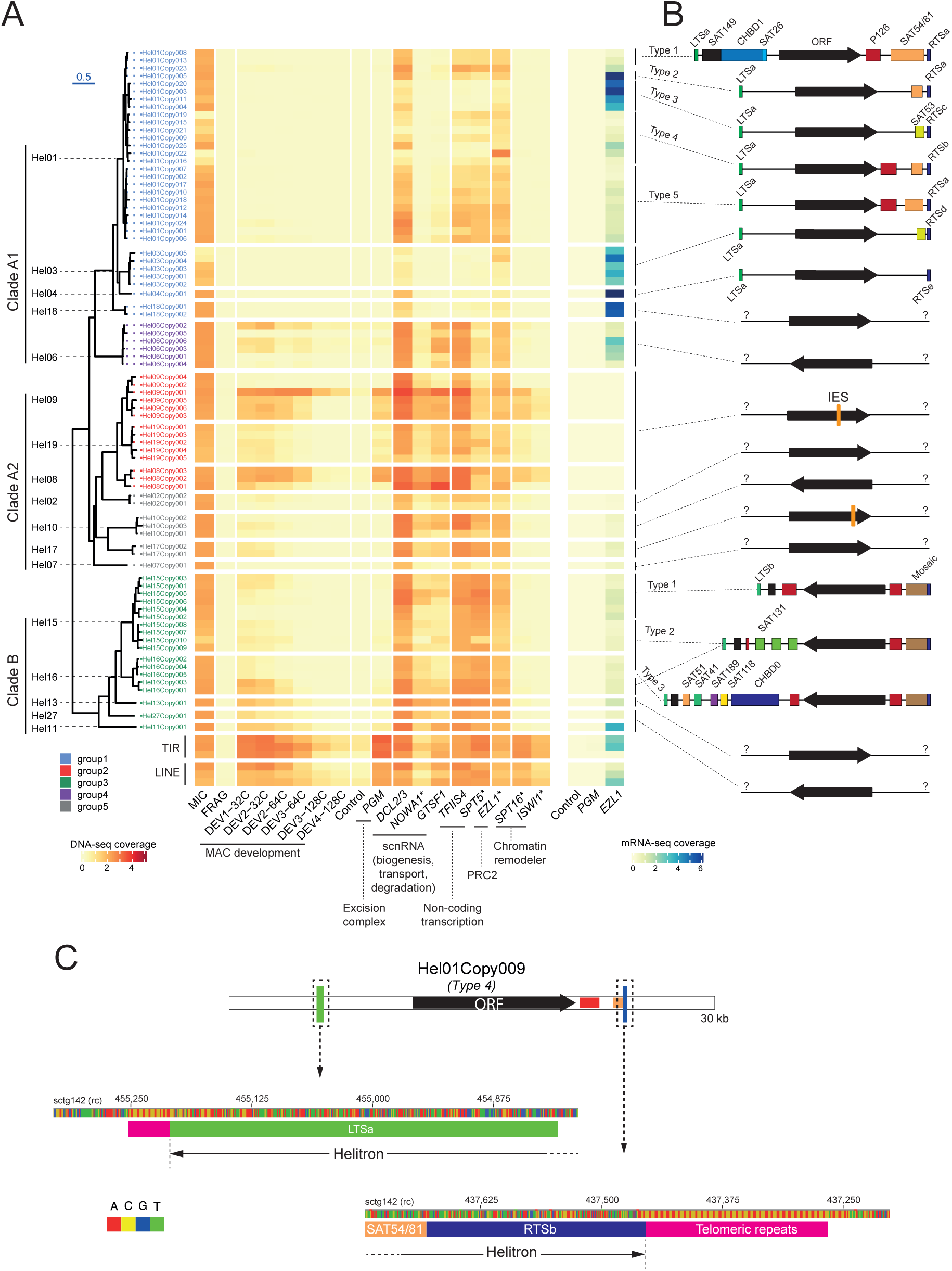
Paramecium Helitron classification and characterization. (A) Phylogenetic tree of Helitron transposase copies found in the P. tetraurelia MIC EZL1 reference assembly. The phyML amino acid tree was made using MACSE to align the last 5000 nt of 79 full-length Helitron transposase ORFs, containing the conserved HUH Y2 (Rep) and PIF1 (Hel) domains (see SupFigure7A). We used this tree to define 5 Helitron Groups. Clade A2 is split into 2 groups based on the presence of a conserved IES in Hel08, Hel09, and Hel19 copies (Group 2, red labels; cf SupFigure7) but not in the other Helitrons in the clade (Group 5, gray labels). The heatmap was made using DNA-Seq and RNA-Seq RPKM coverage after normalizing copies with respect to collapse (Methods). The 3 longest TIR and LINE copies were also added to the heatmap for comparison. Samples used to calculate coverage are described in SupTable2. DNA samples from enriched but not sorted nuclei are designated by “*”. (B) Structure of some Helitron elements showing the position and orientation of the transposase ORF with respect to the supercontig end (on the left). Helitron Left Terminal Sequence (LTS) (green boxes) and Right Terminal Sequence (blue boxes) were identified by manual alignment and curation. The satellites found between the LTS and the RTS are indicated by colored boxes. For the most abundant Helitrons (Hel01/15/16) the combination of LTS-satellites-RTS defines different “Types”. The presence of an IES is indicated for Hel09/19/08/17. The schema is not to scale. See SupFigure16 for more details. (C). Schematic representation of the Hel01Copy009 copy. ORF, LTS, RTS, and minisatellites are represented by colored boxes (green: LTSc, black:ORF, red: P126, orange: SAT54/81, blue: RTSb). In the zoom to the DNA sequence level before LTS and after RTS, flanking telomeric repeats (pink boxes) are found at both extremities.

### Structure of complete *Paramecium* Helitrons

Helitron transposition involves nicking of the top strand at the 5’ end of a Left Terminal Sequence (LTS) which forms a covalent bond with the transposase, unwinding of the double-helix, and introduction of a second nick at the end of the Right Terminal Sequence (RTS), shortly after a stem-loop structure, to form a circular intermediate [64,79].

To determine the full extent of transposable units, we looked for conserved terminal sequences marking the boundary with copy-specific flanks. We used DNA dotplots to compare ∼50 kb genomic regions centered on the ORFs, for all copies of the abundant elements Hel01 (Figure 5A clade A1) and Hel15/Hel16 (Figure 5A clade B). Example dotplots between pairs of different copies of Hel01 are shown in SupFigure15. The ORF sequences present a continuous diagonal, surrounded by a variety of satellites that often differ in length among closely related copies (rectangles along diagonal) and are altogether missing in more distant copies (gaps in the diagonal). Alignment of the last 350 nt before the sequences diverge revealed 3 related consensus sequences, RTSa, RTSb, and RTSc, with a short stem-loop and ending with CCAA (SupFigure17A), an observation reminiscent of the CTRR found at the end of RTS in other eukaryotes [52,80].

Similarly, RTSd and RTSe were found, by homology to RTSa-c, to end Hel03 and Hel04 (Figure 5A&B). Alignment of RTSa-d consensus sequences and the unique RTSe showed characteristic RTS stem-loops with compensatory base changes, in support of this structure (SupFigure17A). The left end of the 10 Hel01 copies that did not extend beyond the end of the assembled supercontig (all Hel01 copies are oriented inwards from the end of supercontigs, see below) revealed a well-conserved LTSa sequence, which could also be identified at some supercontig ends in overhanging long-reads mapped with high confidence. The complete Helitron transposable units thus defined are schematically represented in Figure 5C.

A similar analysis for the 15 copies of Hel15/Hel16 (Figure 5A&B) revealed that most copies ended with an unusual Mosaic satellite (presented in detail in the next section; cf Figure 6A) followed by one of several alternative RTSs (RTSf-m) that could be identified for 13 of the 15 copies (SupFigure16). The latter end with a CCAA motif, like the Hel01/Hel03/Hel04 element copies. At their left end, all 15 copies start with LTSb, homologous to LTSa, or a shortened variant LTSc (Figure 5A&B, SupFigure16A).

**Figure 6.**
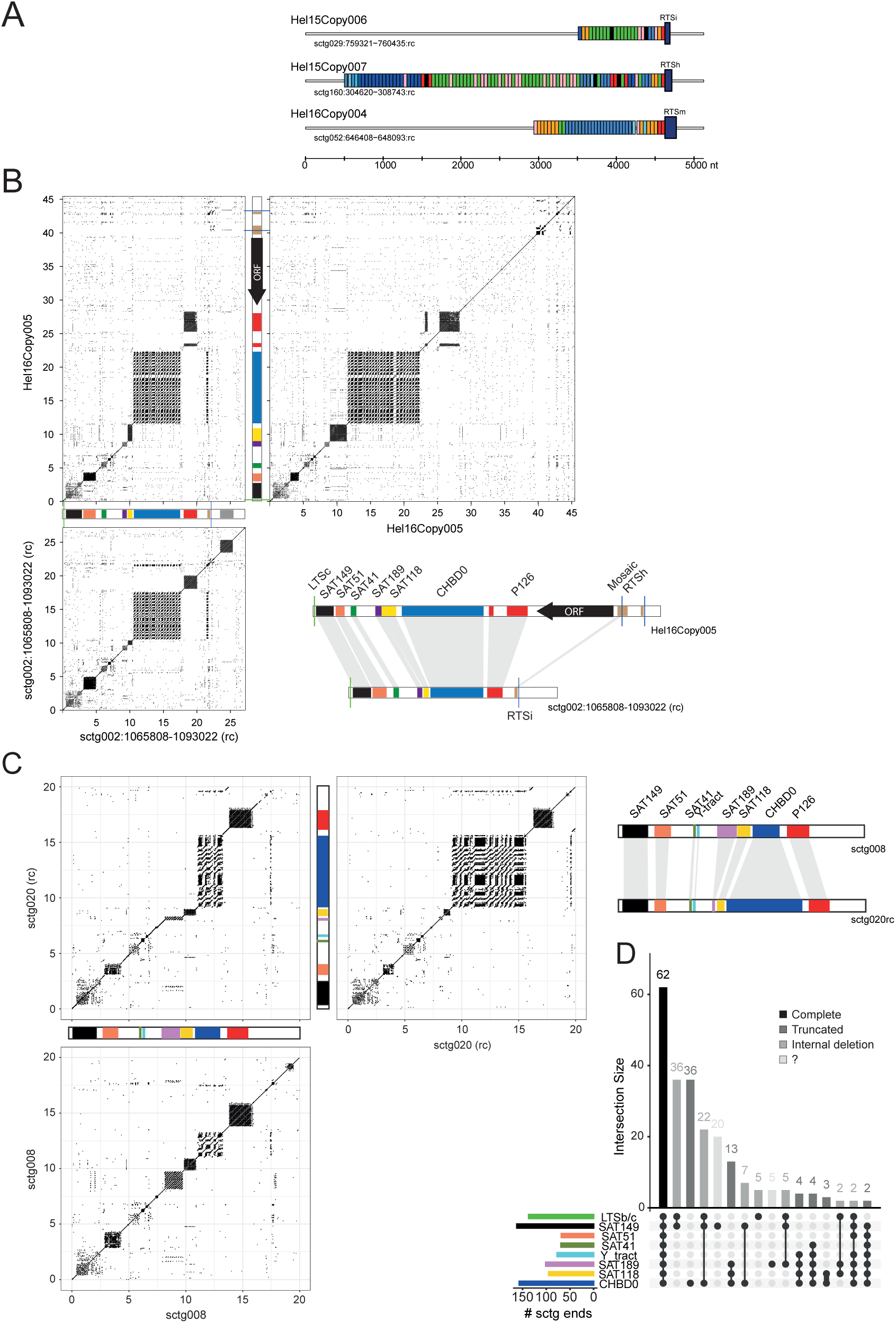
Non-autonomous Helitrons and satellites of clade B Helitrons (Hel15/16). (A) Three examples of the structure of ’Mosaic’ satellite/RTS at the end of Hel15Copy006, Hel15Copy007 and Hel16Copy004. Each colored bar represents an occurrence of a mosaic repeat unit. The RTS are drawn as taller blue rectangles. (B) Three dotplots compare the Hel16Copy005 with itself, with a locus that contains a complete satellite array (right end of sctg002), and this locus with itself. To the right, a schematic comparison of the two loci illustrating loss of the ORF (C) Comparison of the same satellite array found at the extremities of 2 different supercontigs. Dotplots of the 20 kb extremities of supercontigs against themselves show the characteristic series of tandem repeats that appear as squares around the diagonal of identity (SupData4). A motif consisting of at least 100 successive pyrimidines (C or T) was designated “Y_tract”. Dotplots of the 20 kb left end of sctg008 (bottom left) and the 20 kb right end of sctg020 (top right, reverse complement) are shown with these 8 motifs. A dotplot of these two different supercontig ends against each other (top left) shows the same series of motifs, but with different numbers of repeats, so that the satellites now appear as rectangles with a trajectory that diverges from the diagonal of identity. A solid line connecting the rectangles indicates little divergence of the unique sequences between satellite blocks (> 70% identity threshold, 28 nt windows). The diagram (far right) schematizes the expansion or retraction of the tandem repeats. (D) UpSet plot of the association of different satellites found at supercontig ends (n=2 x 187), using the data in SupData4. Each dot indicates the presence of one of the minisatellites. The barplot over the table shows their grouping in supercontig ends. The barplot to the left of the table shows the total occurrence of the different motifs. The black histogram bar shows the supercontig ends with a complete and ordered satellite array (LTSb/c, SAT149, SAT51, SAT41, Y_tract, SAT189, SAT118 and CHBD0). The grey bars show supercontig ends with a truncated and/or deleted satellite array. The very light grey bars represent cases that are difficult to interpret.

Alignment of Helitron copies around LTS and RTS not only show that sequences diverge before LTS and after RTS, but also that there are C_4_A_2_ telomere repeats before LTS and after RTS, suggesting that the copies may have inserted into the telomere repeats. The example in Figure 5C (Hel01Copy009) has a short run of telomere repeats before LTSa and a longer run after RTSb. That this is the general rule was shown by alignments around all putative LTS (SupFigure17B) and RTS (SupFigure17C for clade A1 and SupFigure17D for clade B), whether or not they are associated with full-size ORFs (Methods). The LTSs are often, but not always, flanked by telomere repeats that are at the very beginning or end of the supercontig (SupFigure17B). Remarkably, the RTSs are always followed by short stretches of telomere repeats with the same orientation. This is particularly clear for RTSa-e associated with cladeA1 (SupFigure17C; see also SupData4 scale drawings of supercontig ends).

In all cases where putative LTS and RTS sequences were identified, Helitrons point away from the chromosome’s end towards its central, MAC-destined region: LTS at the end, RTS towards the center. Interestingly, the orientation of full-length ORFs with respect to the orientation of the complete Helitron is variable depending on the element (Figure 5B). In clade A1, the full-length ORFs are oriented in the same direction as the Helitron (except for Hel06), whereas clade B Helitron ORFs have the inverse orientation, pointing towards the chromosome end (except for Hel13). One way or the other, the orientation is always the same for all copies of each element (Figure 5A&B, SupFigure16A, SupData4).

The Pgm-independent genomic compartment is restricted to chromosome ends as are most Helitrons with full-size ORFs. Chromosome ends are also enriched in certain minisatellites (Figure 4). We therefore used previous minisatellite annotations for *P. tetraurelia* (Figure 1C, SupFigure5) and dotplots, which can reveal imperfect tandem repeats whatever their size, for each supercontig end (SupData4) and each locus with a full-length ORF, to annotate the satellites within and around Helitrons. As shown schematically in Figure 5B, the pattern of LTS-satellites-RTS for complete Helitrons differentiates sub-clades, hereafter designated “types”. Clade A1 contains 5 types and clade B, 3 types. Satellite annotations are drawn to scale with respect to all full-size ORFs in SupFigure16A. The manually curated Mosaic satellite that precedes the putative RTS of Hel15/Hel16 copies (Figure 6A, SupFigure16, SupData3, SupData4) is unusual in that it is composed from 11 different ∼44 nt segments with distinct sequences, assembled singly or as tandem repeats in an apparently random order. The generally low number of identical tandem repeats in this satellite made it difficult to detect with the automated pipeline (only the “Blue” repeat segment was detected). The Mosaic loci of Hel15/Hel16 copies all end with the “Red” segment and are followed at a short distance (42-168 nt) by the putative RTS (SupFigure16B). The near-uniform size of the repeat units and their amazing random assembly order raise intriguing questions about their origin, dispersal and evolutionary history.

### Non-autonomous Helitrons are abundant at chromosome ends

A search for LTS sequences in the assembly retrieved many copies not directly followed by Helitron ORFs. In the case of LTSa, 10 of them are followed by sequences very similar to the beginning and end of complete type 1 Hel01 copies, from which they may have been produced by recombination of two related satellites, SAT26 and SAT54/81, present 5’ and 3’ of the ORF (SupFigure18). These 10 LTSa-RTSa sequences, 6.5-9.9 kb in length, appear to represent type 1 Hel01 copies that have lost the ORF, as may also be the case for other, truncated loci. In the case of LTSb/c, 7 different satellites occur between the LTS and the ORF of Hel16Copy005 (Figure 5B, Figure 6B). The same array of satellites was found, without the ORF, at the ends of many supercontigs (SupData3&4; Figure 6C).

Their order from the end of each supercontig is consistent with the global analysis of satellite DNA distance from supercontig ends (Figure 4C). Fifteen of the 62 complete copies that start with LTSb or LTSc, end with the Mosaic satellite and/or an RTSf-RTSm sequence (SupData4). This suggests that the multiple copies of this satellite array may represent defective copies of the Hel16Copy005 type in which the ORF and in most cases the Mosaic-RTS are missing. Given the number of copies of this satellite array that appear to be complete, internally deleted or truncated (Figure 6D), we estimate that at least 62 (complete arrays) and up to 196 (including truncated and internally deleted arrays) of these putative non-autonomous Helitrons have inserted at a chromosome end. The conservation of the order of the satellites in the array and of the unique copy sequences that connect them (cf. dotplots: SupFigure4, Figure6C) suggests that a recent burst of transposition activity spread the arrays around the genome, after the divergence of *P. tetraurelia* from other *aurelia* species.

Similarly, 20 other supercontig ends sharing blocks of P126 minisatellites between LTSb and Mosaic- RTS may be ORF-less derivatives of type 1 Hel15 copies (Figure 5B; SupFigure19; SupData4).

The annotation of full-length Hel01/03/04 and Hel15/16 elements indicated that they contain a variety of minisatellites which may have been dispersed at the ends of MIC chromosomes by transposition. The annotations further suggested that partial, non-autonomous Helitron copies lacking the ORF are much more frequent than full-length ones. If these characteristics hold for other Helitron families with fewer ORF copies, then a large fraction, if not all, of the Pgm-independent compartment would appear to result from Helitron-mediated transposition. To test this hypothesis, we made a compendium of Helitron full-size ORFs, RTS, LTS and the specific satellites which may be assumed to be part of transposable Helitrons located at supercontig ends (50 kb ends; SupData4).

We found that 254 supercontig ends have Helitron annotations and 90% of these end with a Pgm- independent compartment, while 83% of the 74 extremities with no evidence of Helitrons do not end in a Pgm-independent region (Figure 7A). A more chromosome-centric vision shows that the 161 chromosome-size supercontigs (Figure 2D, SupData1) usually have Helitron-associated annotations at one or both ends (both ends, n=98; one end, n=58; neither end, n=5; Figure 7B). The hypothesis that the Pgm-independent compartment found at chromosome ends results from Helitron transposition is well-supported by current data.

**Figure 7.**
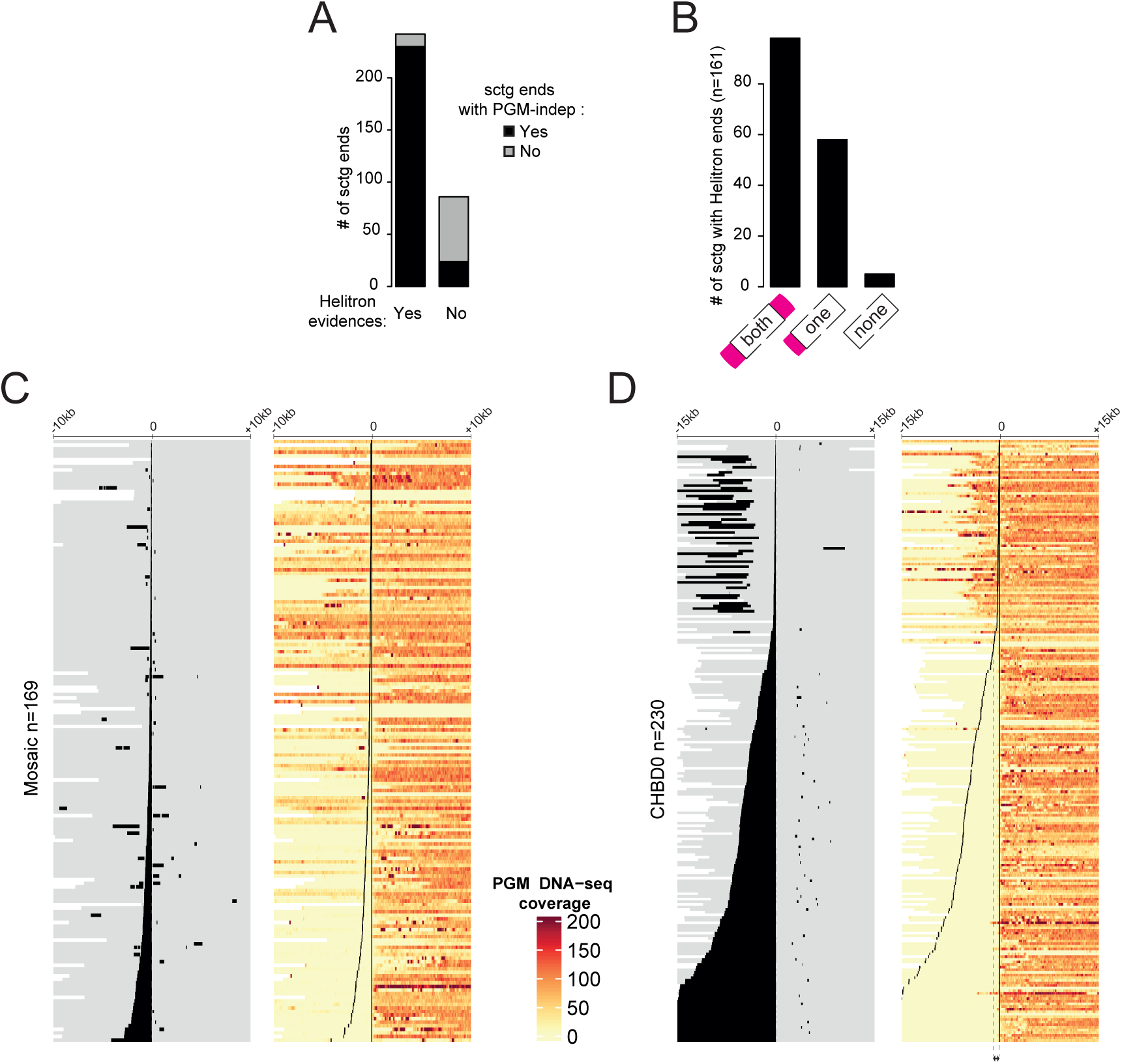
Chromosome ends contain Helitron-associated sequences that define the PGM-independent frontier. (A) The number of supercontig ends for the 161 supercontigs (Figure 2G), with and without Helitron evidence at their ends. Helitron evidence can be an LTS, an RTS, a Helitron ORF, or a characteristic Helitron-associated satellite. See SupData4 for details. In both categories, supercontig ends without terminal PGM-independent regions are grey. (B) The number of supercontigs with Helitron evidence on both sides, on one side or on neither side of the end. (C) Heatmap showing PGM RNAi DNA-seq coverage (bin = 1) within a 20 kb region centered on the most internal boundaries of Mosaic satellite loci (n = 169). The loci (rows) are ordered according to the size of the Mosaic of interest. On the left, the black lines show the position of the Mosaic and the grey parts show the flanking regions. Note that a window can contain multiple occurrences of Mosaic. The white areas show regions that cannot be displayed because they are outside the supercontig. (D) Occurrences of CHBD0 closer to each other than 1 kb were fused (n=230) and displayed as in (D), using 30 kb windows. Note that the largest CHBD0 (bottom rows of heatmap) extend beyond the window. A sharp frontier is visible for CHBD0 loci larger than 1 kb.

A striking property of the Mosaic satellite, and also of the CHBD0 satellite found in the Hel16Copy005-derived satellite array, is their localization on the Pgm-independent side of the frontier between distal Pgm-independent and more proximal Pgm-dependent genomic compartments. This is illustrated in Figure 7C&D for all occurrences of Mosaic (n = 169) and CHBD0 (n=230), using heatmaps of Pgm read coverage (Methods) in large (10 or 15 kb, respectively) intervals flanking the frontier. In both instances, the innermost, proximal extremity of the satellite corresponds almost exactly to the frontier between the Pgm-independent and Pgm-dependent compartments. We also note that the larger the satellite, the sharper the frontier. In some cases (n=39) both CHBD0 and mosaic were found in the same 30 kb windows (SupFigure20). Overall, the compartment frontier appears to be determined by the largest satellite, Mosaic or CHBD0.

## DISCUSSION

Here, we report chromosome-scale germline genome assemblies for seven members of the *Paramecium aurelia* species complex and a nearly telomere-to-telomere long-read assembly and genetic linkage map for *P. tetraurelia*, a model extensively used to study programmed DNA elimination and non-Mendelian heredity [81]. In addition to providing an important resource, the unforeseen results of our study are (i) a karyotype with ∼160 tiny chromosomes; (ii) a novel clade of Helitron-like elements that have remained active in all *aurelia* lineages, with 9-10 kb ORFs; (iii) full-length autonomous and numerous non-autonomous Helitron copies that have invaded chromosome ends; and (iv) chromosome ends eliminated very early in development by an unknown mechanism.

### Costly karyotype

Narrowing down the karyotype of *P. aurelia* has been no small effort given the difficulty of purifying MIC DNA and the absence of genetic or physical maps. Short-read assemblies for 7 *Paramecium* species allowed us to establish TE landscapes and discover minisatellites although the scaffolds did not usually extend into the telomeric regions. Our long-read assembly overcomes the fragmentation issues of the short-read assemblies for the *P. tetraurelia* model. It uses DNA from new MACs depleted of Ezl1, a core component of the PRC2 complex required for most DNA elimination (1/3 of IESs are however eliminated). It is important to keep in mind that the MIC *EZL1* supercontigs provide the best germline assembly possible given current technological bottlenecks, but they do not constitute the reference MIC genome.

Several convergent lines of evidence are in favor of a karyotype for *P. tetraurelia* with about 160 chromosomes, from 300 kb to 1.2 Mb in size. The first line of evidence is the assembly itself, which includes 161 supercontigs containing MAC-destined sequences. Although only 33 of these supercontigs are telomere-to-telomere, the size distribution of supercontigs capped by telomeric repeats at both ends is similar to that of other supercontigs (Figure 2), which suggests that they are also quasi complete. Based on the independent observation that the Pgm-independent compartment is a marker of chromosome ends (Figure 3), we counted 241 supercontig extremities with Pgm-independent genomic compartments (including 85 supercontigs where they are present at both ends) (Figure 7A&B, SupData4). Taken together, this is strong support for a karyotype consisting of at least 120, and possibly as many as 161 chromosomes.

A second line of evidence comes from analysis of aneuploid clones. Indeed, we noticed that chromosomal segregation errors are frequent in *P. tetraurelia*. Among the 39 independent F2 clones that we sequenced, 8 (21%) present evidence of higher read coverage (generally twice more) affecting one or more supercontigs, compared to the global median read coverage (SupFigure9A&B). We also analyzed 74 previously published DNA-Seq datasets prepared from 100% homozygous cell lines. We observed 21 additional cases (i.e. in 28% of cell lines, SupFigure9C), which shows that these duplication events are not specific to the F1’s of this particular hybrid genetic cross. These events affect entire MIC supercontigs 92% of the time i.e. the high coverage is seen all along the supercontig. This indicates that there was a segregation error during meiosis leading to extra chromosome copies (i.e. aneuploidy). Under the hypothesis that a supercontig corresponds to a fragment of a chromosome, one would expect the supercontig(s) corresponding to the other fragment(s) to also be affected each time the chromosome is duplicated. Thus, when a supercontig is detected as duplicated in an aneuploid clone, but is not co-duplicated with any other supercontig, we can conclude that the corresponding MIC chromosome does not include any other MAC-destined supercontig. Using this logic, we identified 19 supercontigs for which we could validate a one-to-one correspondence to a MIC chromosome (SupFigure9C). The size distribution of these 19 validated supercontigs is similar to that of other supercontigs (SupFigure9D), and the proportion that are telomere-capped at both ends (4/19=21%) is the same as in the entire dataset (33/161=20%) (Figure 2G). Although the analysis does not have the power to be exhaustive, it does show that the 161 supercontigs, even if not telomere-to-telomere, are likely to correspond to individual MIC chromosomes.

A last line of evidence comes from analysis of the recombination map. The number of crossovers (COs) per chromosome is generally tightly constrained by both an upper and a lower limit [82]. A survey covering a wide range of taxa showed that in 97% of cases, each chromosome tetrad receives a minimum of 1 and a maximum of 7 COs per meiosis [82]. The lower boundary reflects the requirement of having at least one CO to ensure the proper segregation of chromosomes during meiosis [82–85]. Because of this constraint, small chromosomes have a higher recombination rate (in cM/Mb) than larger ones. Interestingly, *P. tetraurelia* fits perfectly with this pattern: for each of the 161 supercontig groups that could be genotyped, we detected between 0.7 and 3.7 CO per F2 (i.e. 1.4 to 7.4 CO per chromosome tetrad) (Figure 2A). Moreover, the size of the supercontigs is negatively correlated with their recombination rate (Figure 2B). These observations strongly suggest that *P. tetraurelia*, like most eukaryotes, must have at least one CO per chromosome tetrad per meiosis. Under the assumption that supercontigs corresponded to fragments of chromosomes, one would have expected many of them to have a genetic length below 50 cM (i.e. below 0.5 CO per F2: Figure 2A, dotted line). The fact that even the shortest supercontig groups are above this threshold suggests they do correspond to (quasi) full-length MIC chromosomes.

The reason why *Paramecium* has so many chromosomes is that its genome has been shaped by a series of at least 3 whole genome duplications (WGDs) [7,86]. Such events have occurred repeatedly in various eukaryotic lineages such as plants [87], vertebrates [88–90], and yeast [91]. The pathway of resolution of these dramatic events, whose immediate consequence is to double the number of chromosomes, is that one of each pair of duplicated genes accumulates mutations and eventually decays while the other copy remains under selection, leading to a reduction in genome size. In most taxa, chromosome end-to-end fusion and/or other large-scale rearrangements subsequently reduce the total number of chromosomes back towards the pre-WGD number [87–89,92,93]. However, *Paramecium* appears to be less prone to chromosome fusions/rearrangements than most other paleopolyploid eukaryotes: the 3 most recent WGDs remain clearly visible in contemporary genomes, with extensive syntenic blocks of genes and only a few translocations [22]. Thus, the last 3 WGDs in the *Paramecium* lineage have led to an 8-fold karyotype expansion, from 2n = 20 chromosomes to 2n = 160. A similar pattern has been reported in ferns (from 2n = 18 chromosomes in *Salvinia natans* to 2n = 1440 in *Ophoglossum reticulatum;* [94]).

The high number of chromosomes is expected to increase the risk of producing aneuploid offspring. For instance, in budding yeasts, the rate of meiotic segregation error is 0.15% per chromosome per tetrad [95]. With such an error rate, the frequency of aneuploid meiosis is expected to increase from ∼2% in a species with 16 chromosomes (like budding yeasts) (1-0.9985^16), to 21% in a species with 160 chromosomes (1-0.9985^160). In *P. tetraurelia*, we observed that 21% to 28% of clones carry at least one aneuploid chromosome. It should be stressed that these numbers are certainly underestimates of the true chromosomal segregation error rate, because only the viable offspring can be observed. Hence, it is hard to escape the fact that such a high rate of aneuploidy will have a fitness cost. This raises the question as to why *Paramecium* maintains so many chromosomes. We can hypothesize that part of the answer is that end-to-end fusion of chromosomes, as found in other taxa, may be inhibited by the Helitrons that structure and cap chromosome ends.

### A new class of Helitrons

Eukaryotic Helitrons are a class of DNA transposons proposed to transpose via a rolling-circle replication mechanism [96]. The first elements to be described lacked terminal inverted repeats but possessed conserved 5’-TC and CTRR-3’ termini as well as a short hairpin close to the 3’ end and were found to insert within 5’-AT-3’ dinucleotides with no target site duplication. Related elements, Helitron2 and Helentron [97], possess short asymmetric terminal inverted repeats and a hairpin structure at each end. Helitron2 transposons insert within 5’-T(T/C)-3’ dinucleotides while Helentrons insert within 5’-TT-3’ and contain an additional apurinic-apyrimidinic endonuclease domain within the transposase [98–101]. Our phylogeny (Figure 1H) shows that all *Paramecium* elements emerge in a specific clade distinct from the two previously defined clades, HLE1 and HLE2 [52], supporting ancient acquisition and diversification of Helitrons in *Paramecium*. We did not find copies of all 29 elements in each of the species examined, which may be due in part to the incomplete and fragmented nature of short-read assemblies at the ends of chromosomes.

Nevertheless, elements from all 3 major clades (A1, A2, and B) were found in each species. Although frequent Horizontal Transposon Transfer of Helitrons is well-documented in other eukaryotic taxa [102–104], we found no evidence for Horizontal Transposon Transfer between *Paramecium* species: the transposase tree is consistent with the species tree for those elements present in several species (SupFigure7B).

An additional argument for ancient *Helitron* acquisition and diversification in *Paramecium* is the remarkable presence of an IES that interrupts a conserved region of the transposase ORF at exactly the same position in all copies of 5 different elements in an A2 subclade (SupFigure7). These elements are not in the Pgm-independent compartment and at least some copies are eliminated sufficiently late to allow the prior excision of the IES by the Pgm excision complex (SupFigure7 and 14), leaving a narrow window for expression of the transposase from the developing MAC. This IES likely derives from the insertion of a *Tc1/mariner* element (or a mobile, non-autonomous IES; [14]) into an ancestral Helitron ORF before diversification of the elements and before subsequent *aurelia* speciation, which is thought to have started several hundred million years ago [105]. This indicates that the expression constraints due to the IES have not led to the loss of that subclade, and more generally that Helitron transposition activity persisted over a very large time scale in *P. aurelia* species.

The other available ciliate germline genome assemblies do contain evidence of Helitrons although they are not abundant (< 0.5 % of the genome) and are highly fragmented [17,18]. We found homology to *Paramecium Helitron* ORFs by tblastn at only a few *T. thermophila* loci, the best scoring hits located in centromeric regions of some of the 5 MIC chromosomes [18]. In *Oxytricha*, more distantly related to *Paramecium*, 3 of 301 RepeatScout consensus sequences were classified as *Helitrons* in a genome with very high repeat content [17], although we failed to obtain any tblastn hit on that genome. It is unclear whether all ciliate Helitrons belong to this same, third clade since the available genomes and annotations are too limited.

Two unusual features of *Paramecium* Helitrons suggest a unique integration specificity that targets them to telomeric repeats. First, the identification of LTS and RTS sequences for clade A1 elements Hel01/Hel03/Hel04 and clade B elements Hel15/Hel16 revealed an absolute orientation bias at chromosome ends, the LTS always being closest to the 5’ end of supercontigs, and the RTS more internal. This is true for both types of elements, even though the ORFs are in opposite orientations in the two cases (Figure 5B). While we have not identified the LTS and RTS of other elements, we note that ORF orientation is always the same for all genomic copies (pointing away from the supercontig end or towards it, depending on the element), suggesting that this absolute strand bias is a general feature of *Paramecium* Helitrons. Secondly, all annotated full-length copies are flanked by CCC(C/A)AA telomere repeats on both sides, though these are often reduced to just a few degenerate repeats 3’ of the RTS of Hel15/Hel16 copies (SupFigure16). These repeats vary widely in length and sequence (alternation of C_4_A_2_ and C_3_A_3_ repeats) among copies of the same element and thus do not appear to be part of the TEs themselves, unlike the terminal telomere repeats present in the Tc1/mariner element TBE1 in *Oxytricha* [106,107]. Nor can they be presumed to have been added by telomerase to the ends of some linear transposition intermediate before integration, as proposed for the Tel-1 element in *Tetrahymena* [108], since this would result in GGG(G/T)TT repeats 3’ of the RTS instead of the observed CCC(C/A)AA. A more likely hypothesis is that these Helitrons specifically insert into the C-rich strand of telomere repeats: this would constrain their orientation at chromosome ends, fully accounting for the absolute bias observed. Insertion within MIC telomeres would both lengthen chromosomes and create internal blocks of telomere repeats of the same polarity, which are indeed frequently found near supercontig ends.

Another remarkable feature is the abundance and variety of satellites, duplications and higher-order repeats present in annotated full-length copies, between the ORF and the conserved ends. Figure 5 and SupFigure16 only show the most conspicuous satellites that could be detected automatically, but closer inspection further reveals old, degenerate satellites, microsatellites and non-contiguous duplications. Together these different types of repeats occupy almost all of the non-coding sequences on both sides of the ORF, which represent from ∼50% up to 75% of the total lengths (SupFigure15). HLEs are prone to capture adjacent genes or other genomic sequences during transposition, and different models have been proposed for how this may occur [80,97,109,110].

*Paramecium* Helitrons may have captured satellites – but no gene fragments or other MAC sequences – through preferential integration into the Pgm-independent compartment, which is devoid of any gene but replete with satellites.

The occurrence of distinct sets of satellites among copies of the same element (defining Hel01 types 1-5 and Hel15/Hel6 types 1-3, Figure 5) indicates that they can change with transposition cycles. It should be noted that the presence of captured satellites between the LTS and the ORF cannot be directly explained by the 3’-bypass model for capture, in which failure to terminate the unwinding of the transposing strand at the end of the RTS results in the inclusion of downstream flanking sequences until another possible termination site is reached. However, it could be explained by insertion of one Helitron copy within another, followed by deletion of the termini and ORF of the internal copy. Nested insertions could also explain the formation of elements with a reversed ORF orientation, as has been proposed for some *Fusarium* elements [101]. In some animals and plants, defective copies of Helitron-like elements lacking the ORF were shown to contain various types of satellites, including some with centromeric functions, which are thought to have been dispersed across genomes through non-autonomous transposition [100,111–113]. Although ORF-containing, autonomous elements harboring the same satellites are rarely observed in *P. tetraurelia*, we showed that at least 2 types of potentially autonomous elements (Hel01 type 1, SupFigure18; Hel16 type 3, Figure 7B) appear to have given rise to ORF-less derivatives which contain the same satellite sets and are present at many more supercontig ends. Their dispersal by non-autonomous transposition may be relevant to the developmental processing of germline chromosomes, since two complex satellites from Hel16 type 3 (CHBD0 and the Mosaic) precisely coincide with the boundary of the Pgm- independent compartment. Helitron-mediated non-autonomous transposition could also have contributed to the multiplication of centromeres -- whose sequences have not yet been identified – as implied by the peculiar karyotype of *P. aurelia* species, consisting of a large number of very short chromosomes.

Our data indicated that the ORFs of *Paramecium* Helitrons have been under purifying selection. This may simply reflect recent transposition activity, in line with the idea that rapidly disappearing ORFs are continuously replaced by new insertions. An alternative, but not exclusive, explanation is that Helitron transposases may provide advantageous or essential functions to the host. First, they may be required for their own elimination during development of the somatic MAC (since this does not involve the Pgm endonuclease), thus limiting the rate of transposition in the germline MIC. If we assume that the MAC ensures all gene expression, then transcription of a Helitron transposase ORF would originate from an element in the new developing MAC or from an element that failed to be excised from the maternal MAC. Another possibility is that elements that recurrently insert within telomeric repeats may contribute to the protection of chromosome ends against replication-dependent erosion, much as non-LTR retrotransposons do in some *Drosophila* species lacking telomerase (Saint-Leandre et al. 2019). *Paramecium* Helitrons may also play some structural role at chromosome ends, for instance in the formation of the “bouquet” during meiotic prophase, a cluster of telomeres attached to the nuclear membrane which is thought to facilitate homolog pairing in many organisms [115,116]. They could even act like the meiotic pairing centers of *C. elegans*, specialized chromosome sites that stabilize homolog pairing [117,118]. Frequent transposition of such a diverse set of elements at chromosome ends would then rapidly affect homolog recognition between populations, explaining the high rates of F2 lethality frequently observed in crosses between different isolates of the same species [15], and possibly driving speciation.

Another exciting possibility is that Helitron transposases may contribute to the programmed rearrangement of rDNA loci. Indeed, in *Paramecium*, 9 kb rDNA units encoding 18 and 28S rRNA are found as tandem arrays within dedicated MAC chromosomes, whereas only 4 single rDNA units have been found in the MIC genome. These MIC loci contain a little more than a unit, with direct repeats of ∼ 300 bp at either end, and differ from each other by the sequence of the non-transcribed spacer [119]. It was proposed that these MIC units are excised as circles via homologous recombination between the ends during MAC development, and that rolling-circle replication then generates tandem repeats. This model would explain the observation that adjacent units in the MAC tandem repeats usually have the same spacer [119]. The tempting hypothesis that a Helitron transposase is involved in this process would reveal a second example of TE domestication, alongside that of the Pgm endonuclease [27].

### Models for Pgm-independent early elimination of chromosome ends

The germline genome of *P. tetraurelia* is organized into distinct regions based on their developmental fate. The constitutive and the variable MAC regions are, respectively, always and partially retained in the new MAC. The MIC-limited region, which includes IESs, TEs and satellites, is consistently eliminated during MAC development. Previous studies have demonstrated that separate yet overlapping pathways are involved in the Pgm-dependent elimination of IESs and TEs [34–36] (Figure 8A). Our findings now reveal the existence of a completely novel pathway for the elimination of the Pgm-independent compartment, first reported by [16]. This compartment is eliminated very early, before IESs are excised (Figures 3C and 8A) [26]. This newly defined compartment is located at the ends of germline chromosomes and consists of most Helitrons with full-size ORFS and multiple non-autonomous Helitron derivatives, including the CHBD0-containing satellite array (Figures 5-7, SupFigure15-17). The latter is positioned directly adjacent to a Pgm- dependent compartment, prompting the question of how the boundary between these two compartments is defined. We have shown that Helitron-associated Mosaic and CHBD0 satellites define the boundary (Figure 7C-D). Interestingly, predicted G4s are not found in the Mosaic satellite, but are significantly enriched within the CHBD0 repeat (70% of predicted G4 are in CHBD0) located in the proximal part of the Pgm-independent compartment (Figure 4A). This observation suggests that G4s may play a role in defining the boundaries between the compartments. This is reminiscent of the function of the G4-binding protein Lia3, which plays a key role in definition of the boundaries of germline-limited sequences in *Tetrahymena* [120]. Lia3 interacts with G-rich polypurine tracts (A_5_G_5_) located on either side of germline-limited elements, which are eliminated by the Pgm homolog, Tpb2p. In the absence of Lia3, alternative boundaries are used. Notably, no homolog of Lia3 has been identified in the *Paramecium* genome as Lia3 appears to be specific to *Tetrahymena*.

**Figure 8.**
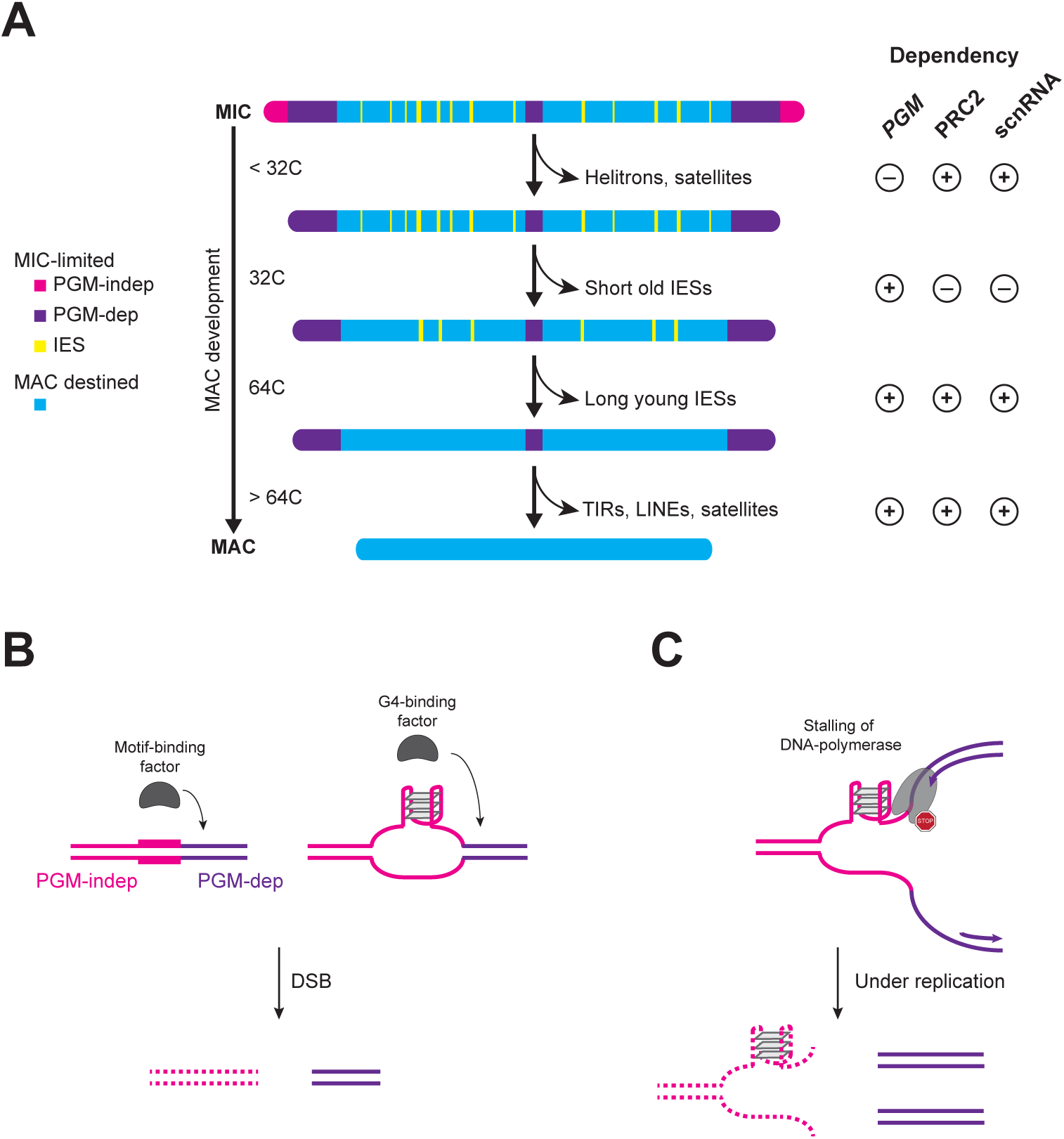
Models for Pgm-independent early elimination of chromosome ends. (A) MIC chromosomes end with sequences eliminated independently of Pgm endonuclease activity (pink), and contain Helitrons and associated satellites. The elimination of these regions involves the scnRNA RNA pathway and PRC2. These regions are eliminated earlier in new MAC development (before 32C endoreplication state) than Pgm-dependent regions (purple). Pgm-dependent DNA elimination [26] occurs between 32C-128 C endoreplication state and concerns all IESs and remaining TEs. The MAC-variable regions (seagreen) are not eliminated completely and contribute to the heterogeneity of MAC chromosomes [6]. The gene-rich constitutive-MAC (blue) regions are found in all MACs. (B) Model to explain the mechanism behind the disappearance of PGM-independent/Early eliminated regions. The first hypothesis is that a factor binds a motif (e.g. the mosaic satellite) or a DNA structure (e.g. G4) and this binding promotes introduction of a double-strand break (DSB), so that the PGM-independent region would not be replicated. (C) A second, non-exclusive hypothesis is that a particular DNA conformation, such as G4, would stall the progression of the DNA polymerase resulting in under-replication of the end of the chromosome.

To explain the elimination of the Pgm-independent compartment, we propose two alternative, non-exclusive models. In the first scenario (Figure 8B), the presence of the Mosaic satellite or G4s promotes the formation of DNA double-strand breaks at the boundary between the two compartments. One possibility would be that the Mosaic sequence and the G4 structures mediate the recruitment of an unknown endonuclease either directly or indirectly through a G4/Mosaic- binding protein, and this would eventually lead to the elimination of the G4/Mosaic-containing region. Similarly, during class switch recombination in B cells, double-strand breaks are initiated by base modification catalyzed by AID (activation-induced deaminase) that directly binds to G4s [121]. Alternatively, one can imagine that G4 structures formed within the CHBD0 are not properly resolved and stall the DNA replication machinery, which can also cause DNA double-strand breaks [122,123]. Indeed, highly stable G4s can form during replication, and the failure to resolve G4s results in genome instability. G4-associated genome instability appears to largely result from the stalling of the replicative polymerase at the G4 structure and subsequent uncoupling between the replicative helicase and the DNA polymerase [124].

In the second scenario (Figure 8C), the stalling of the replicative DNA polymerase at G4 structures results in under-replication of the Pgm-independent compartment. We imagine here that the resolution of G4 structures during DNA replication prevents mutagenic events, including double-strand break formation. However, in the absence of G4 resolution, the prolonged stalling of the DNA polymerase at G4s and a permanent uncoupling between the replicative helicase and the polymerase would lead to under-replication of the G4-containing region. This scenario is very reminiscent of common fragile sites in vertebrates [125] and raises the possibility that fragile sites, like the one responsible for some *Paramecium* cortical mutants [126], could be developmentally programmed for DNA elimination, via under-replication (Figure 8C).

To explain why depletion of the scnRNA-PRC2 pathway results in normal replication of the G4- containing region, we envision that G4s are resolved under such conditions, preventing under-replication. Unfolding of G4s requires the functions of specialized DNA polymerases, helicases, and single-stranded DNA binding proteins [124]. Remarkably, Helitrons contain a putative Pif1 helicase, known in other organisms to be involved in G4 resolution [124]. The expression of at least one gene required for G4 resolution might be negatively controlled by the scnRNA-PRC2 pathway, perhaps through the deposition of repressive histone marks. Under normal conditions, the gene would be repressed, leaving the G4s unresolved, while it would no longer be repressed in scnRNA-PRC2 depleted cells, allowing the resolution of G4s and endoreplication of the region. It is also conceivable that, similarly to G4s, Mosaic can form DNA structures that become destabilized in the absence of the scnRNA-PRC2 pathway. Further investigation will be necessary to elucidate the underlying mechanisms.

Our data reveal a unique mechanism that specifically eliminates the ends of germline chromosomes. Interestingly, the loss of sub-telomeric and telomeric sequences of the germline chromosomes also occurs in *Tetrahymena* [18] and all nematode species where PDE has been reported [127–131]. In the free-living nematode *Oscheius tipuale*, no internal break site is generated and only the sub- telomeric regions that contain Helitron-like elements are eliminated [128,129]. A conserved 30-nt palindromic motif is associated with the break site [129], while no sequence feature has been identified in the parasitic nematodes Ascaris and Parascaris [127,132–134]. So far, there is no report of a protein that physically interacts with the motif or at the break sites, and the underlying mechanisms remain to be discovered. Similarly to the potential role of G4s in *P. tetraurelia*, DNA structures may be important for the removal of germline chromosomal ends in these organisms. The recurrent elimination of the terminal regions of germline chromosomes across various phyla may indicate convergent evolution and suggests that these germline-specific regions could have a crucial role for the chromosomes, that is yet to be discovered.

## Conclusions

The key finding of our study is that the germline genome of *P. aurelia* consists of numerous tiny chromosomes (n ∼160, 300 kb to 1.2 Mb in size), with an exceptionally high recombination rate (420 cM/Mb). The chromosomes terminate in a novel genomic compartment that appears to be the result of transposition of a new class of Helitrons, that insert in C_4_A_2_ telomere repeats. This compartment is eliminated very early during development of the somatic nucleus, by a mechanism independent of the well-studied PiggyMac domesticated transposase complex that was thought until now to be responsible for all transposon elimination.

## MATERIALS AND METHODS

### Strains and cell culture

*Paramecium* cells were grown in wheat grass powder (WGP) (Pines International) infusion medium bacterized the day before use with *Klebsiella pneumoniae*, unless otherwise stated, and supplemented with 0.8 µg/mL β-sitosterol (Merck). Cultivation and autogamy were carried out at 27°C as described [135,136].

The *Paramecium* strains used in this study are listed in SupTable1: *tetraurelia* (strain 51, unless stated otherwise), *octaurelia* 138, *biaurelia* V1-4, *pentaurelia* 87, *primaurelia* AZ9-3, *sonneborni* ATCC 30995, *sexaurelia* AZ8-4.

### Illumina sequencing and assembly of vegetative MIC DNA

Previously published [14] MIC DNA small-insert libraries of vegetative MIC DNA were used. For assemblies, read pairs containing a MAC IES junction, identified using ParTIES software [78] and MAC reference genomes [11,14], were removed to reduce ambiguity. The 250 nt filtered read pairs were fused with Flash version 1.2.11 [137] using default parameters to obtain ∼450 nt fragments. The fragments were assembled into MIC contigs by the Newbler overlap-layout-consensus assembler (version 2.9 Roche Diagnostics), with parameters (-mi 100 -ml 99). Because of the DNA quantities required for mate-pair libraries, we used DNA from new MACs after *PGM* silencing by RNAi (*PGM* DNA) [25]. It was not possible to obtain *PGM*-RNAi DNA for *P. sonneborni*, so only contigs are available for this species.

Large-scale cultures of *tetraurelia* 51, *octaurelia* 138, *biaurelia* V1-4, *pentaurelia* 87, *primaurelia* AZ9- 3, and *sexaurelia* AZ8-4 were collected at the end of MAC development in *PGM* RNAi conditions.

Whole genomic DNA was isolated and used to construct four *Illumina* mate-pair libraries with insert sizes of 3-5 kb, 5-8 kb, 9-11 kb and 14 kb (only the first three sizes were available for *P tetraurelia*) to build scaffolds with the MIC contigs. The scaffolder SSpace basic v2 [138] was used, with parameters (-a 0.5 -k 7). Gap closing was accomplished with SOAPdenovo2 GapCloser software [139]. Assembly statistics are described in SupTable1 and sequencing datasets used or generated for this study are described in SupTable2.

### Sequencing DNA from sorted new developing MACs after RNAi-depletion of factors

As in [140], large-scale, 500 ml cultures of *P. tetraurelia* were collected at T30 (30 hours after T0 of autogamy, time at which 50% of cells have fragmented old MACs). RNAi KD was performed using L4440 derivatives carrying the following inserts: *DCL2/3*, *DCL2/3/5*, *DCL5* [35,141] or *TFIIS4* [36]. As detailed in [26], nuclei of the RNAi-depleted cells were enriched by centrifugation through a 2.1 M sucrose layer prior to flow cytometry using PgmL1 antibody labeling to separate new developing MACs from old MAC fragments. Sequencing librairies were then made using total DNA from the sorted new MAC fraction with the NEBNEXT Ultra II DNA library Prep Kit (New England Biolabs) for *DCL2/3*, *DCL2/3/5*, *DCL5,* or as previously described [26] for *TFIIS4*. The libraries were subjected to paired-end sequencing on the Illumina platform.

### Oxford Nanopore sequencing and assembly of *P. tetraurelia EZL1*-RNAi DNA

Unrearranged DNA was extracted from new developing MACs of *Paramecium tetraurelia* Ezl1- depleted cells [34,46]. Four libraries were generated and sequenced using Oxford Nanopore Technology (ONT). The reads were filtered with Porechop (https://github.com/rrwick/Porechop). Reads containing a MAC IES junction were removed to reduce ambiguity, as described above for the Illumina assemblies. Data were assembled with SMARTdenovo [142] and polished with Racon [143] using the ONT reads as in [144]. The assembly was then subjected to two successive waves of polishing with Pilon [145] (v1.23 --mindepth 10 --minqual 30) using the sorted vegetative MIC Illumina reads. We used the recombination map (see next paragraph) to break chimeras or join contigs to obtain supercontigs. The assembly "ptetraurelia_51_EZL1_SmartDeNovo_v1.0" is throughout the manuscript referred to as the “MIC *EZL1* assembly” (see SupTable1).

### Linkage map and recombination rate for *Paramecium tetraurelia*

To build a genetic map of *P. tetraurelia*, we performed genetic crosses between strain 51 and strain 32. F1 clones from several different pairs of conjugants were allowed to undergo autogamy to generate entirely homozygous F2 clones. Total cell DNA was extracted from 42 independent F2 clones as described in [146]. The DNA samples were sequenced to an average read depth of ∼50X within MAC-destined regions (given that we extracted total DNA, most of sequence reads derive from the MAC genome, and hence we have a very limited coverage in MIC-restricted regions). Reads were mapped to the *P. tetraurelia* strain 51 MIC genome assembly with BWA-MEM v.0.7.15 [147] and then filtered with SAMtools v.1.3.1 [148]. Duplicate reads were removed by Picard tools v.1.98 [149]. We called variants with GATK v.3.3 [150]. This variant calling procedure was replicated using the *P. tetraurelia* strain 32 MAC genome assembly.

To build the genetic map, we considered only biallelic SNPs (excluding indels) with good quality scores that were called consistently on both genome assemblies. Given that F2’s underwent autogamy, they are expected to be entirely homozygous. We therefore excluded SNPs that were genotyped as heterozygous. We also excluded SNPs that showed strong deviations from Mendelian expectations (see details in Supplementary Methods). We retained 220,944 SNPs to build the genetic map. Two individuals (F2.11 and F2.12) were excluded because they derive from the same meiosis as F2.10. We also excluded one individual (F2.7) that appeared to result from an abnormal autogamy process (most of its scaffolds are heterozygous). In the end, our analysis of meiotic recombination events is based on 39 independent F2’s.

The strain 51 MIC reference genome consists of 187 supercontigs, including 164 supercontigs that match scaffolds of the MAC assembly. We identified three cases where two supercontigs encompassed contiguous segments of a same MAC scaffold, indicating that they derive from a same MIC chromosome. These pairs were grouped, resulting in a total of 161 supercontig groups matching MAC scaffolds. We excluded the 23 supercontigs that do not encompass any scaffold of the MAC assembly. The 161 supercontig groups that we selected to build the genetic map cover 101.4 Mb (97.3 % of the MIC reference genome assembly).

For each F2, we identified crossover (COs) and non-crossover (NCOs) recombination events (see details in Supplementary Methods). Because of the low number of callable SNPs in MAC-variable and MIC-limited regions (0.06 SNPs per kb), we had very limited power to detect recombination in these regions. The SNP density is also limited in IESs (0.27 SNPs per kb compared to 3.0 SNPs per kb in MAC-constitutive regions), but COs can nevertheless be detected in IESs thanks to the presence of SNPs in flanking MAC-constitutive regions. Thus, to quantify recombination rate, we restricted our analyses to MAC-constitutive regions and IESs, that cover 74 Mb (70.9% of the MIC genome) and include 99.3% of all detected COs.

We quantified CO interference using the coefficient of coincidence (CoC), as described in [151] (see details in Supplementary Methods).

### Collapse estimation

Based on the coverage distribution shown in SupFigure12, the genomic regions with coverage above the peak were extracted, then the coverage was divided by the median coverage to obtain a collapse score. The estimate of global genome collapse was calculated by multiplying the size of the regions by the collapse scores.

### Assembly Completeness

Kmer Analysis Toolkit v. 2.4.2 [54] was used to evaluate the completeness of genome assemblies, using K-mer frequency histograms (k = 27) and vegetative MIC DNA-seq datasets for the 7 *P. aurelia* species.

### MIC Telomeres

To identify MIC telomere repeats, MIC Illumina paired-end reads improperly mapped to the corresponding MAC assembly were analyzed using Jellyfish [152] to count k-mers (k-mer lengths of 18, 21 and 24 nt) corresponding to 3 exact repeats of 6, 7 or 8 nucleotide microsatellite repeat, taking into account all permutations. Aside from repeats containing only A and T, the top abundance k-mers include 18-mers with 3 repeats of 5’-CCCCAA-3’ and of 5’-CCCAAA-3’ hexamers. For further analyses, we used the Perl regular expression (CCC(C|A)AA){3,} to identify telomere repeats: at least 3 occurrences of the same mixture of C_4_A_2_ and C_3_A_3_ hexamers found in *Paramecium* MAC DNA [76], starting with validation that perfect triplets of either 5’-CCCCAA-3’ or 5’-CCCAAA-3’ are restricted to the extremities of the *P. tetraurelia* EZL1 ONT supercontigs, unlike the highly abundant ((A|T){6}){3} k-mers, which are distributed all along supercontigs (SupFigure11 ).

To determine the junctions between MIC telomere repeats and adjacent sequence, we defined telomere reads to be *P. tetraurelia* MIC Illumina HiSeq 101 nt read pairs (N = 90703803) that contained, in at least one read, a match to at least 3 consecutive repeats of a mixture of GGGGTT and GGGTTT hexamers, in either orientation. Telomere read pairs (N = 29771; 0.0328% of total read pairs) were fused using FLASH v1.2.11 [137] with default parameters to yield 20861 extended fragments of mean size 163 nt. PCR duplicates (N=93) were removed to yield 20768 fragments of mean size 165 nt.

Telomere Repeat Clusters were generated by clustering telomere-repeat containing fragments with cd-hit-est version 4.8.1 (-c 0.99 -n 10 -G 0 -A 75 -g 1 -gap -8 -gap-ext -3) [153], after removing the telomere repeats to avoid spurious clustering. With these options, clustering in “accurate” mode is based on a local alignment requiring 99% nucleotide identity using a 10 nt word size, with gap and gap extension penalties slightly larger than default (default is -6 and -1, respectively), the alignment covering at least 75 nt for each sequence in the cluster. The complete sequences (i.e. with the telomere repeats) of each cluster were aligned using MAFFT (v7.40) (option - adjustdirectionaccurately) [154]. Heatmap overviews of the Telomere Repeat Cluster alignments were generated using a custom R script (SupData2). Alignment consensus sequences were mapped to genomes using Bowtie2 (v2.3.4.3 --very-sensitive-local) [155]. The mapping revealed some redundancy in the set of junctions, since some loci were identified by more than one telomere junction consensus sequence. This included cases where an internal junction and an edge junction mapped to the same location as well as cases where two clusters in antisense orientation mapped to the same location.

### Synteny between MAC and MIC sequences

Assemblies were masked using RepeatMasker (v-open-4.0.8 -no_is -nowlow -x -species paramecium). For each species, MAC and MIC sequences were grouped using RagTag (v1.0.2 ragtag.py scaffold -w -f 2000 -I 0.5 -u) [156]. All MIC/MAC sequence groups were manually curated, using Circos (v0.69-6, [157] to display the sequence alignments (nucmer v3.1 filtered with -i 0.99 -l 1000, [158]. The SupData1_synteny_MIC_MAC.html file shows the graphics used to curate the comparison between the *P. tetraurelia* EZL1 ONT assembly and the MAC assembly.

### Genomic compartments

IESs from [14,25] were remapped on MIC genomes using a custom Perl wrapper (remap_IES.pl, see Zenodo) and Bowtie2 [155] (v2.3.4.3 --very-sensitive) alignments guided by the groups based on synteny between MAC and MIC sequences. As previously documented [26] we characterized “buried” IESs on MIC-limited regions using ParTIES (MILORD method, [78]).

We determined non-overlapping genomic compartments: MAC-constitutive, MAC-variable, Other Eliminated Regions (OES, eliminated regions apart from IESs). For this purpose, we used MAC DNA and MIC DNA bigwig (bin=1) sequencing coverage files. For each dataset, the threshold was adjusted (see SupFigure12_Compartments). For each MIC genome, the MAC-constitutive compartment was defined by regions with a MAC coverage ≥ ∼100X after fusing contiguous regions (R GenomicRanges package, reduce method, min.gapwidth=100). Previously annotated IESs were excluded. Remaining regions selected for a MAC coverage ≥ 5X, were fused as previously described (min.gapwidth=100nt), then filtered for regions longer than 100nt defining the MAC-variable compartment. The OES compartment was defined by remaining regions with a MIC DNA coverage ≥ 5X, fused (min.gapwidth=100nt), then filtered for regions longer than 100nt. SupFigure12_Compartments shows the proportion of each compartment for each MIC genome. The OES compartment for the *EZL1* ONT assembly was divided into two sub-compartments: *PGM*- dependent and *PGM*-independent, using the same threshold strategy (≥ 5X, then fused at min.gapwidth=100 and filtered for segments with length ≥ 100nt) on *PGM* DNA.

### Gene annotation

Previously annotated genes on MAC genomes [11,53] were remapped to MIC genomes using a custom Perl script (remap_gene.pl, see Zenodo) that uses refined BLAT (v36x2 -fine -maxIntron=500) [159] alignments guided by MAC/MIC sequence groups (see previous section). On each MIC genome the MAC-destined compartment (MAC-constitutive and -variable; see previous section) was masked using bedtools (v2.26.0 maskfasta). The remaining regions (MIC-limited regions) were annotated for genes using EuGene (v4.1) software and the procedure previously described [53]. Only genes with a length ≥ 100nt and G+C content < 50% were kept.

### Transposable Element landscape

The TEdenovo module of REPET version 2.5 [160] was used for identification of Transposable Elements (TE) for each Illumina assembly, after adding all available curated Paramecium TE [16] to Repbase [161] v19.06. Default TEdenovo parameters were used (except for BLR_sensitivity : 0; TRFmaxPeriod : 500) and the MIC-specific (masked constitutive MAC) assembly was used. TEannot was run with the same parameters on the unmasked MIC assembly, and the annotations were used for human curation of the reference TEs proposed by TEdenovo. This led to discovery of Helitrons in *Paramecium* genomes. For the subsequent round of annotation, manually curated Helitron ORFs and the reference TEs for all species were pooled and used to annotate each assembly, including the *P. tetraurelia* EZL1 long-read assembly, with TEannot. Coverage of each assembly by different TE superfamilies was calculated using custom R scripts and the Genomic Ranges package (code and data in Zenodo).

### TE protein domains and phylogeny

To identify TE independently of REPET, we searched for 24 PFAM domains curated from TEs frequently found in fungi [162] in the MIC Illumina assemblies. The constitutive MAC was masked using RepeatMasker and the genomes were translated in 6 reading frames using the transeq program of the EMBOSS suite (http://emboss.open-bio.org/). HMMER 3.2. (http://hmmer.org) hmmsearch was used to search for the TE domains in the translated genome assemblies using default parameters. Using a cutoff domain e-value of 0.0001 to obtain only some of the least decayed TE copies, the nucleotide sequences corresponding to each hit for the genomes under consideration were extracted then aligned with MACSE v. 2.07 [163,164]. SeaView version 5.0.4 [165] was used to build a phylogenetic tree with phyML [166] (default parameters) from the MACSE alignment for each PFAM domain. Annotation and visualization of trees was enhanced using the R ggtree package [167,168].

### dN/dS computation

The phylogenetic tree of full size Helitron transposase copies was computed with phyML [166], based on the protein sequence alignment obtained with MAFFT using the L-INS-i method [154]. We used bio++ v.3.0.0 libraries [169–171] to estimate the dN/dS on each branch of the phylogenetic tree. In a first step, we used a homogeneous codon model implemented in *bppml* to infer the most likely branch lengths, codon frequencies at the root, and parameters of the YN98 (F3X4) substitution model [172], which allows for different nucleotide content dynamics across codon positions. In a second step, we used the *MapNH* substitution mapping method [173] to count synonymous and non-synonymous substitutions along each branch [174].

### Minisatellite identification

Our strategy was to use the sensitive MREPS software to identify tandem repeats in assemblies, and TAREAN to identify tandem repeats in sequencing reads. MREPS [175] v2.6 (-minsize 300 -minperiod 30 -exp 6 -res 150) was used to find minisatellite DNA with a repeat period of at least 30 nt in genome assemblies. Custom R scripts were used to align and build a consensus from the repeats at each locus. The consensus sequences were clustered using cd-hit-est (version 4.8.1) [153], with a first set of parameters for small repeats (-c 0.8 -n 5 -A 100 -g 1 -G 0 -sc 1 -r 0 -d 0) and a second set for longer repeats (-c 0.9 -n 9 -A 30 -g 1 -G 0 -sc 1 -r 0 -d 0).

To identify minisatellite using sequencing reads, TAREAN software [176] was used with the default parameters for paired-end Illumina reads. For each species, reads were trimmed with sickle software [177] at quality -q 20, and trimmed reads were sampled to obtain reads that correspond to ∼ 0.5X to ∼1X estimated coverage of the genome. Six independent samples of reads were run for each species. The consensus sequences for the six runs were compared by blastn to determine a non-redundant set of consensus sequences for each species. The non-redundant sets of consensus sequences (for both MREPS and TAREAN workflows) were mapped to assemblies using RepeatMasker version open-4.0.8 (-s -gccalc -no_is -nolow) [178]. One TAREAN limitation is the requirement of a certain abundance of a repeat in the genome for detection.

### G-quadruplex identification

The R package pqsfinder version 2.18.0 (-deep=TRUE and a min score of 70) [179] was used to computationally identify potential quadruplex-forming sequences in an assembly.

### Dotplot comparisons

Dotplots were used for fine-grained visualization of tandem repeats that were too degenerate or too small for detection by the automated minisatellite pipelines. For dotplots of sequences of the same length, a custom R script was used to plot a distance matrix using the DistanceMatrix method of DECIPHER (v2.28.0; window 28, Hamming distance threshold 8) and ggplot2_v3.5.1. For non-self dotplots of Helitrons, the EMBOSS v6.6.0.0 [180] dotmatcher function was used (-graph data - windowsize 28 -threshold 54 or -threshold 60) to generate the matrix, which was drawn using R version 4.3.0 graphics::plot.

### Software

mRNA-seq data were mapped using HISAT2 (v2.1.0 --min-intronlen 20 --max-intronlen 100) [181]. DNA-seq data were mapped using BOWTIE2 (v2.2.9 --local ) [155]. Small RNA-seq data were mapped using BWA (aln v0.7.15 0 no mismatch) [182]. Read counts on sequence features (genes,TEs, minisatellite) were calculated with htseq-count (v0.11.2, -stranded=yes) [183] on mappings after quality filtering with SAMtools (v1.9, q>=30 for data mapped on *Illumina* assemblies and q>=0 data mapped on ONT assemblies) [148]. Data were analyzed and displayed using R (v4.3.0) statistical computing framework [184] and R packages (Biostrings-v2.70.1 [185], ComplexHeatmap-v2.18.0, DECIPHER-v2.28.0 [186], GenomicRanges-v1.54.1 [187], ggdist-v3.3.1 [188], ggtree-v3.10.0 [167], karyoploteR-v1.28.0 [189], magrittr-v2.0.3, rtracklayer-v1.62.0 [190], seqinr-v4.2-36 [191], tidyverse-v2.0.0 [192], UpSetR-v1.4.0 [193]).

## Data availability

Sequencing data are available in the European Nucleotide Archive under Project PRJEB98202 (https://www.ebi.ac.uk/ena/browser/view/PRJEB98202) and Project PRJEB98215 (https://www.ebi.ac.uk/ena/browser/view/PRJEB98215) and are described in the SupTables 1 & 2. The statistical data and scripts (bash, R or perl) used to generate results and images were deposited at Zenodo (https://doi.org/10.5281/zenodo.17205922). The SupData1 – SupData4 mentioned in this article can be downloaded from Zenodo (https://doi.org/10.5281/zenodo.17206076). All genome and annotation files are available from the ParameciumDB [194] download section (https://paramecium.i2bc.paris-saclay.fr/download/).

## Abbreviations

CHBD0: Chessboard 0 minisatellite
CO: Crossover
DCL: Dicer-like
DEV: Development
EZL1: Enhancer of Zest-like 1
FRAG: Fragmentation
HLE: Helitron-like Element
IES: Internal Eliminated Sequence
LINE: Long Interspersed Nuclear Element
LTR: Long Terminal Repeat
LTS: Left Terminal Sequence
MAC: Macronucleus
MDS: MAC-destined sequence
MIC: Micronucleus
MLS: MIC-limited Sequence
MITE: Miniature Inverted Repeat Transposable Element
NCO: Non-Crossover
OES: Other Eliminated Sequence
ONT: Oxford Nanopore Technology
PDE: Programmed DNA Elimination
PGM: PiggyMac
PRC2: Polycomb Repressive Complex 2
RPKM: Reads per kilobase per million mapped reads
RTS: Right Terminal Sequence
SINE: Short Interspersed Nuclear Element
SNP: Single Nucleotide Polymorphism
TE: Transposable Element
TFIIS4: transcription elongation factor II paralog 4
TIR: Terminal inverted repeat DNA transposon
WGD: Whole Genome Duplication

## ACKNOWLEDGEMENTS

We acknowledge use of the computing facilities of the CC LBBE/PRABI and the BIOI2 platform and computing facilities of the I2BC. We thank Pascaline Tirand, Fanny Culot, Nelly Sainsard, Cristina Delawarde and Vincent Maupu-Massamba for technical assistance. We dedicate this article to our dearly missed colleagues Janine Beisson and Sophie Malinsky, who played essential roles in the international ciliate research community and in our French *Paramecium* laboratories.

## Author contributions

OA played a major role in data curation, formal analysis, software development, and preparation of the manuscript. GP, IN and AP performed genetic crosses and sample preparation for the genetic linkage map. AC, JMA and KL were involved in aquistion and formal analysis of the sequence data. JA, IL and HQ played a major role in annotation, formal analysis and interpretation of repeated sequences. FG, CMP, SB, AF, SM, NM, NS, VR, AT, AdV and CZ performed RNAi and flow cytometry experiments and prepared samples for sequencing. EL annotated and interpreted repeated sequence data. MB, LD, EM, SD and LS were responsible for conception, supervision and funding of the project. OA, MB, LD, EM, SD and LS wrote and revised the manuscript, and contributed to data visualisation. LD, EM and LS performed formal analysis, annotation and data curation. KL and SD coordinated the project. All living authors have approved the manuscript.

## Supplementary Information

### HTML files

SupData directories contain an html file for interactive exploration of supplementary data, the Rmd script and data used to generate the html file, and in some cases, embedded images. The html file can be opened with a web browser (JavaScript should be enabled in the browser Preferences).

SupData1_Synteny_MIC_MAC

### Synteny between *P. tetraurelia* supercontigs of the MIC *EZL1*-RNAi assembly and scaffolds of the MAC v2 assembly

In the Circos drawings, from the center to the periphery, the arcs (red, sequence and query in the same orientation; blue, sequence and query in opposite orientations) show the syntenic regions identified by nucmer (v3.1, filtered with delta-filter -m -i 0.99 -l 1000). The darker the color the greater the sequence identity. The curves give coding gene density (orange), TE density (blue), and satellite (tandem repeat) density (green). The histograms show *PGM-*RNAi coverage (blue), MAC coverage (purple) and MIC coverage (red). The interactive table summarizes the data and allows the user to customize table appearance, sort by column, download a csv or excel file and search for a supercontig or scaffold. It is possible to scroll through the images without displacing the table and to jump from any image back to the table. This data was used to prepare Figure 2D.

SupData2_TelomereJunctions

### Telomere Repeat Cluster overviews

The interactive table describes intrinsic characteristics of each telomere repeat cluster, its mapping to different genome assemblies and provides a link to the overview images, which can be scrolled independently of the table. This data was used to prepare Figure 2E & F.

SupData3_Poster

### Exhaustive overview of supercontigs and their annotations

The 187 supercontigs of the *P. tetraurelia EZL1*-RNAi genome assembly were drawn to scale with annotations of telomere repeats, satellites, curated Helitron ORFs, curated HLE start and end (LTS and RTS) consensus sequences, REPET automated annotations of LINE and TIR superfamily elements and IESs (detailed legend at the top of the html file). The drawings also show MAC coverage and genome compartments. The interactive table summarizes information about genome compartments and provides links to the drawings.

SupData4_EndGame

### Zoom of supercontig ends

The 50 kb at the ends of each of the 187 supercontigs of the *P. tetraurelia EZL1*-RNAi ONT genome assembly are drawn to show annotations of telomere repeats, satellites, curated Helitron ORFs, Helitron start and end consensus sequences (LTS, RTS), REPET automated annotations of LINE and TIR superfamily elements and IES (detailed legend in the html file). The drawings also show MAC coverage and genome compartments. The interactive table summarizes the information and provides links to the drawings. Each drawing is accompanied by a link to a dotplot of the first 20 kb of the end with annotation of minisatellites.

