## SupplementalFigures for "The tiny germline chromosomes of *Paramecium aurelia* have an exceptionally high recombination rate and are capped by a new class of Helitrons"

SupFigure 1

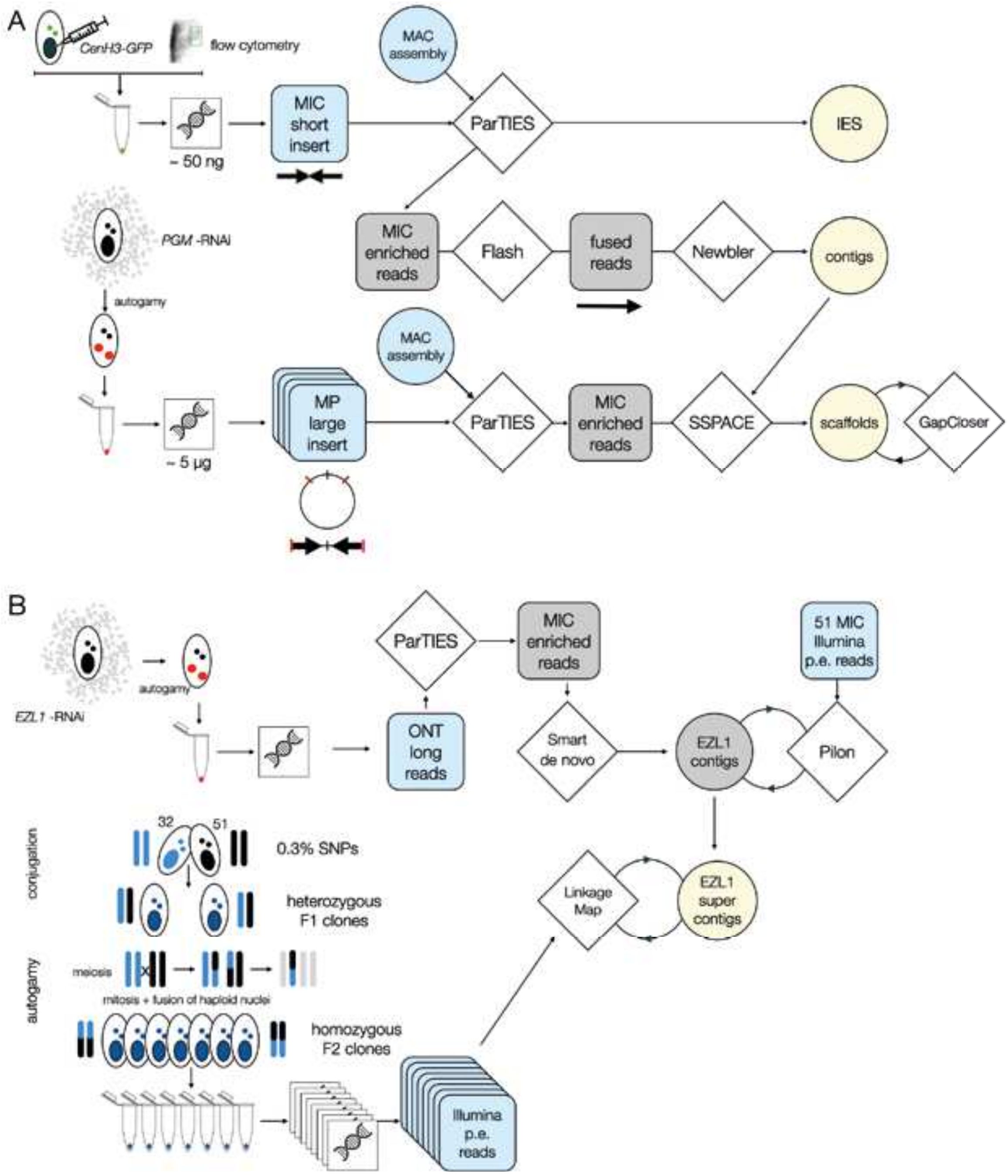


**SupFigure1 Assembly workflows**

(A) Illumina assemblies were made with vegetative MICs obtained by fluorescence-activated sorting of nuclei from cells expressing GFP-tagged CenH3a, a MIC-specific marker [16,34]. DNA was extracted from the sorted MICs and Nextera librairies constructed for paired-end sequencing. Illumina paired-end reads were enriched in MIC DNA by removing any read pairs with a MAC IES junction in one mate. Contigs were assembled with Newbler, scaffolded with SSpace and PGM-RNAi mate-pair librairies, and gap-filled with GapCloser (SoapDeNovo v2) as detailed in Methods. (B) For P tetraurelia, Oxford Nanopore Technology long reads were obtained using DNA from new MACs that developed in cells depleted of Ezl1, the catalytic subunit of PRC2 [34,45,46] required for Programmed DNA Elimination and for transposon silencing. Long reads that originated from fragments of the old MAC as judged by presence of MAC IES junctions were removed. The long reads were assembled with SMARTdenovo and the contigs were polished using Pilon and Illumina reads (Methods). A recombination map was made using F2 clones of a P tetraurelia stock 32 x P tetraurelia stock 51 cross (cf. Methods and SupFigure8). Information from the recombination map was used to break chimeras and build supercontigs.

SupFigure 2

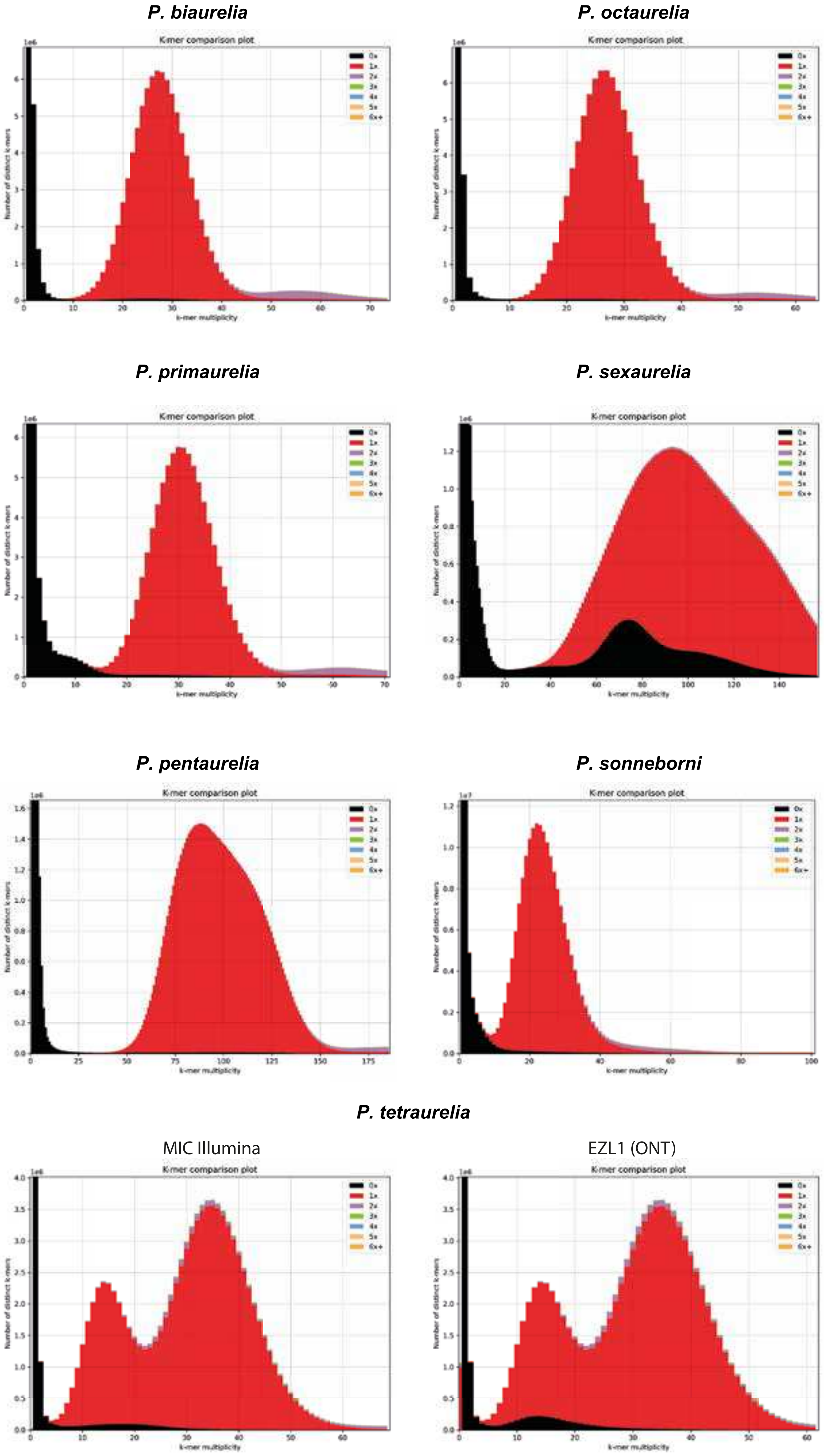


**Assembly spectra copy number plots**
Prepared using the Kmer Analysis Toolkit v. 2.4.2; [54], these K-mer frequency histograms show the K-mer content from MIC Illumina reads found in the corresponding P. aurelia genome reference assemblies. In red, k-mers found once in the assembly. The k-mers in black mostly represent sequencing errors that did not make it into the assembly. The 1X Kmers (red) overwhelmingly predominate in each of the plots, providing a strong argument that the assemblies are complete. The sequenced DNA comes from 100% homozygous populations, as expected for a clonal line expanded after autogamy (self-fertilization), thus small amounts of 2X (mauve) coverage likely arises not because of heterozygosity but rather because paralogous genes share some identical k-mers. Note that the fact that there are two main peaks, in particular for P. tetraurelia, stems from MAC contamination of the sorted vegetative MICs [16,195].

SupFigure 3

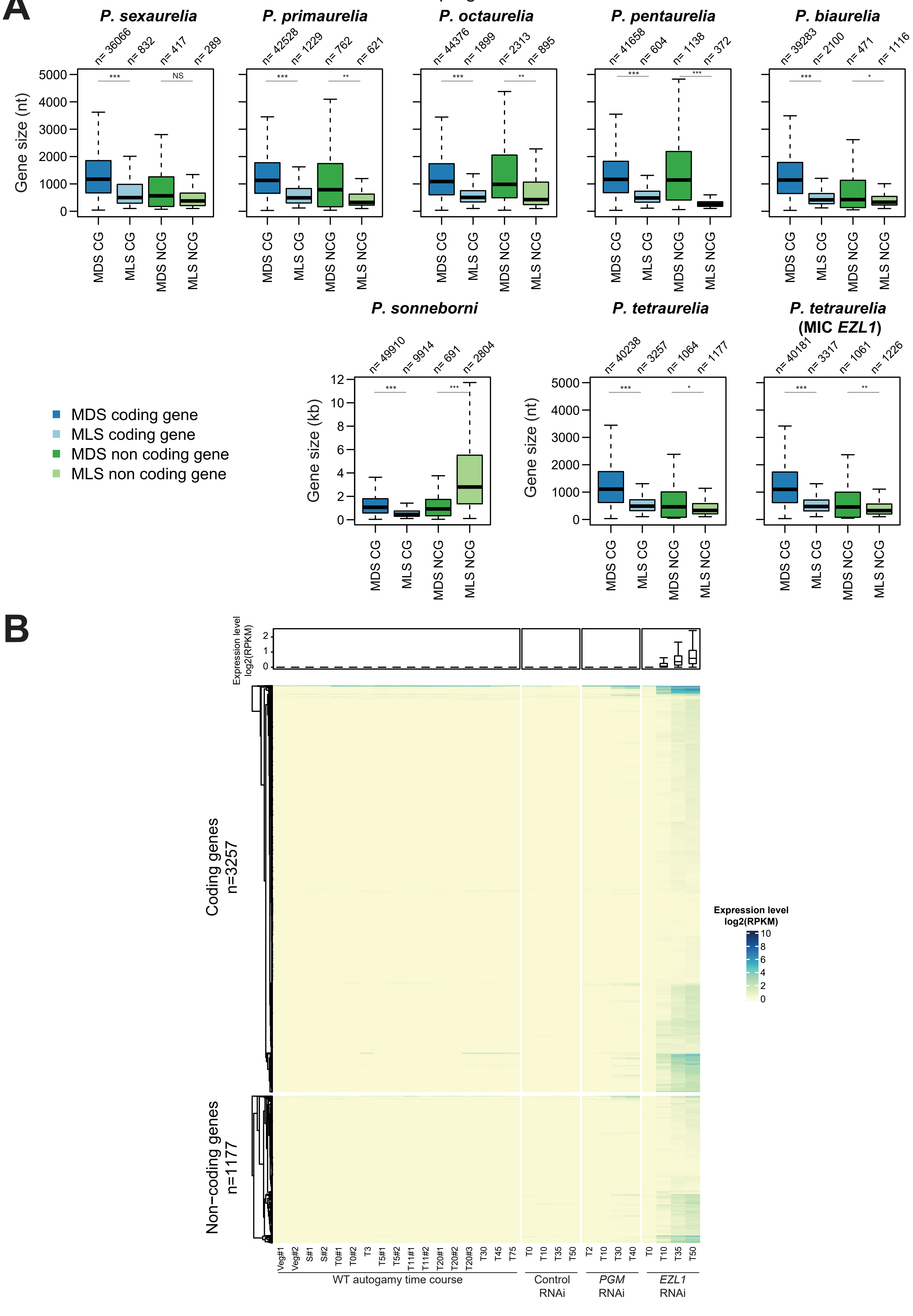
A
P. sexaurelia
P. primaurelia
P. octaurelia
P. pentaurelia
P. biaurelia
n= 36066
n= 832
n= 417
n= 289
n= 42528
n= 1229
n= 762
n= 621
n= 44376
n= 1899
n= 2313
n= 895
n= 41658
n= 604
n= 1138
n= 372
n= 39283
n= 2100
n= 471
n= 1116
Gene size (nt)
***
NS
**
*
MDS CG
MLS CG
MDS NCG
MLS NCG
P. sonneborni
P. tetraurelia
P. tetraurelia (MIC EZL1)
n= 49910
n= 9914
n= 691
n= 2804
n= 40238
n= 3257
n= 1064
n= 1177
n= 40181
n= 3317
n= 1061
n= 1226
Gene size (kb)
MDS coding gene
MLS coding gene
MDS non coding gene
MLS non coding gene
B
Expression level log2(RPKM)
Coding genes n=3257
Non-coding genes n=1177
Veg#1
Veg#2
S#1
S#2
T0#1
T0#2
T3
T5#1
T5#2
T11#1
T11#2
T20#1
T20#2
T20#3
T30
T45
T75
T0
T10
T35
T50
T2
T10
T30
T40
T0
T10
T35
T50
WT autogamy time course
Control RNAi
PGM RNAi
EZL1 RNAi

**SupFigure3 Size and expression of putative MIC-limited genes**

(A). For MIC-limited sequences (MLS), putative gene size distributions were compared to those for published annotation of MAC-destined sequences (MDS) [11,53]. The boxplots (sample size indicated on top of each box) show the size distributions of protein-coding (CG) and non-coding (NCG) genes in MLS and MDS. P-values were calculated using a t-test, and the statistical significance is indicated as follows: ***<1e-50, **<1e-10, *<1e-2 or NS (non-significant). The smaller size of the MLS distributions, especially for coding genes, suggests that the MLS gene annotations contain pseudogenes and false positives. Note that the MLS NCG in P. sonneborni are much longer than in the other species, probably corresponding to pseudogenes subsequent to recent HT from other aurelia genomes (see [55]). (B) Heatmap showing the expression level (log2 RPKM) of MLS coding and non-coding genes. mRNA-Seq samples from wild-type autogamy time courses (Arnaiz et al 2017) and four autogamy time points from 3 silencing conditions (Control, PGM and EZL1 RNAi) [46,140]. Boxplots above the heatmap summarize the expression levels of these genes.

SupFigure 4

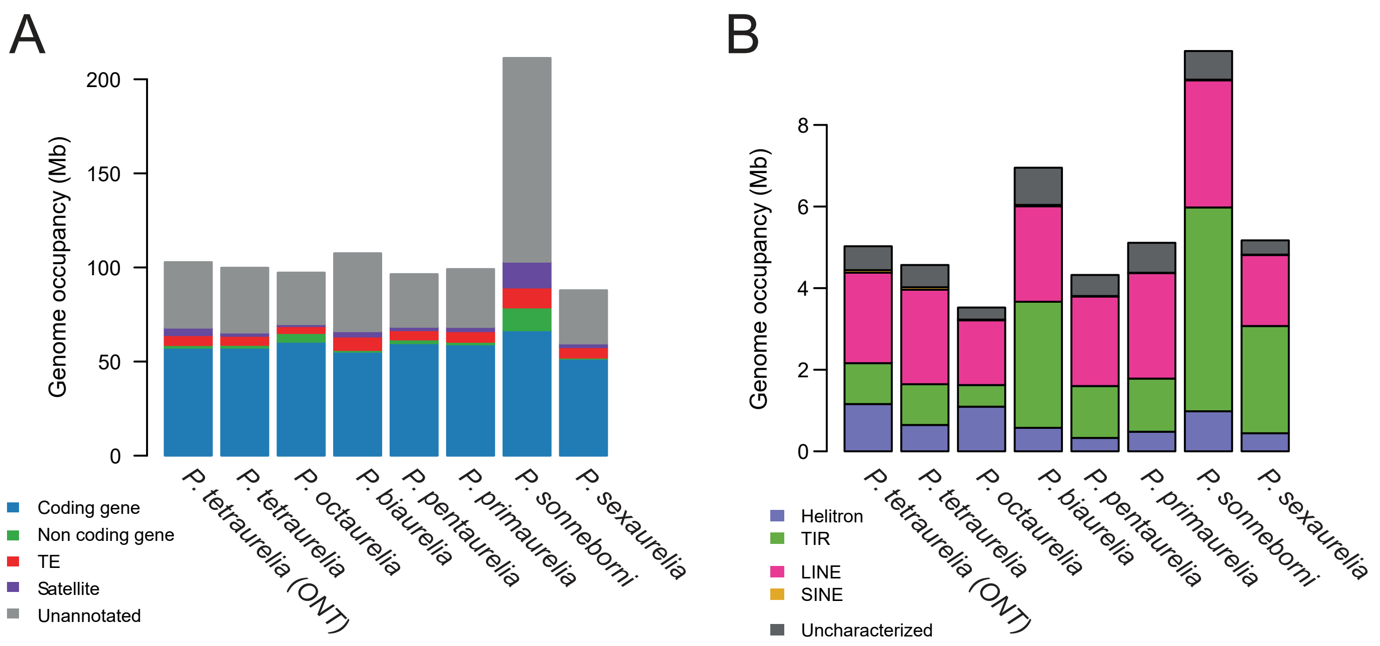


**SupFigure4  Occupancy of MIC genome assemblies by sequence features.**

(A) Bar plots showing the genome occupancy of MIC genome annotations in Mb for each assembly. (B) TEs annotated using REPET were classified as Helitrons, TIRs, LINEs, SINEs and uncharacterized elements. Bar plots show their MIC genome occupancy in Mb.

**SupFigure5 (below)**

(A) Visualization of the 15 most conserved minisatellites in P. aurelia genomes, after mapping the libraries of minisatellite DNA consensus units to MIC genomes of each species using RepeatMasker. These satellites correspond to the tops of the heatmaps in Figure 1C. Each row corresponds to a minisatellite and each column to a target species. If there were hits, the intensity of the blue rectangle is proportional to the similarity of the best hit to the target genome. (B) Heatmap ordered by abundance of minisatellites identified in the P. tetraurelia MIC EZL1 assembly. Labels to the right give the size of the repeat unit (nt) followed by the name of the minisatellite, names to the left correspond to IDs of minisatellites referred to in Results and Figure

SupFigure 5

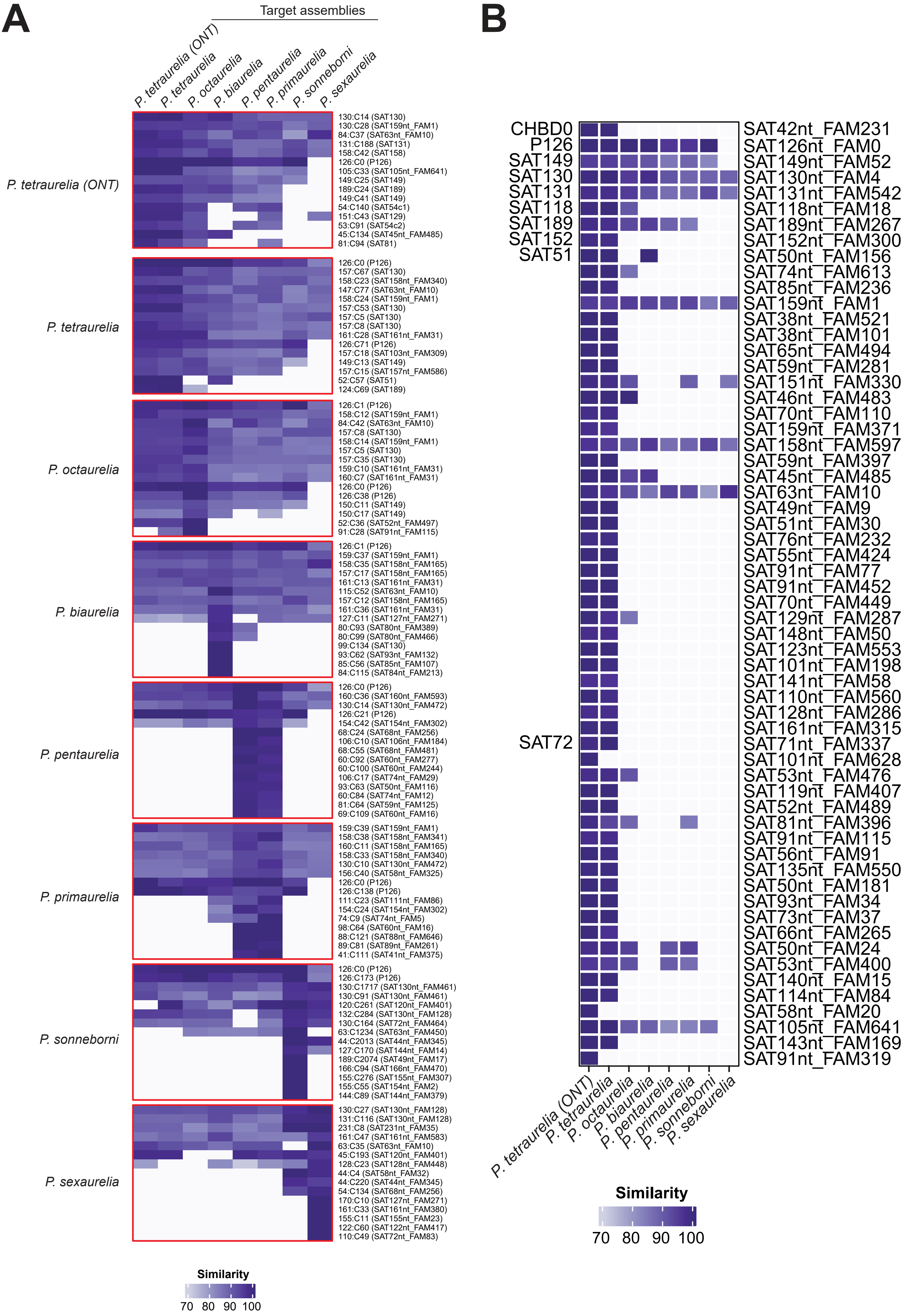
A
Target assemblies
P. tetraurelia (ONT)
P. tetraurelia
P. octaurelia
P. biaurelia
P. pentaurelia
P. primaurelia
P. sonneborni
P. sexaurelia
P. tetraurelia (ONT)
130:C14 (SAT130)
130:C28 (SAT159nt_FAM1)
84:C37 (SAT63nt_FAM10)
131:C188 (SAT131)
158:C42 (SAT158)
126:C0 (P126)
105:C33 (SAT105nt_FAM641)
149:C25 (SAT149)
189:C24 (SAT189)
149:C41 (SAT149)
54:C140 (SAT54c1)
151:C43 (SAT129)
53:C91 (SAT54c2)
45:C134 (SAT45nt_FAM485)
81:C94 (SAT81)
P. tetraurelia
126:C0 (P126)
157:C67 (SAT130)
158:C23 (SAT158nt_FAM340)
147:C77 (SAT63nt_FAM10)
158:C24 (SAT159nt_FAM1)
157:C53 (SAT130)
157:C5 (SAT130)
157:C8 (SAT130)
161:C28 (SAT161nt_FAM31)
126:C71 (P126)
157:C18 (SAT103nt_FAM309)
149:C13 (SAT149)
157:C15 (SAT157nt_FAM586)
52:C57 (SAT51)
124:C69 (SAT189)
P. octaurelia
126:C1 (P126)
158:C12 (SAT159nt_FAM1)
84:C42 (SAT63nt_FAM10)
157:C8 (SAT130)
158:C14 (SAT159nt_FAM1)
157:C5 (SAT130)
157:C35 (SAT130)
159:C10 (SAT161nt_FAM31)
160:C7 (SAT161nt_FAM31)
126:C0 (P126)
126:C38 (P126)
150:C11 (SAT149)
150:C17 (SAT149)
52:C36 (SAT52nt_FAM497)
91:C28 (SAT91nt_FAM115)
P. biaurelia
126:C1 (P126)
159:C37 (SAT159nt_FAM1)
158:C35 (SAT158nt_FAM165)
157:C17 (SAT158nt_FAM165)
161:C13 (SAT161nt_FAM31)
115:C52 (SAT63nt_FAM10)
157:C12 (SAT158nt_FAM165)
161:C36 (SAT161nt_FAM31)
127:C11 (SAT127nt_FAM271)
80:C93 (SAT80nt_FAM389)
80:C99 (SAT80nt_FAM466)
99:C134 (SAT130)
93:C62 (SAT93nt_FAM132)
85:C56 (SAT85nt_FAM107)
84:C115 (SAT84nt_FAM213)
P. pentaurelia
126:C0 (P126)
160:C36 (SAT160nt_FAM593)
130:C14 (SAT130nt_FAM472)
126:C21 (P126)
154:C42 (SAT154nt_FAM302)
68:C24 (SAT68nt_FAM256)
106:C10 (SAT106nt_FAM184)
68:C55 (SAT68nt_FAM481)
60:C92 (SAT60nt_FAM277)
60:C100 (SAT60nt_FAM244)
106:C17 (SAT74nt_FAM29)
93:C63 (SAT50nt_FAM116)
60:C84 (SAT74nt_FAM12)
81:C64 (SAT59nt_FAM125)
69:C109 (SAT60nt_FAM16)
P. primaurelia
159:C39 (SAT159nt_FAM1)
158:C38 (SAT158nt_FAM341)
160:C11 (SAT158nt_FAM165)
158:C33 (SAT158nt_FAM340)
130:C10 (SAT130nt_FAM472)
156:C40 (SAT58nt_FAM325)
126:C0 (P126)
126:C138 (P126)
111:C23 (SAT111nt_FAM86)
154:C24 (SAT154nt_FAM302)
74:C9 (SAT74nt_FAM5)
98:C64 (SAT60nt_FAM16)
88:C121 (SAT88nt_FAM646)
89:C81 (SAT89nt_FAM261)
41:C111 (SAT41nt_FAM375)
P. sonneborni
126:C0 (P126)
126:C173 (P126)
130:C1717 (SAT130nt_FAM461)
130:C91 (SAT130nt_FAM461)
120:C261 (SAT120nt_FAM401)
132:C284 (SAT130nt_FAM128)
130:C164 (SAT72nt_FAM464)
63:C1234 (SAT63nt_FAM450)
44:C2013 (SAT44nt_FAM345)
127:C170 (SAT144nt_FAM14)
189:C2074 (SAT49nt_FAM17)
166:C94 (SAT166nt_FAM470)
155:C276 (SAT155nt_FAM307)
155:C55 (SAT154nt_FAM2)
144:C89 (SAT144nt_FAM379)
P. sexaurelia
130:C27 (SAT130nt_FAM128)
131:C116 (SAT130nt_FAM128)
231:C8 (SAT231nt_FAM35)
161:C47 (SAT161nt_FAM583)
63:C35 (SAT63nt_FAM10)
45:C193 (SAT120nt_FAM401)
128:C23 (SAT128nt_FAM448)
44:C4 (SAT58nt_FAM32)
44:C220 (SAT44nt_FAM345)
54:C134 (SAT68nt_FAM256)
170:C10 (SAT127nt_FAM271)
161:C33 (SAT161nt_FAM380)
155:C11 (SAT155nt_FAM23)
122:C60 (SAT122nt_FAM417)
110:C49 (SAT72nt_FAM83)
Similarity
70 80 90 100
B
CHBD0
P126
SAT149
SAT130
SAT131
SAT118
SAT189
SAT152
SAT51
SAT72
SAT42nt_FAM231
SAT126nt_FAM0
SAT149nt_FAM52
SAT130nt_FAM4
SAT131nt_FAM542
SAT118nt_FAM18
SAT189nt_FAM267
SAT152nt_FAM300
SAT50nt_FAM156
SAT74nt_FAM613
SAT85nt_FAM236
SAT159nt_FAM1
SAT38nt_FAM521
SAT38nt_FAM101
SAT65nt_FAM494
SAT59nt_FAM281
SAT151nt_FAM330
SAT46nt_FAM483
SAT70nt_FAM110
SAT159nt_FAM371
SAT158nt_FAM597
SAT59nt_FAM397
SAT45nt_FAM485
SAT63nt_FAM10
SAT49nt_FAM9
SAT51nt_FAM30
SAT76nt_FAM232
SAT55nt_FAM424
SAT91nt_FAM77
SAT91nt_FAM452
SAT70nt_FAM449
SAT129nt_FAM287
SAT148nt_FAM50
SAT123nt_FAM553
SAT101nt_FAM198
SAT141nt_FAM58
SAT110nt_FAM560
SAT128nt_FAM286
SAT161nt_FAM315
SAT71nt_FAM337
SAT101nt_FAM628
SAT53nt_FAM476
SAT119nt_FAM407
SAT52nt_FAM489
SAT81nt_FAM396
SAT91nt_FAM115
SAT56nt_FAM91
SAT135nt_FAM550
SAT50nt_FAM181
SAT93nt_FAM34
SAT73nt_FAM37
SAT66nt_FAM265
SAT50nt_FAM24
SAT53nt_FAM400
SAT140nt_FAM15
SAT114nt_FAM84
SAT58nt_FAM20
SAT105nt_FAM641
SAT143nt_FAM169
SAT91nt_FAM319
P. tetraurelia (ONT)
P. tetraurelia
P. octaurelia
P. biaurelia
P. pentaurelia
P. primaurelia
P. sonneborni
P. sexaurelia
Similarity
70 80 90 100

SupFigure 6

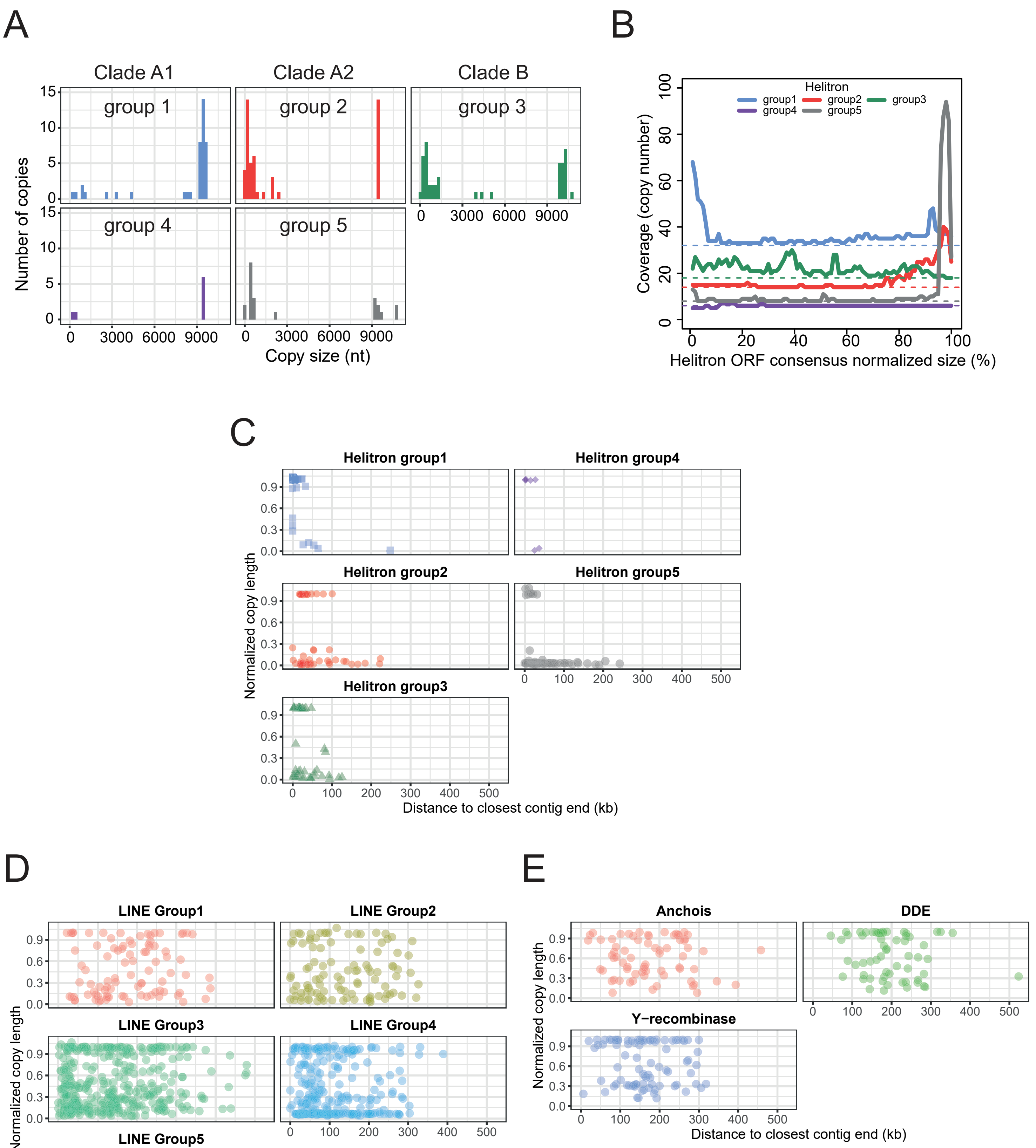


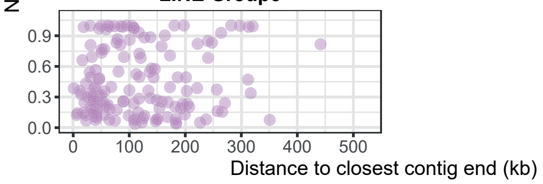


**SupFigure6 Helitron HelRep ORF size distribution, decay and distance to nearest contig end (P. tetraurelia EZL1 assembly)**

(A) Histogram of Helitron HelRep transposase ORF size by group (as defined by the phylogeny, SupFigure7). (B) To see which part of consensus sequences, representing the ancestral Helitrons, are still present in extant copies, we calculated the distribution of all copies along the Helitron consensus sequences grouped according to protein phylogeny. The colored dashed lines indicate the number of copies that cover at least 85% of the best matching consensus for each group. Additional coverage reveals a bias of decayed copies towards one or the other end of the ORF, especially for group5 but also group1 and group2. There are many short, decayed copies for group3 Helitron ORFs, but they are evenly distributed. (C-E) Relative size of Helitron (C), LINE (D), and Tc1/Mariner (E) transposase ORFs (fraction of complete ORF size) as a function of distance from supercontig ends.

### SupFigure 7

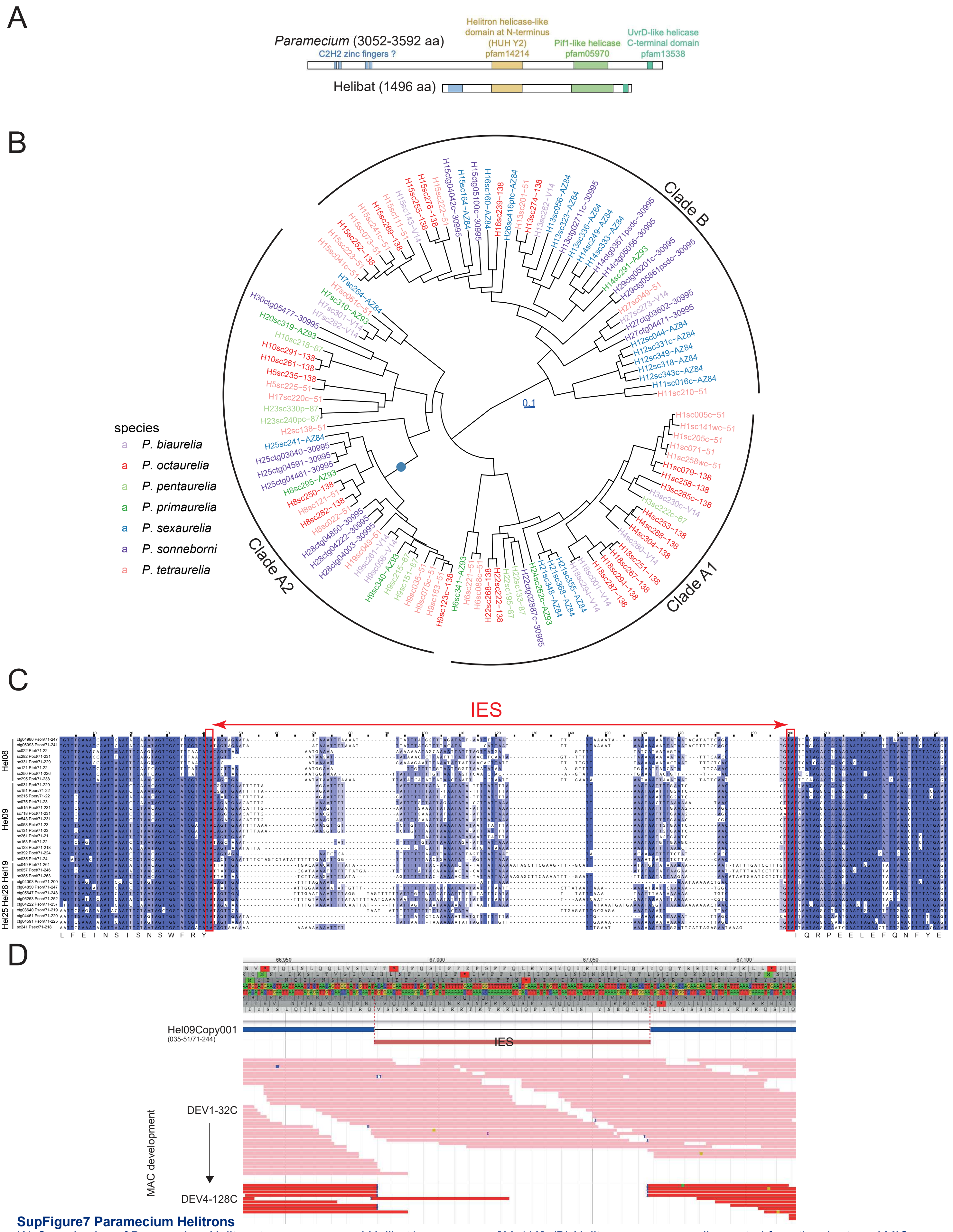


**SupFigure7 Paramecium Helitrons**

(A) Organization of Paramecium Helitron transposases and Helibat1 transposase [63,110]. (B) Helitrons were manually curated from the short read MIC assemblies, involving arbitrary correction of rare indels and in-phase stop codons ("c" suffix) to restore ORFs, align translation with MAFFT and construct a phyML tree [166]. The presence of an IES, at the same position, in the ORFs of one branch of the A2 clade, is marked by a blue dot. (C) Multiple alignment of Helitron ORF loci from many Paramecium species showing the sequences flanking the IES excision site. The red rectangles indicate the TA used for IES excision. The conceptual consensus translation figures below the alignment. (D) ParameciumDB genome browser screenshot of Hel09Copy001, containing a conserved IES presented in (B), confirming that this IES is correctly excised. The ORF is in blue and the conserved IES in red. Mapped reads from time-course samples at two MAC developmental stages (DEV1-32C, pink; DEV4-128C, red) show that the IES is excised between 32C and 128C endoreplication timing before the Helitron itself is completely eliminated.

SupFigure 8

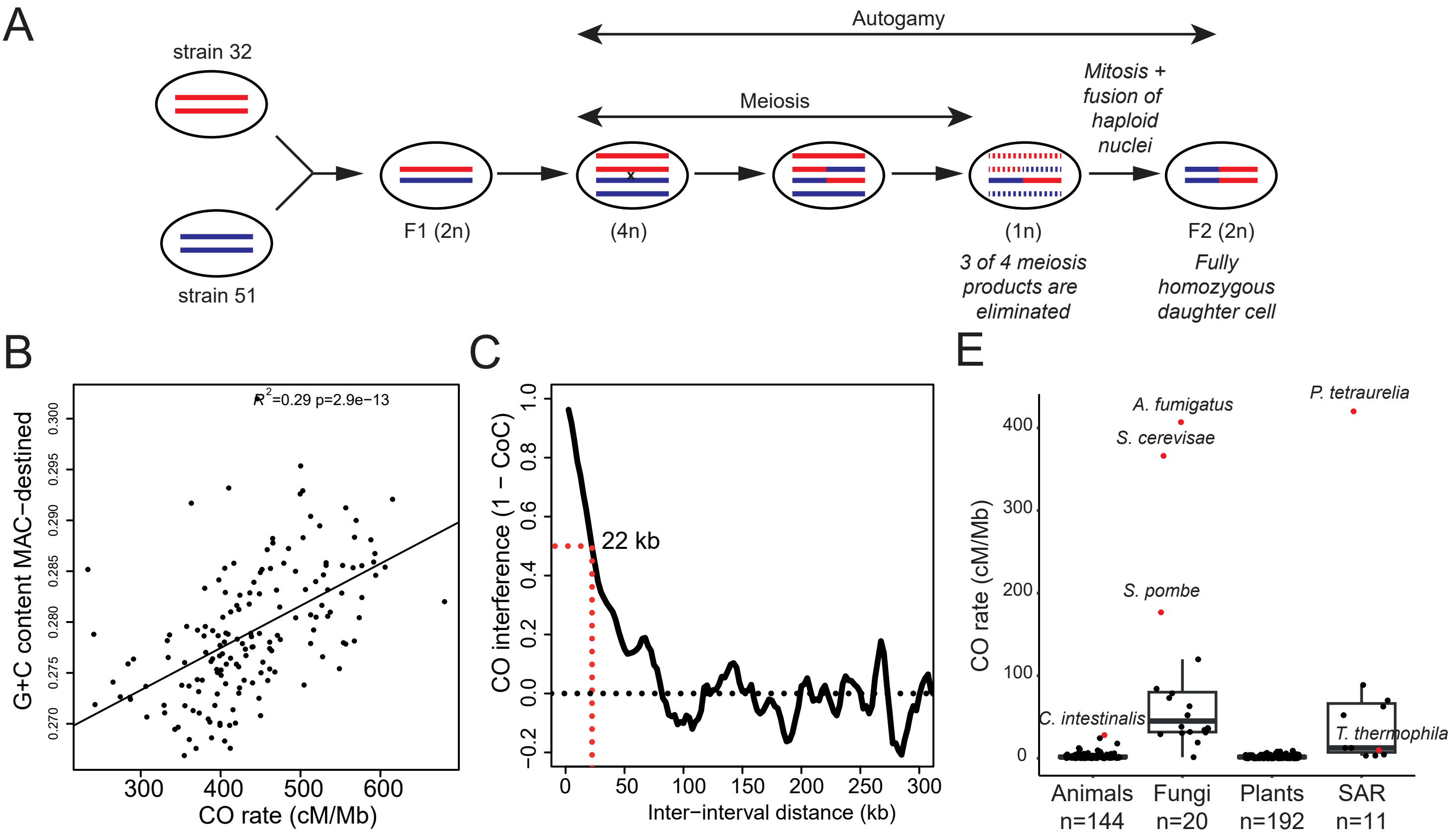


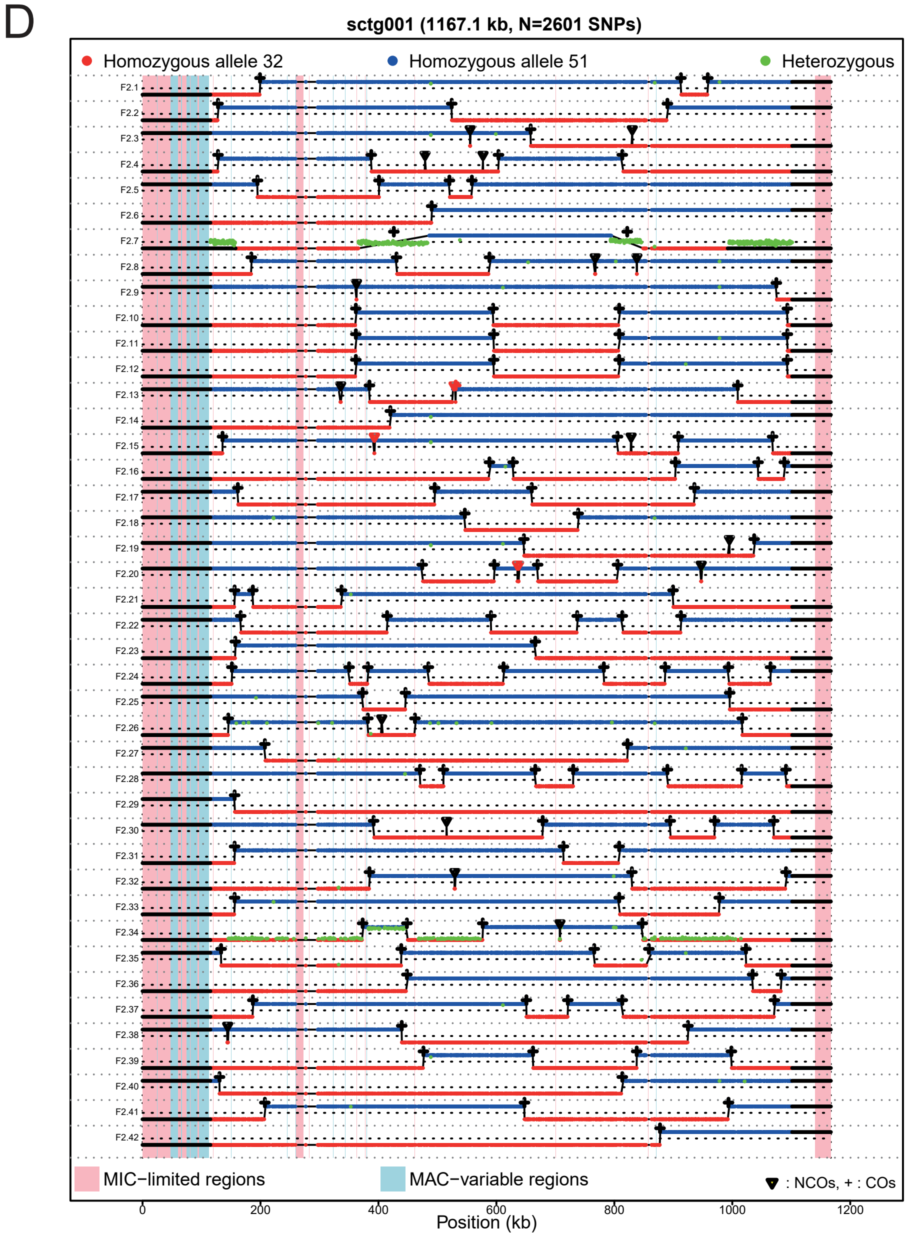


**SupFigure8 Analysis of meiotic recombination**

(A) Schematic representation of the genetic cross between P. tetraurelia strain 51 x strain 32, followed by autogamy yielding 100% homozygous F2 cells. The DNA from 42 F2 clones was sequenced and genotyped.
(B) Relationship between the G+C-content of MAC-destined regions and their recombination rate. The 161 points correspond to 158 supercontigs of the MIC genome assembly, and to 3 groups of 2 supercontigs that are linked based on the MAC genome assembly. The good correlation (R2 = 0.29, p= 2.9e-13) suggests that Paramecium is subject to GC-biased gene conversion, whereby GC/AT heteroduplex is repaired in favor of the GC allele [196]. (C) Crossover interference (1 –CoC) measured in 39 F2s (CoC: coefficient of coincidence). For each inter-interval distance, the CoC was calculated individually for all possible interval pairs genome-wide, and the average is plotted. CO interference decreases with increasing distance and reaches 0.5 for a distance of 22 kb. (D) Recombination map of MIC supercontig sctg001. The genotypes of the 42 F2's (F2.1 to F2.42) are shown along sctg001. Each dot corresponds to a SNP, colored according to its genotype (red: strain 32 allele, blue: strain 51 allele, green: heterozygous genotype). In a given F2, the frequency of reads corresponding to the strain 32 allele is indicated by the vertical position of the SNP (the horizontal dotted line corresponds to an allele frequency of 50%). This allows the distinction between truly heterozygous segments (as observed in F2.7) and artefacts of DNA contamination (as observed in F2.34). Crossovers are indicated by a cross, and non-crossover recombination events by a red triangle. Regions corresponding to MIC-limited or MAC-variable compartments are colored in pink and blue respectively. (E) Boxplots of recombination rate (cM/Mb) for a variety of eukaryotes. With a recombination rate of 420 cM/Mb, P. tetraurelia has the highest value reported to date. Note that the ciliate Tetrahymena thermophila, with a MIC genome of 200 Mb but only 5 large chromosomes [18], has a recombination rate of 9.9 cM/Mb [197], ruling out nuclear dimorphism per se as a determinant of high recombination rate.

SupFigure 9

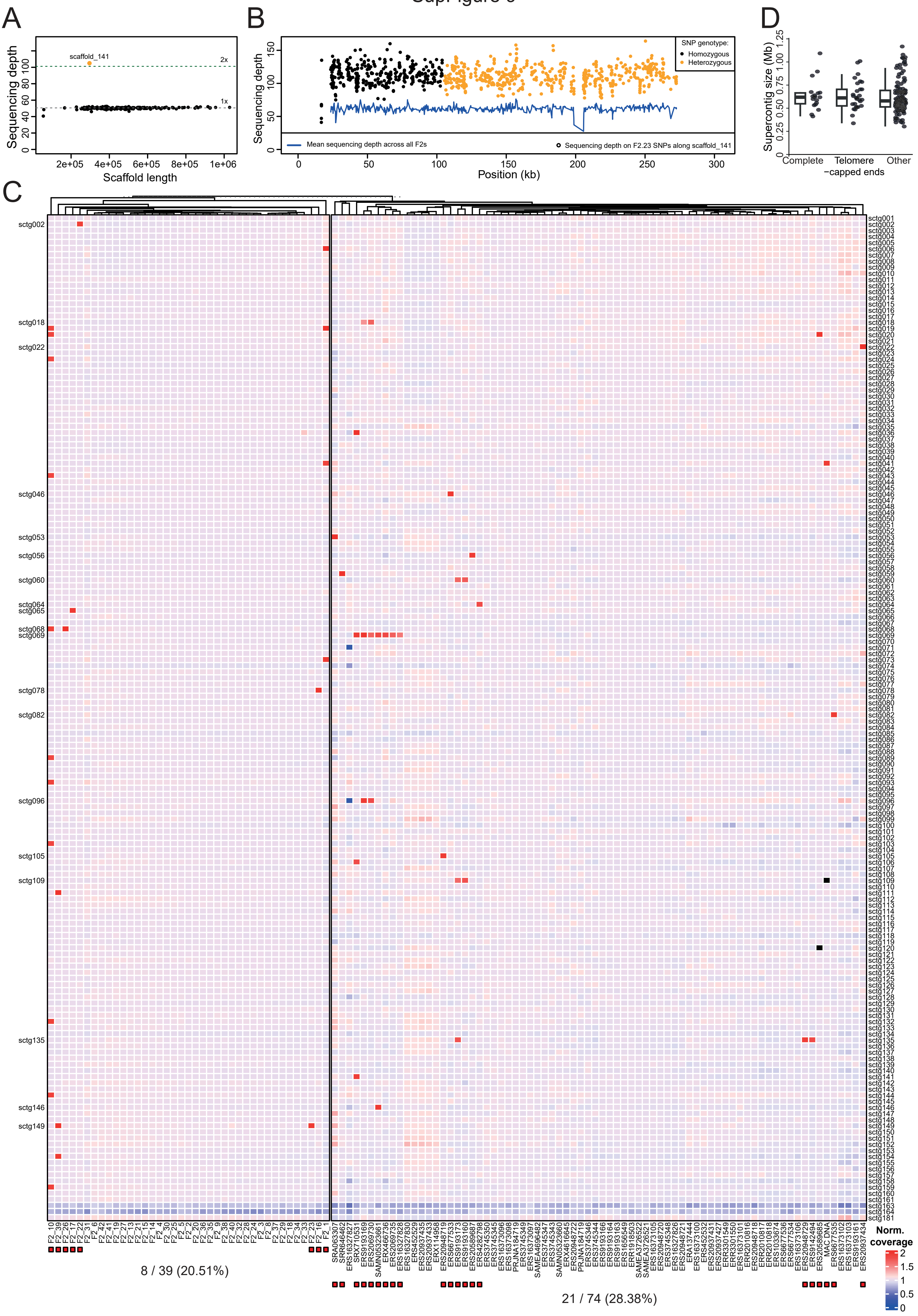
A
Sequencing depth
scaffold_141
2x
1x
Scaffold length
B
SNP genotype:
Homozygous
Heterozygous
Mean sequencing depth across all F2s
Sequencing depth on F2.23 SNPs along scaffold_141
Position (kb)
D
Supercontig size (Mb)
Complete
Telomere −capped ends
Other
C
8 / 39 (20.51%)
21 / 74 (28.38%)
Norm. coverage

**SupFigure 9 Aneuploidy** (A) Example of aneuploidy in an F2 individual (F2.23). Each dot corresponds to a MAC scaffold. In F2.23, all scaffolds have a sequencing depth of ~55x, except scaffold_141 with ~110x. Black dots represent scaffolds with an entirely homozygous genotype, while the yellow dot corresponds to scaffold_141, which contains heterozygous tracks.(B) Coverage of SNPs (sample F2.23 described in A) along scaffold_141; each dot corresponds to one SNP (black, homozygous; yellow, heterozygous). In F2.23, a large segment of scaffold_141 was genotyped as heterozygous. The sequencing depth is uniform along the entire scaffold_141 (including the heterozygous segments) and is twice as high as the average sequencing depth (blue line) in other F2's. All these observations indicate that scaffold_141 has a doubled ploidy (4n) in the F2.23 germline genome, whereas all other scaffolds have a normal ploidy. (C) Heatmap showing the level of aneuploidy of MIC EZL1 supercontigs in different samples (39 F2s of the genetic cross and 74 samples from 100% homozygous cell lines). The supercontigs are sorted by decreasing size (rows) and the samples (accession numbers beneath the heatmap) are ordered by hierarchical clustering of the normalized coverage values (columns). Normalized coverage values are coded in colors that range from blue (no coverage) to white (median coverage) to red (doubled coverage). Coverage greater than 2 is in black. If only a single supercontig is over-covered in one or more samples, that supercontig – which likely represents a complete chromosome (see Discussion) -- is indicated to the left of the heatmap. (D) Boxplots showing that these probably complete chromosomes have the same size distribution as the rest of the supercontigs. The length of supercontigs containing MIC-limited sequences were classified as 'Complete' chromosomes (defined in the previous panel), supercontigs capped by telomeric repeats at both ends (see Figure 2G), and the remaining supercontigs.

### SupFigure 10

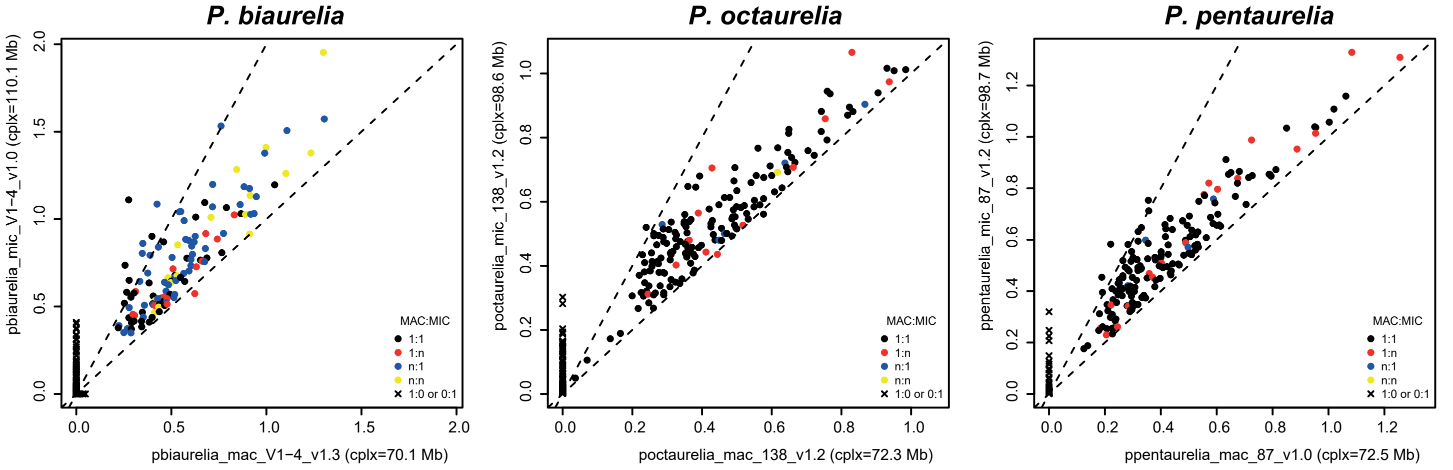


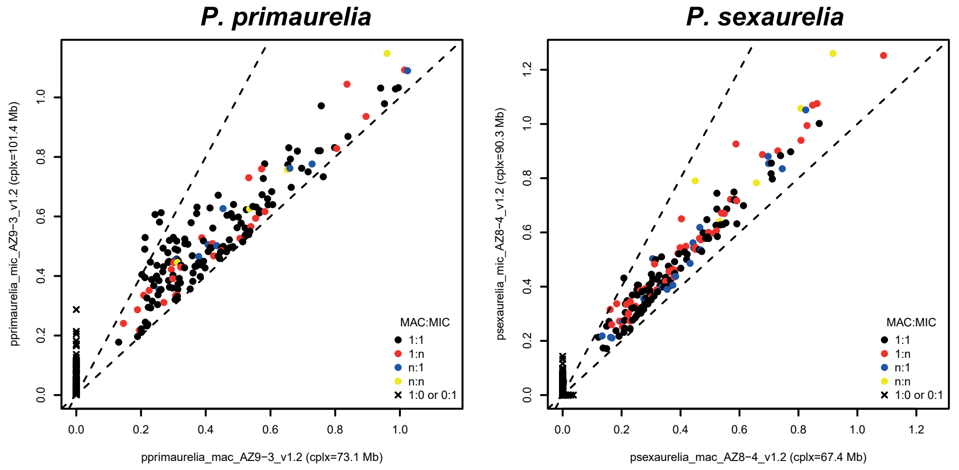


#### SupFigure10 Correspondence between MIC and MAC assembly scaffolds

Each dot shows MIC scaffold size as a function of the corresponding MAC scaffold size (cf. Methods for alignment and clustering details). Black dots indicate a 1:1 MIC/MAC relationship; red dots a 1:n relationship (one MAC for many MIC scaffolds); blue dots an n:1 relationship (many MAC for one MIC scaffold); yellow dots an n:n relationship (many MAC to many MIC scaffolds). Singlets (no MIC/MAC correspondence) are represented by black crosses. The whole of the data suggests a 1:1 MIC/MAC chromosome relationship across the aurelia species complex (cf. Figure 2).

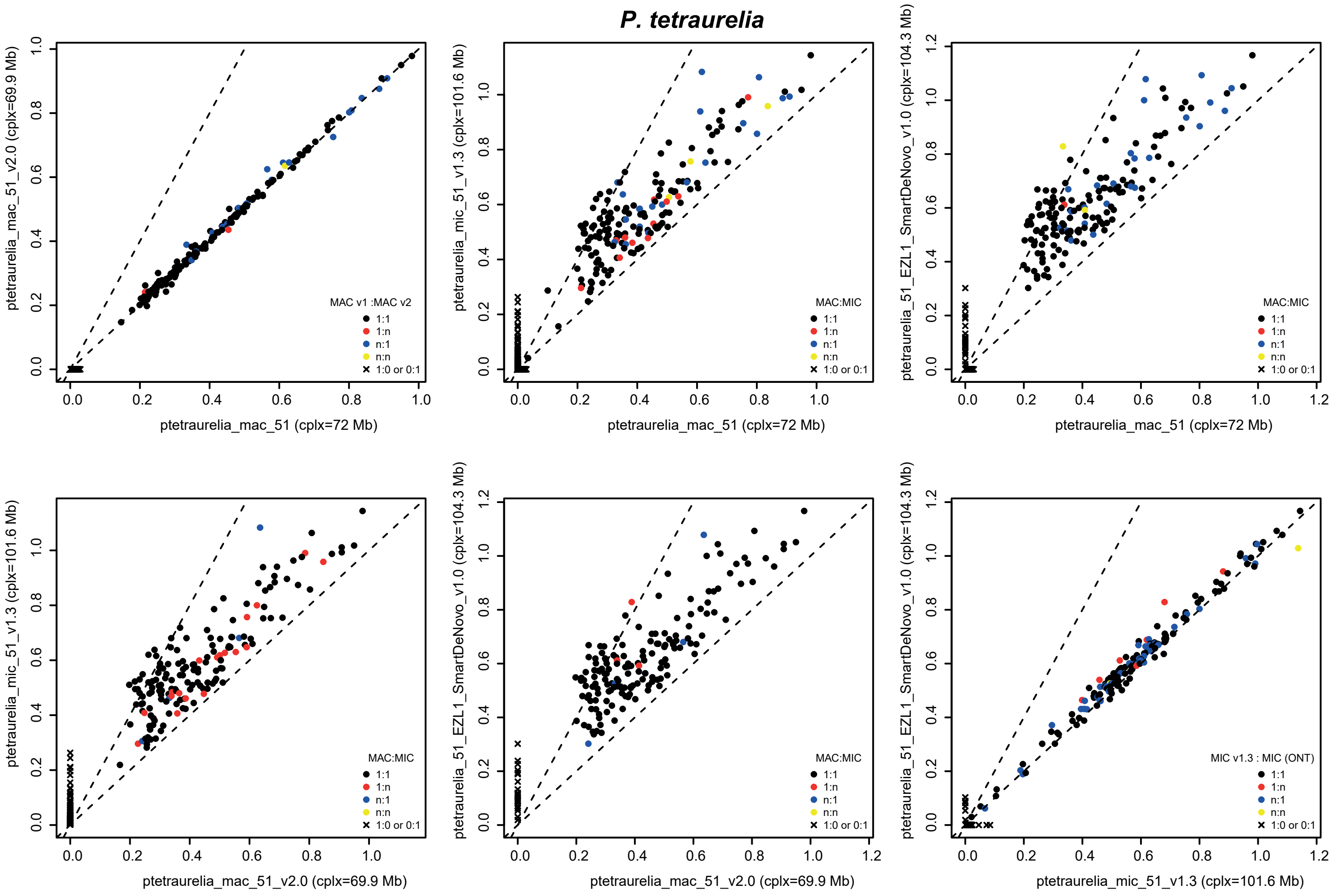

SupFigure 11

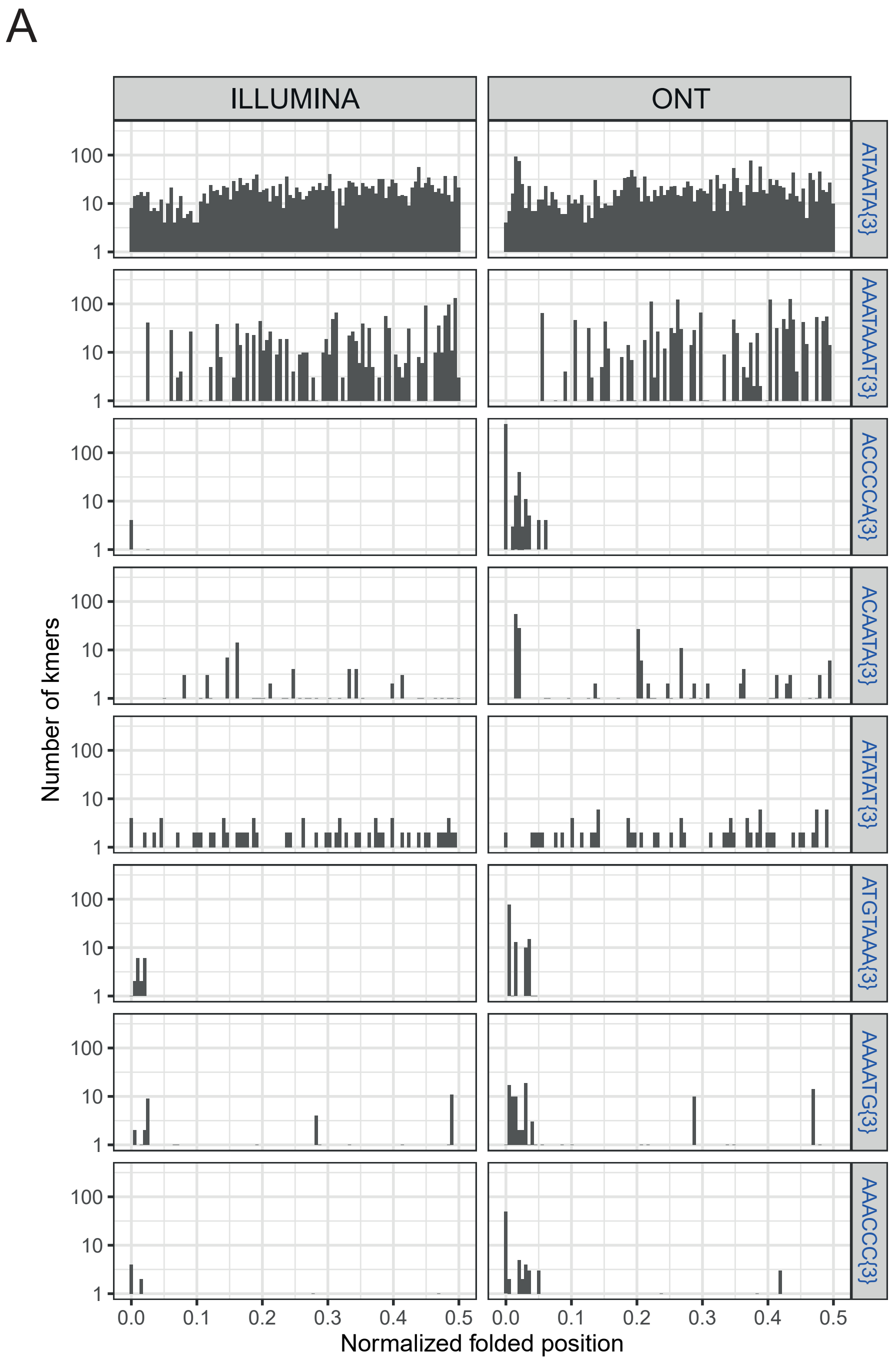


**SupFigure11 Identification of MIC telomere repeats.**

(A) k-mers of length 18, 21 and 24 nt, corresponding to 3 exact repeats ("triplets") of a 6, 7 or 8 nt motif, were counted in MIC Illumina paired-end reads that did not map properly to the corresponding MAC assembly (see Methods). The histograms show, for the 8 most abundant microsatellite triplets in P. tetraurelia, the distance of the triplets from contig ends (ranging from 0.0 for triplets at the contig end to 0.5 for triplets at the center of the contig). In addition to triplets containing only A and T, distributed evenly along the contigs, the top abundance triplets include 18-mers with 3 repeats of 5'-CCCCAA-3' and of 5'-CCCAAA-3' hexamers. These putative MIC telomere repeats are located at the ends of the EZL1-RNAI ONT supercontigs and the short-read MIC scaffolds. However, they are far less abundant in the Illumina MIC assembly.

SupFigure 12

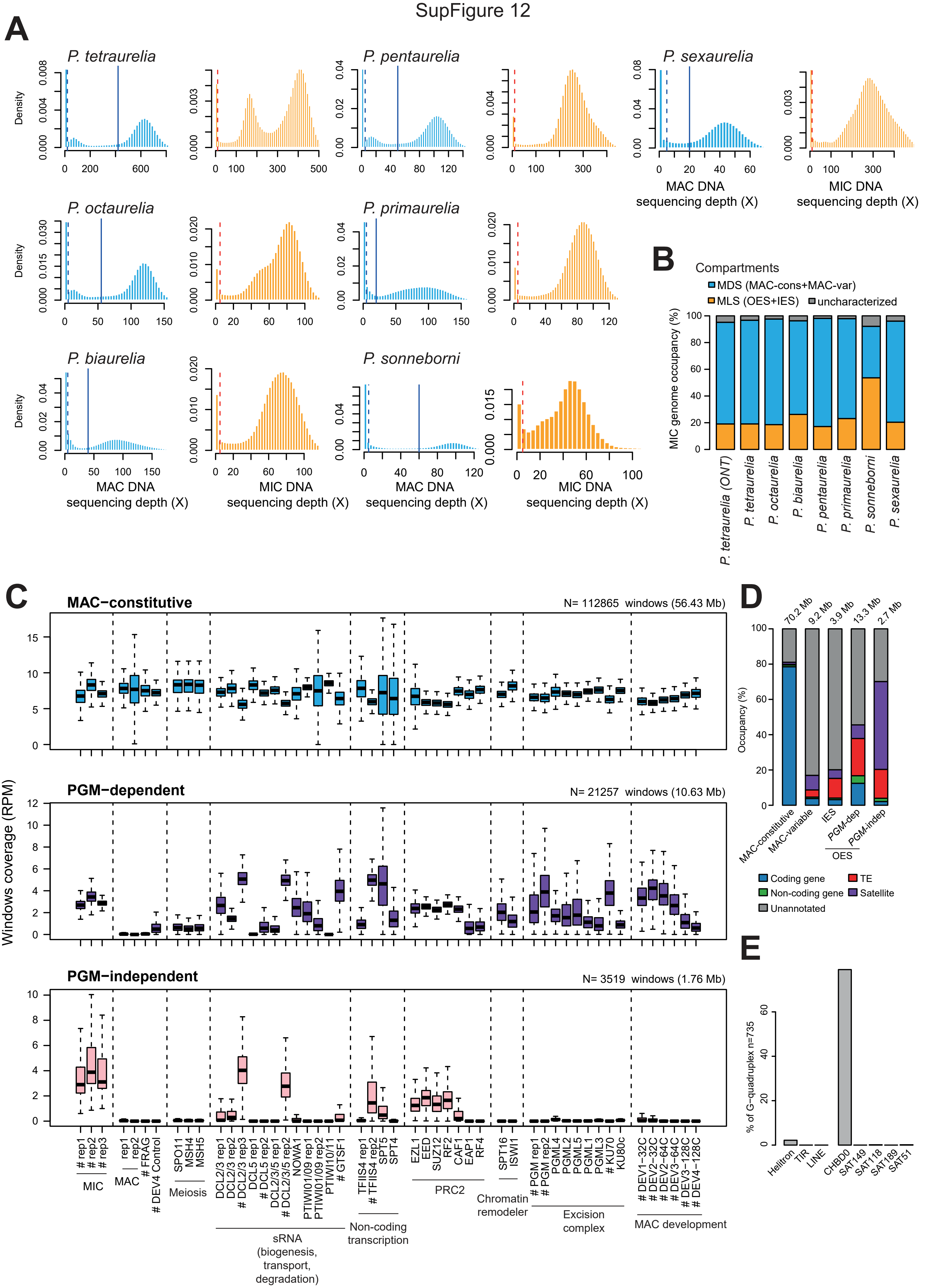

**SupFigure12 Definition and coverage of Paramecium MIC genomic compartments**

(A) Histograms of Illumina MIC (orange) and MAC (blue) sequencing depth for the MIC genomes (cf. Methods). The vertical blue line in the MAC coverage gives the lower threshold for the MAC-constitutive compartment. Coverage between the dotted vertical line and the solid vertical line was used to define the MAC-variable compartment. The remaining MIC regions with a MIC coverage above the value given by the dotted red line are defined as the MIC-limited sequence compartment. (B) Compartment occupancy (%) of MIC genomes by MDS (blue) and MLS (orange). MDS contains both the MAC-constitutive and the MAC-variable compartments. MLS contains both IES and OES (Other Eliminated Sequences). The uncharacterized regions of the MIC are indicated in grey. (C) For the P. tetraurelia EZL1 assembly, normalized DNA coverage (RPM) of non-overlapping 500 nt windows separated into 3 groups, as a function of their compartment (MAC-constitutive, Pgm-dependent, or Pgm-independent). DNA-Seq samples correspond to different types of nuclei or RNAi conditions. Samples in which nuclei were sorted are designated by "#". The boxplots summarize the presence of DNA in each compartment, for different DNA-Seq samples grouped by pathway (excision complex, scnRNA biogenesis and degradation, non-coding transcription, PRC2 components, and chromatin remodelers). These DNA-seq datasets are described in SupTable2. (D) Barplots show the proportion of each annotated sequence feature in the genome compartments (see Figure 3 and Figure 4). The size of each compartment is given at the top of the bars. (E) Proportion of predicted G-quadruplex loci present in the PGM-independent compartment (n=735; Figure 4A) that overlap annotated TEs (Helitron, TIR or LINE) or satellites (CHBD0, SAT149, SAT118, SAT189 and SAT51).

### SupFigure 13

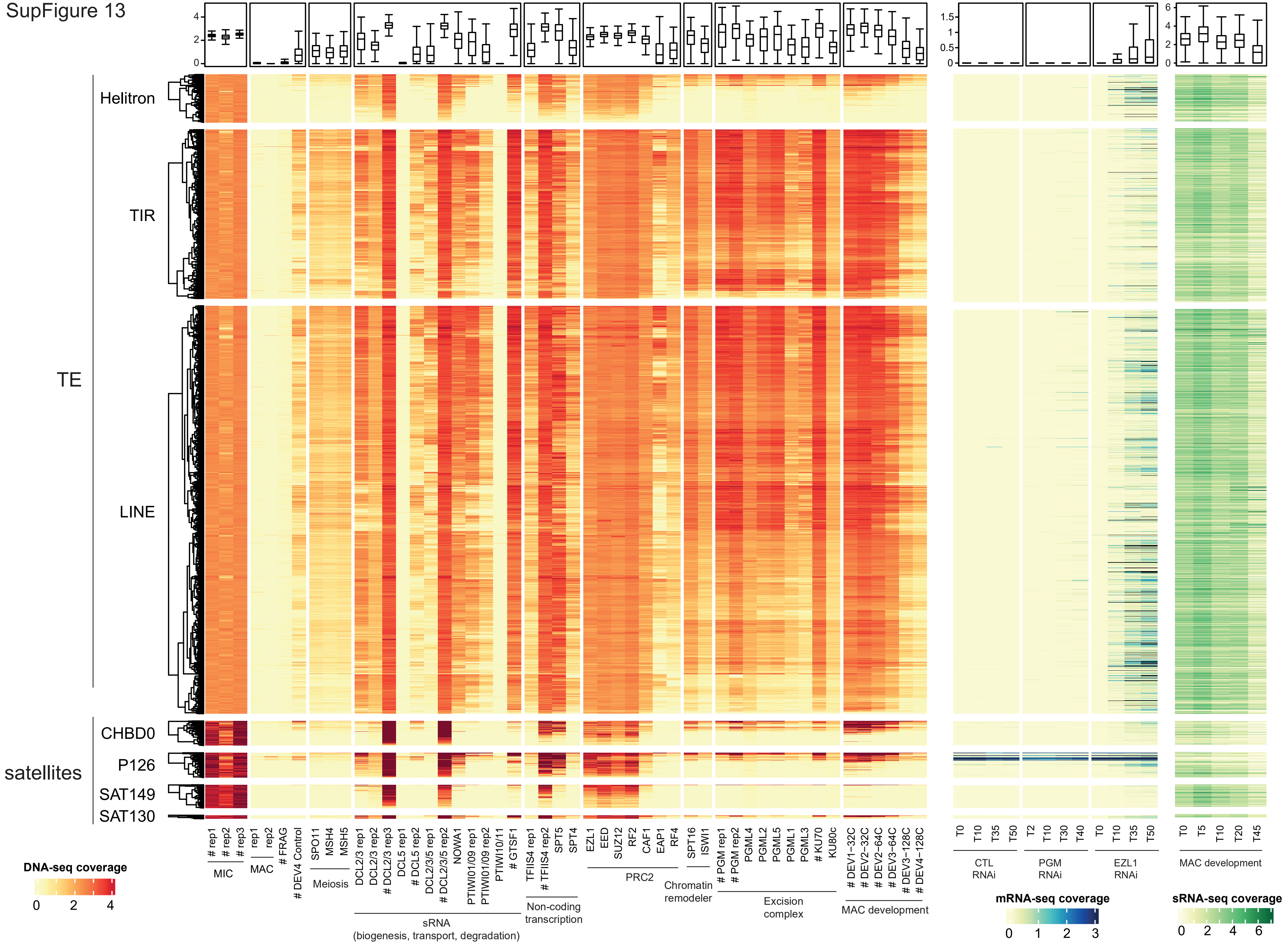

**SupFigure13  DNA and RNA coverage of P. tetraurelia Repeated Elements**

The heatmap shows the DNA-Seq coverage (orange palette), mRNA-Seq coverage (blue palette) and sRNA-Seq coverage (green palette) in RPKM (log2) for all REPET TE copies (Helitron, TIR and LINE superfamilies, minimum size 200 nt) and the four most abundant minisatellites (CHBD0, P126, SAT149 and SAT130, for regions of at least 200 nt). Samples used to calculate coverage are grouped by experiment and pathway (more details about samples in SupTable2). The DNA-seq sample labels preceded by “#” are those produced from sorted, as opposed to enriched, nuclei.

SupFigure 14

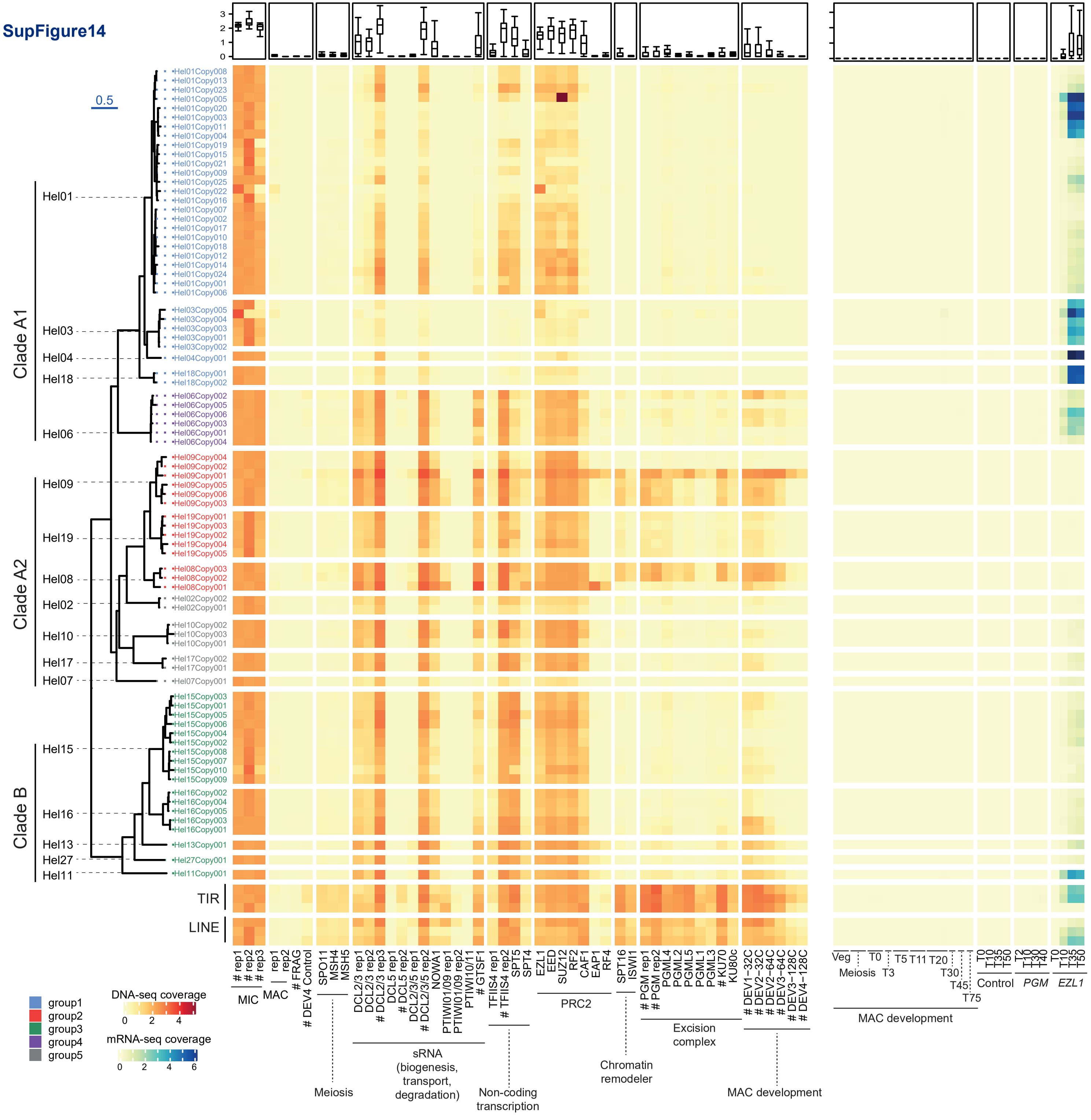

**SupFigure14 DNA and RNA coverage of the longest P. tetraurelia Helitron ORFs**

The heatmap shows the DNA-Seq coverage (orange palette) grouped by development stages or PDE pathway and mRNA-Seq coverage (blue palette) in RPKM (log2) for all P. tetraurelia 9-10 kb Helitron ORFs and the three longest TIR and LINE copies (in the same order as in Figure 5). Samples used to calculate coverage are described in SupTable2. The DNA-seq samples preceded by “#” are those produced from sorted, as opposed to enriched, nuclei.

Text

SupFigure 15

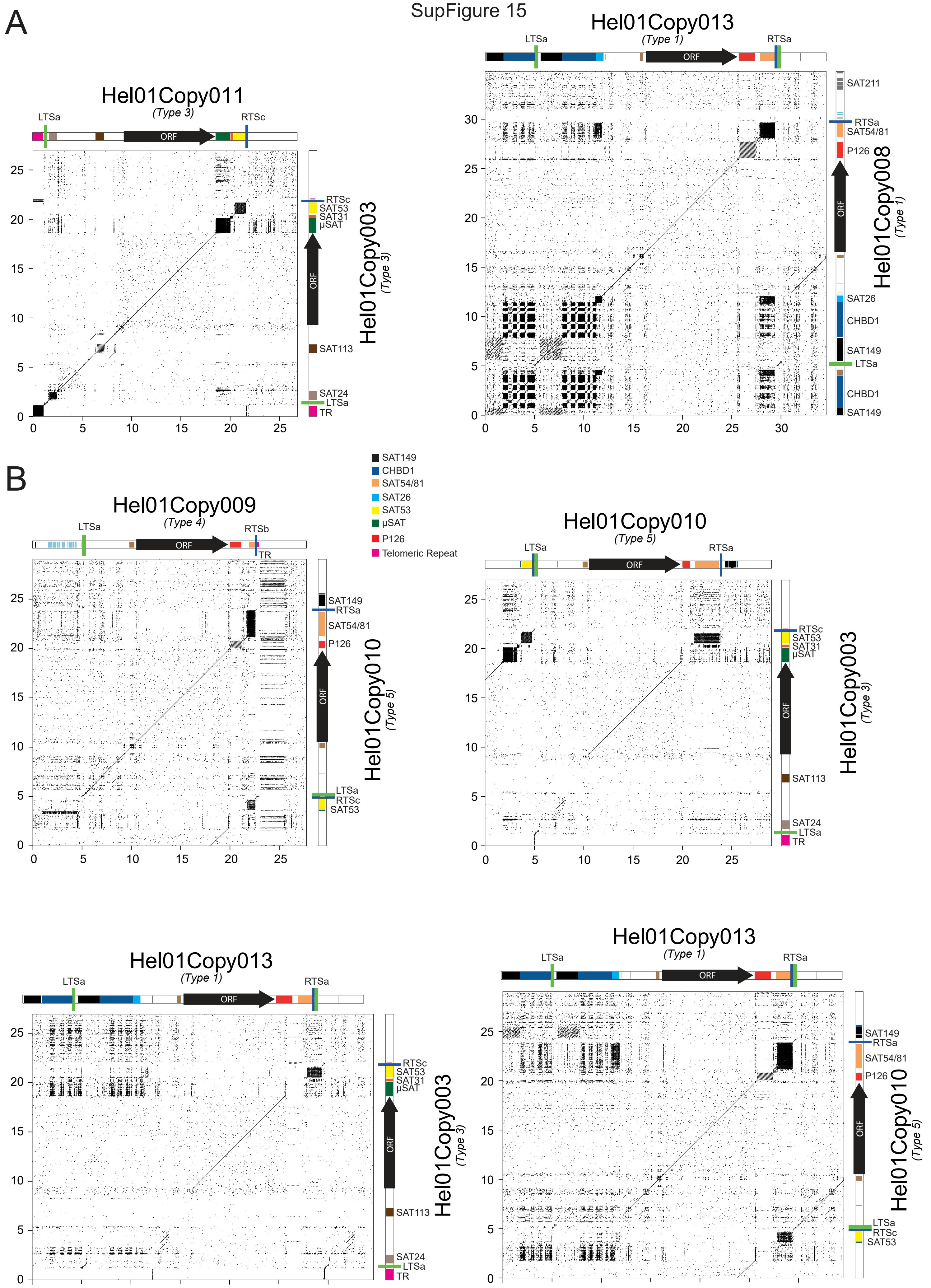

**SupFigure15 Nucleotide alignment of different Hel01 copies**

(A). Dotplots showing the similarity between copies of the same Hel01 Type. On the left, Hel01Copy011 and Hel01Copy003 from Type 3. On the right, Hel01Copy013 and Hel01Copy008 from Type1. (B). Dotplots showing similarity between copies of different Hel01 Types. Each dot corresponds to at least 70% identity over a 28 nt window. Annotations (LTS, RTS, ORF, satellites) for each copy are drawn above and to the right of each dotplot. The legend presents the different annotated satellites. Microsatellite (µSAT) consists of (TTA/TTG)n. The scale in kb is given on the left and bottom axes of each dotplot.

### SupFigure 16

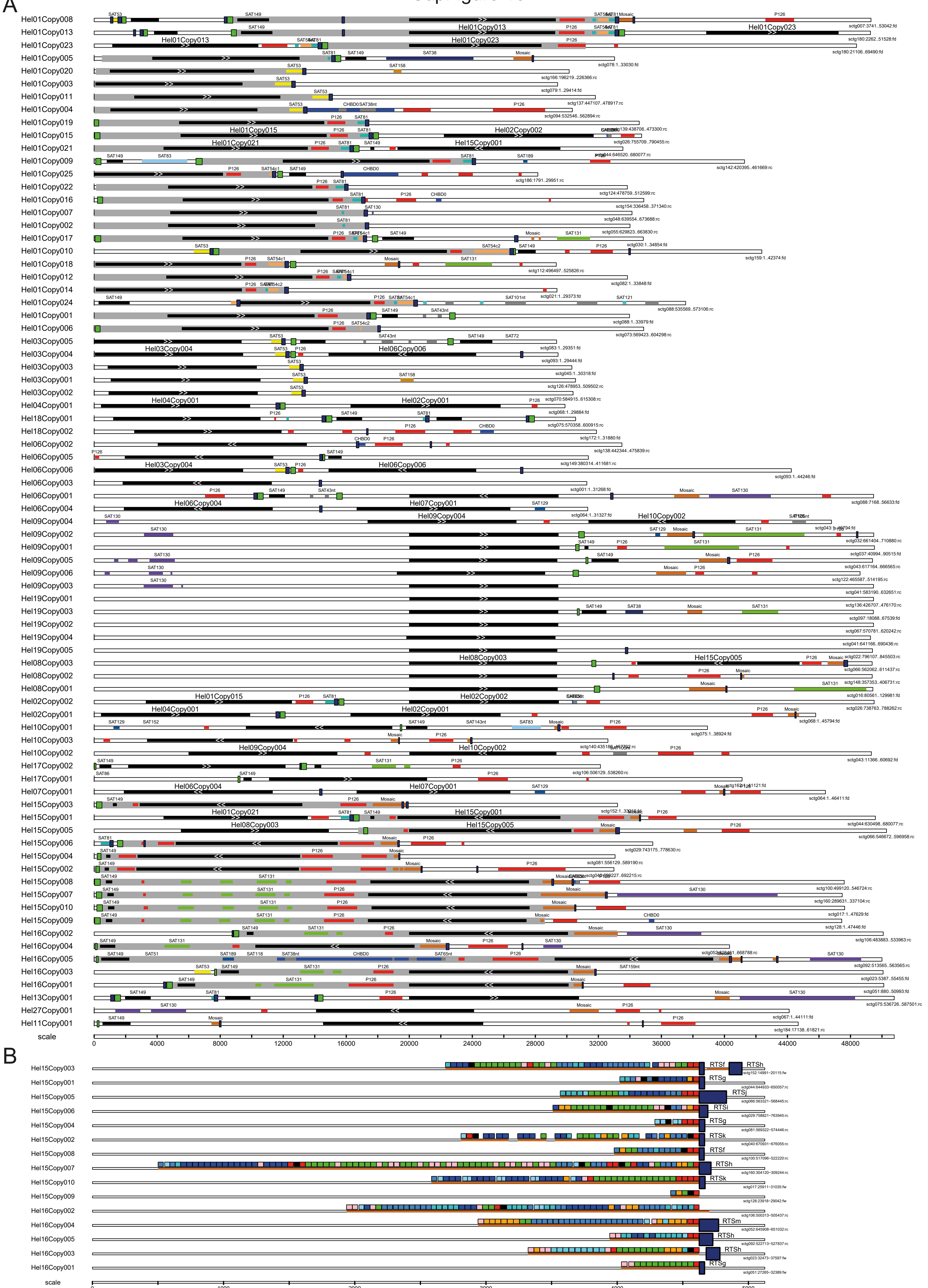

**SupFigure16 Annotations of helitron copies with full-length ORFs**

(A) Scale drawings showing all supercontig regions with Helitrons containing full-length ORFs. The closest extremity of the supercontig is shown on the left and the genomic position is indicated on the right with forward (fd) and reverse complement (rc) orientation. The RepHel ORF (black box) is oriented with ”>>”, LTS (lime green box) and RTS (navy blue box) sequences delimiting the putative Helitron element (gray rectangles). Satellite DNA is colored as in SupData3 and SupData4. Nucleotide scale is indicated at the bottom. (B) Zoom of Hel15/ Hel16 copies to show mosaic satellite repeat structure and the presence or absence of a putative RTS (navy blue. Each “color” of the mosaic satellite represents a distinct ~44 nt repeat unit.

SupFigure 17

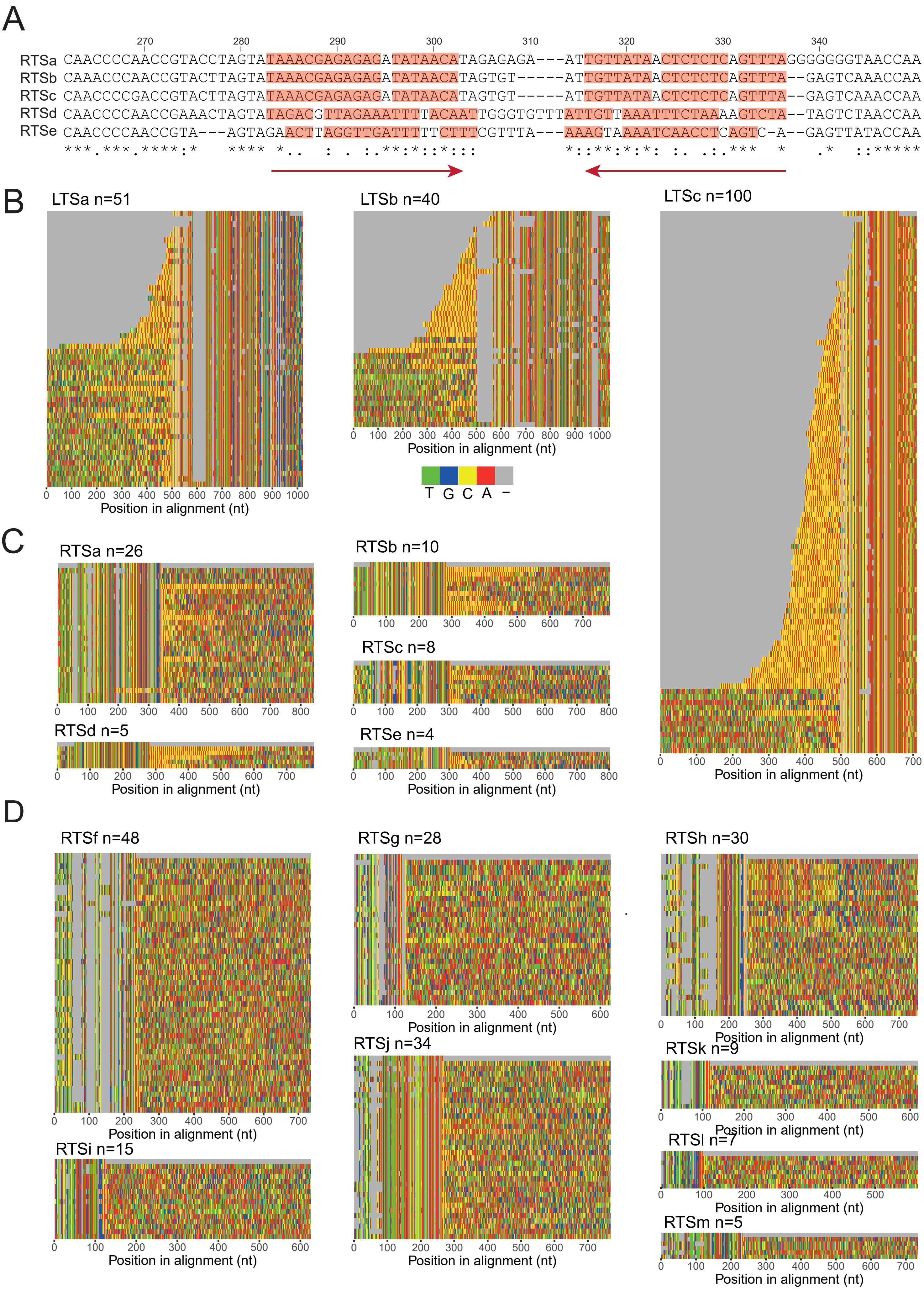

**SupFigure17 Structural characterisation of Helitron ends and insertion sites**

(A) Alignment of different Right Terminal Sequence (RTS) consensus sequences for Paramecium Helitrons. The arrows beneath the alignment indicate the hairpin (positions 280-340), typically found at the 3’ end. These RTS end with CCAA (positions 347-350). (B) Alignment of the5’ and 3’ flanking sequences of 500 and 100 nucleotides, respectively, for left terminal sequence (LTS) loci. As in Figure 2E, color indicates the type of nucleotide. Note the telomeric repeats (red/yellow) before most copies of LTSa, b or c. (C) Alignment of RTSa to RTSe copies associated with Clade A1 Helitrons, with 5' and 3' flanking sequences of 100 and 500 nucleotides, respectively. Note telomeric repeats after most RTS copies. (D) Alignment of RTSf to RTSm copies associated with Clade B Helitrons, with 3’ 500-nt flanking sequences.

SupFigure 18

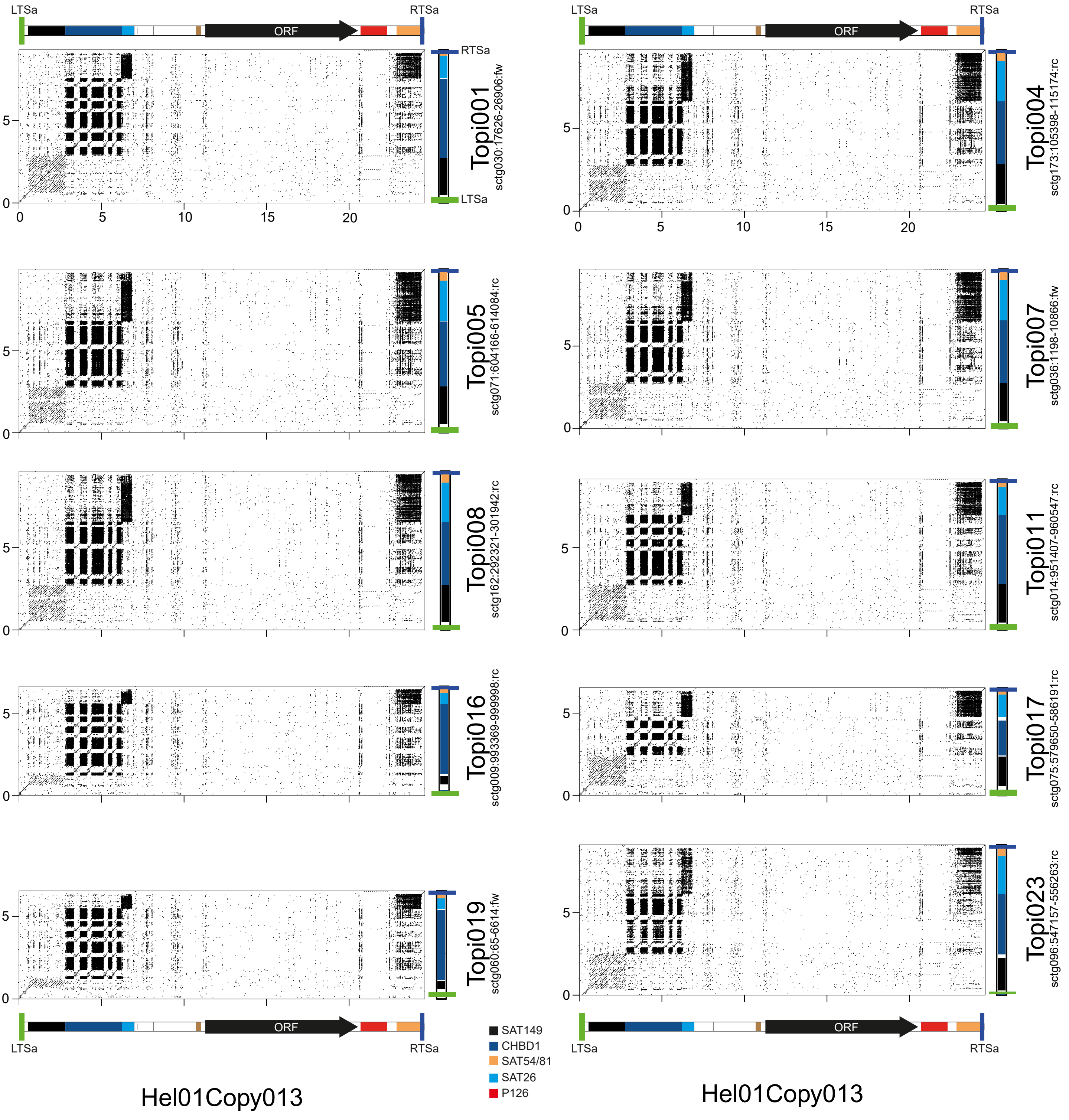


**SupFigure18 Structure of Hel01 ORF-less copies**

Dotplots as in SupFigure15, illustrating 10 examples of Hel01 copies with internal deletions that encompass the ORF (y-axis, labeled with “Topi” prefix) with complete Hel01Copy013 (x-axis).

SupFigure 19

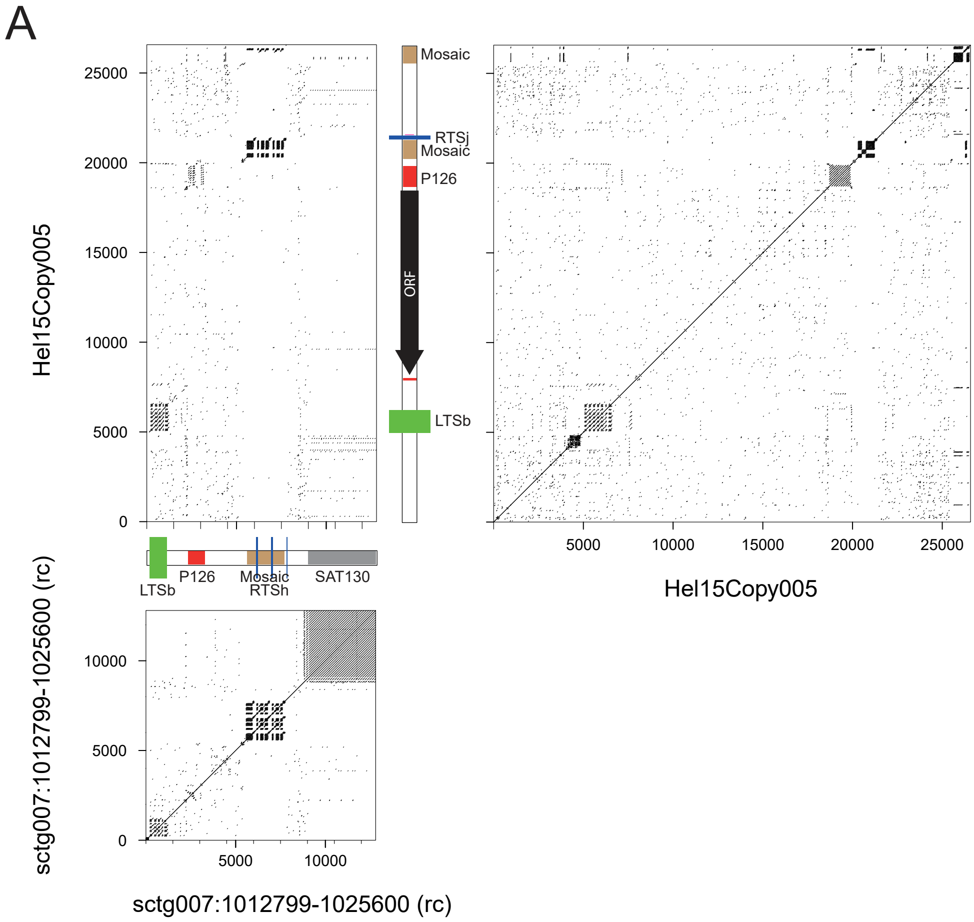


**SupFigure19 Structure of Hel15/Hel16 ORF-less copies**

As in SupFigures 17-18. (A) Comparison of Hel15Copy005 with itself, an ORF-less locus (right end of sctg007) and of the ORF-less locus with itself

SupFigure 20

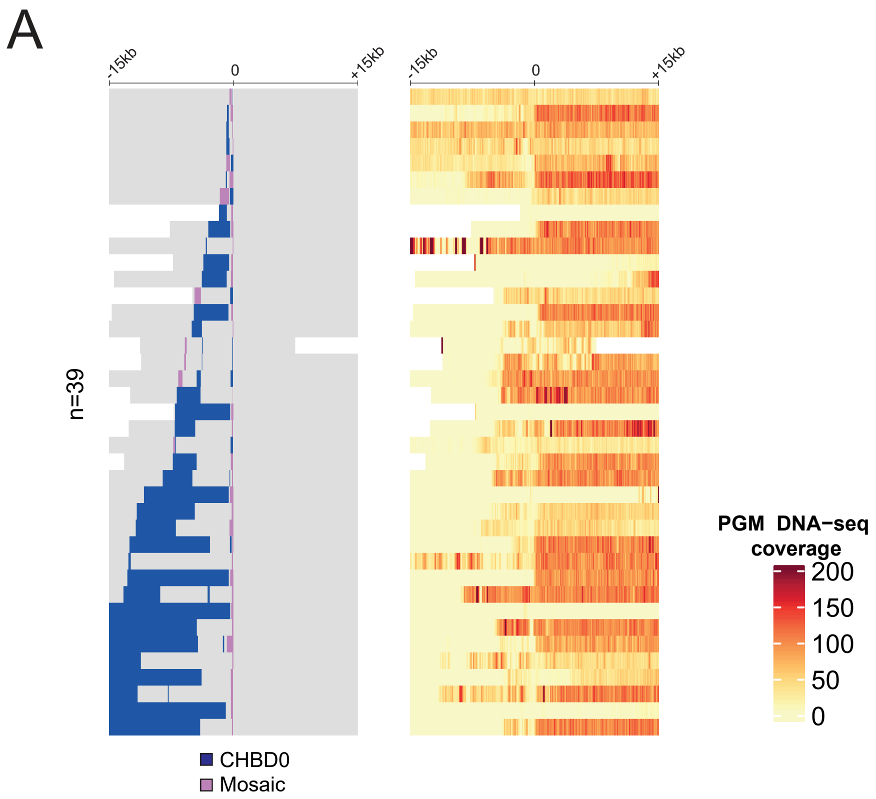


**SupFigure20 Loci that contain both CHBD0 and Mosaic satellite**

As in Figure7 C-D, heatmap for 39 loci that contain CHBD0 (blue) and Mosaic (orchid) separated by less than 15 kb centered in a 30 kb window.
